# Protein Interactome of the Cancerous Inhibitor of Protein Phosphatase 2A (CIP2A) in Th17 Cells

**DOI:** 10.1101/809459

**Authors:** Mohd Moin Khan, Tommi Välikangas, Meraj Hasan Khan, Robert Moulder, Ubaid Ullah, Santosh Dilip Bhosale, Elina Komsi, Umar Butt, Xi Qiao, Jukka Westermarck, Laura L Elo, Riitta Lahesmaa

**Affiliations:** Turku Bioscience Centre, University of Turku and Åbo Akademi University, FI-20520 Turku, Finland; Turku Doctoral Programme of Molecular Medicine (TuDMM), Medical Faculty, University of Turku, Turku, Finland; Doctoral Programme in Mathematics and Computer Sciences (MATTI), University of Turku, Turku, Finland; Institute of Biomedicine, University of Turku, Turku, Finland

**Author notes:** These authors contributed equally to this work (Mohd Moin Khan, Tommi Välikangas, Meraj Hasan Khan). **Correspondence** Professor Riitta Lahesmaa, Turku Bioscience Centre, Tykistökatu 6A (7th floor), Turku 20520, FINLAND.

**Keywords:** CIP2A, Interactome, Mass Spectrometry (MS), Selected Reaction Monitoring (SRM) targeted mass spectrometry, confocal microscopy, Pathway analysis

## Abstract

Cancerous inhibitor of protein phosphatase 2A (CIP2A) is involved in immune response, cancer progression, and in Alzheimer’s disease. However, an understanding of the mechanistic basis of its function in this wide spectrum of physiological and pathological processes is limited due to its poorly characterized interaction networks. Here we present the first systematic characterization of the CIP2A interactome by affinity-purification mass spectrometry combined with validation by selected reaction monitoring targeted mass spectrometry (SRM-MS) analysis in Th17 cells. In addition to the known regulatory subunit of protein phosphatase PP2A, the catalytic subunit of protein PP2A was found to be interacting with CIP2A. Furthermore, the regulatory (PPP1R18, and PPP1R12A) and catalytic (PPP1CA) subunits of phosphatase PP1 were identified among the top novel CIP2A interactors. Evaluation of the ontologies associated with the proteins in this interactome revealed that they were linked with RNA metabolic processing and splicing, protein traffic, cytoskeleton regulation and ubiquitin-mediated protein degradation processes. Taken together, this network of protein-protein interactions will be important for understanding and further exploring the biological processes and mechanisms regulated by CIP2A both in physiological and pathological conditions.

**Highlights:**

▪ The first characterisation of the CIP2A interactome in Th17 cells.
▪ Key interactions were validated by targeted SRM-MS proteomics, western blot and confocal microscopy.
▪ Pathway analysis of the interactome revealed interrelationships with proteins across a broad range of processes, in particular associated with mRNA processing.

## INTRODUCTION

PP2A is a recognized tumor suppressor that imparts crucial functions in regulating cell proliferation, differentiation, and apoptosis [1] – [7]. Accordingly, down regulation of PP2A has been observed as a feature in many cancers. Cancerous Inhibitor of PP2A (CIP2A) was first characterized as an endogenous inhibitor of protein phosphatase 2A (PP2A) in cancer cells [8]. Previous studies have shown that many of the known actions of CIP2A are mediated through inhibition of PP2A [9], [10]. Overexpression of CIP2A has been associated with the poor prognosis in several human malignancies, and CIP2A has thus been seen as a potential therapeutic target for cancer therapy [8], [11], [12]. CIP2A supports the activity of many critical onco-proteins (Akt, MYC, E2F1) and promotes malignancy in most cancer types via PP2A inhibition [8]. Structurally, CIP2A forms a homodimer and this dimerization promotes its interaction with the PP2A regulatory subunits B56α and B56γ [13]. Interestingly, inhibition of either CIP2A dimerization or B56α/γ expression destabilizes CIP2A in cancer cells [13]. In Alzheimer’s disease (AD), enhanced expression of CIP2A in AD brains and neurons results in PP2A inhibition and tau protein hyperphosphorylation, which leads to abnormal tau localization with memory deficits and other pathological conditions [14].

Th17 cells are found at mucosal surfaces where they regulate tissue homeostasis [15] – [18]. Dysregulation of Th17 cell differentiation, however, has been associated with a number of autoimmune inflammatory diseases [19], [20]. Following our earlier report on the involvement of CIP2A in T cell activation [21], we subsequently observed that CIP2A negatively regulates T helper (Th)17 cell differentiation, whereby CIP2A depletion resulted in enhanced IL17 expression both in human and mouse Th17 cells (Khan M.M et al., submitted).Thus, CIP2A could have important implications in the differentiation and regulation of Th17 cells. Although CIP2A has been shown to regulate a range of pathological and physiological conditions, there has not been any systematic analysis of the full range of signaling pathways and processes regulated by CIP2A.

In the current study, we used immunoprecipitation followed by label free quantitative proteomics to gain insight into the proteins that are associated with CIP2A. We present the first interactome of CIP2A, with validation of interactors by selected reaction monitoring (SRM) targeted mass spectrometry as well as by Western blotting and confocal microscopy. In addition to its known interaction with the regulatory subunit A and B of phosphatase PP2A, our data demonstrated CIP2A interaction with the PP2A catalytic subunits namely PPP2CA and PPP2CB. Furthermore, both the regulatory (PPP1R18, and PPP1R12A) and catalytic (PPP1CA) subunits of phosphatase PP1 were identified among the top interactors of CIP2A. This data further implicates the involvement of CIP2A in a number of biological processes that it has previously not been associated to, and provides a basis for comprehensive understanding of CIP2A for translational studies targeting CIP2A in human diseases.

## RESULTS

### Mass spectrometry analysis identify novel CIP2A binding partners

To identify the proteins interacting with CIP2A and thereby facilitate the understanding of its functions in human Th17 cells, immunoprecipitation (IP) was performed after 72 h of polarization of naïve T cells towards the Th17 lineage **(Fig. 1A; S1A-C).** The cell model for the study was rationalized by our recent findings indicating that CIP2A is involved in T-cell activation [21] and its particular importance for Th17 cell differentiation (Khan MM et al., submitted). Two CIP2A specific antibodies (Ab1 and Ab2) recognizing distinct regions of CIP2A were used in two biological replicates, together with their respective IgGs (IgG1 and IgG2) as control antibodies. Ab1 was a monoclonal antibody targeting the N-terminal structured region of the protein (residues surrounding Val 342) [13], while Ab2 was of polyclonal origin targeting unstructured C-terminus [22] (Fig. 1B).

**Figure 1.**
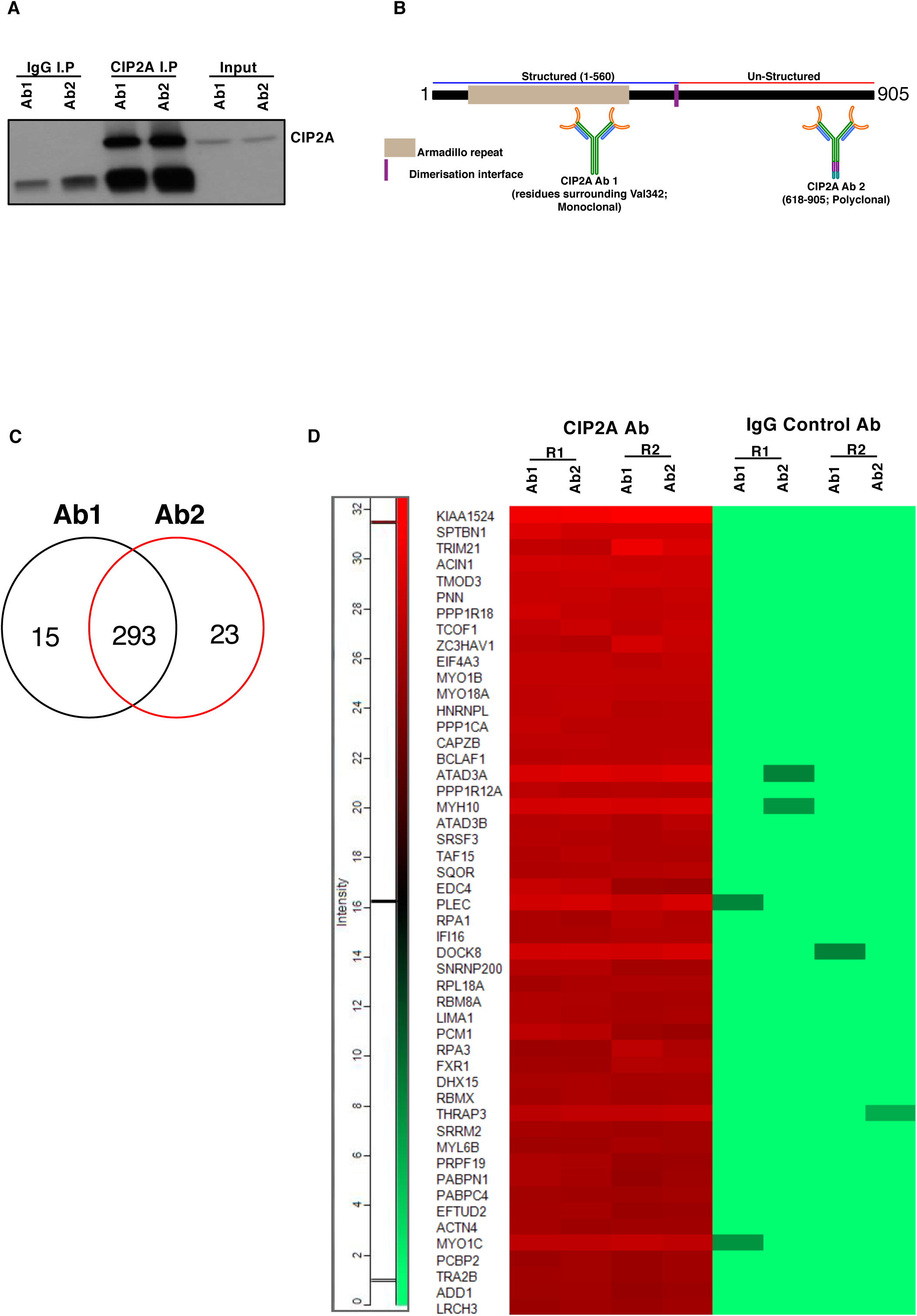
CIP2A interactome top protein interaction in Th17 cells. **(A)** Western Blot (WB) analysis of CIP2A immunoprecipitated (IP) with two antibodies (Ab1 and Ab2) specific to different regions of CIP2A in 72h polarised Th17 cells. For each CIP2A targeting antibody, an IgG control antibody (IgG1 and IgG2) was used. The figure is a representative WB of two experiments. Input, IgG control IP and CIP2A IP reactions are shown. **(B)** Schematic diagram of the human CIP2A protein structure. Regions specific to the antibodies used in the pulldown are shown. Ab1: against residues surrounding valine 343 (monoclonal), Ab2: 618-905 peptide (polyclonal). Protein domain borders are based on previous publication (Wang et al., 2017) in which the Armadillo repeat, and the dimerization interface are shown in the CIP2A structured domain studied by Wang et al., 2017. **(C)** A Venn diagram showing the number of common and unique CIP2A interacting proteins identified with two distinct antibodies targeting CIP2A based on the mass spectrometry (ms) analysis. For details, please see the method section. **(D)** Heatmap representation of the top fifty CIP2A interactors identified with both antibodies. R1 and R2 represent data from two biological replicates. Normalized spectral intensities are plotted as a heatmap to display protein abundance in respective IP reactions.

The availability of highly specific antibodies against CIP2A facilitated the IP of endogenous CIP2A without the requirement for overexpression. After preparation for proteomic analysis, the digested samples were analyzed by LC-MS/MS to identify the CIP2A associated proteins. Analysis of the data revealed 680 proteins meeting the criteria of identification of two or more unique peptides with a confidence threshold of 99%. In order to distinguish non-specific interactions and common contaminants, statistical analysis of the LFQ intensity data was made using the Significance Analysis of *INTeractome* (*SAINT*) algorithm [23]. Using a threshold for the calculated SAINT probability scores of greater than or equal to 0.95, and exclusion of proteins represented in the contaminant’s repository [24] with a frequency of >60%, the list of candidate interacting proteins was reduced to 331 (**Supplementary Table 1A-C**) with more than 90% overlap in common proteins identified with both antibodies specific to CIP2A **(Fig. 1C; S1D; Supplementary Table 1A-C)**. To exemplify the interactions between CIP2A and its interaction partners, the top fifty (50) common proteins detected with both antibodies based on the detected signal intensity is plotted in the form of heatmap **(Fig. 1D)**.

### Distribution of CIP2A interactome between the nucleus and cytoplasm

Analysis of the cellular location of CIP2A in epithelial cells has indicated that it is mainly expressed in the cytoplasm [8], [22]. Further data indicate some representation in the nuclear fraction of epithelial cancer cells [25]. To evaluate the distribution of CIP2A in Th17 cells, confocal microscopy measurements were conducted in this study. For these measurements, 4,6-diamidino-2-phenylindole (DAPI), and Phalloidin were used to stain the nuclear and cytoplasmic fractions, respectively (**Fig 2A**). Statistical analysis of the confocal microscopy images revealed CIP2A is predominantly located in the cytoplasmic compartment of Th17 cells, in addition to some nuclear expression, (**Fig 2B**). Confirmatory measurements were provided with Western Blot (WB) analysis of the cytoplasmic and nuclear fractionations of 72h polarised Th17 cells. For these experiments, vimentin and tubulin were used as control markers for the nuclear and cytoplasmic fractions, respectively (**Fig 2C; S2A**). Whilst the observed expression of CIP2A was greater in the cytoplasmic region, CIP2A was also detected in the nuclear region of 72h differentiated Th17 cells (**Fig 2C; S2A**).

**Figure 2.**
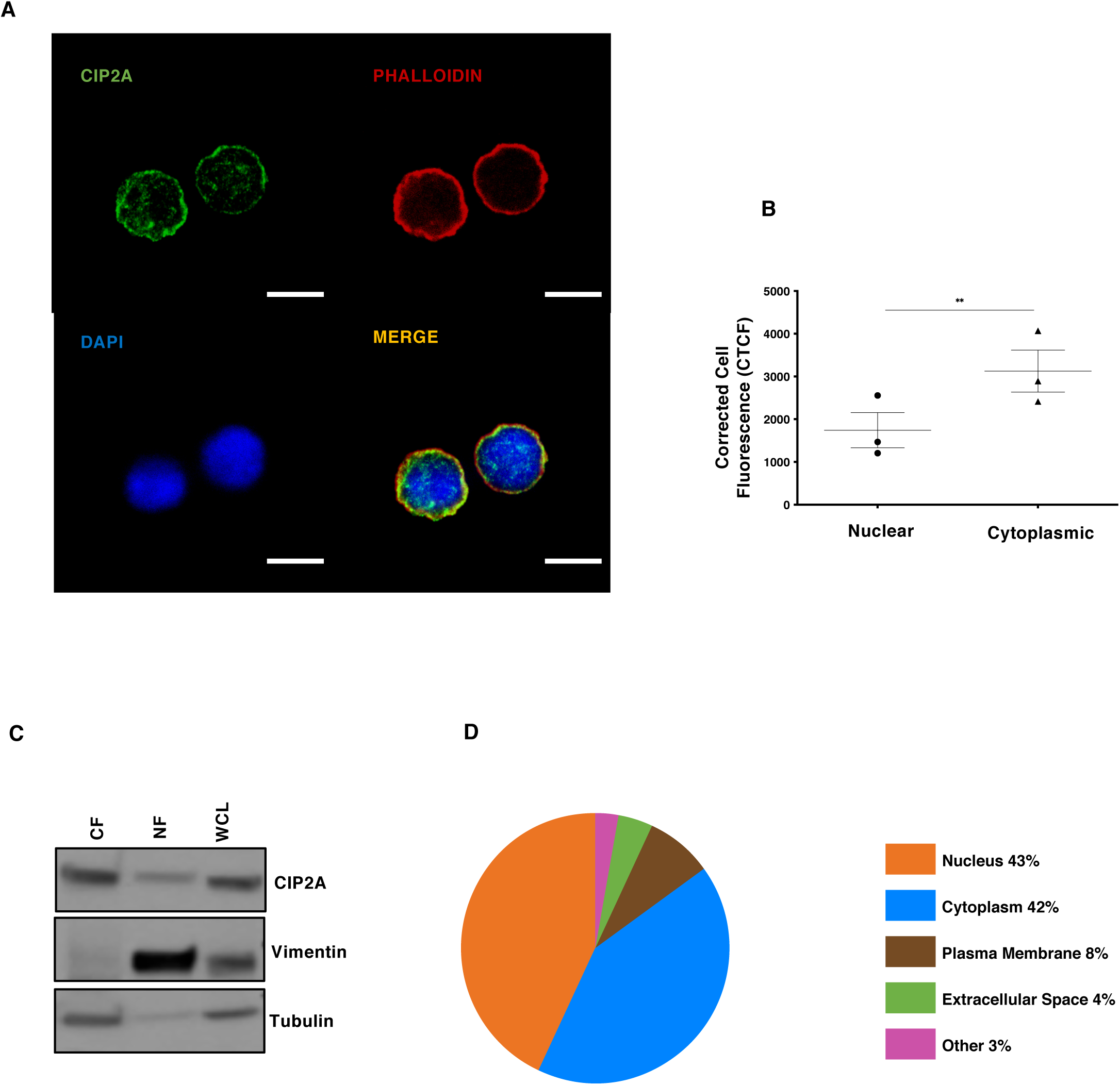
Cellular distribution of CIP2A and the proportions of CIP2A interactome in Th17 cells. **(A)** Immunostaining for the study of CIP2A localisation in 72h polarised Th17 cells. A representative image of three replicates are shown. Cells were stained for endogenous CIP2A (green). Phalloidin (red) and DAPI (blue) were used to stain cytoplasmic and nuclear regions, respectively. Scale bar is 7μm. **(B)** Statistical analyses of CIP2A expression for cellular localization studies from three replicates using GraphPad Prism version 8.0d for Mac OS X (GraphPad Software). Statistical analysis was made by a two-tailed paired Student’s T-test, where p-value ** represent <0.01. The error bars represent standard error of the mean. **(C)** Representative WB analysis of CIP2A localisation in the cytoplasmic and nuclear fractions of 72h polarised Th17 cells. Vimentin and Tubulin were used as controls for the nuclear and cytoplasmic fractions, respectively. WCL: Whole Cell lysate, NF: Nuclear Fraction, and CF: Cytoplasmic Fraction. **(D)** Enriched cellular locations of the proteins in the CIP2A interactome presented in the form of pie chart. The enrichment analysis was performed using the Ingenuity pathway analysis (IPA; Qiagen).

Ingenuity Pathways Analysis (IPA; Qiagen) was used to map cellular locations of the proteins interacting with CIP2A. The analysis revealed that the cytoplasm (42%) and nucleus (43%) constituted the most frequent cellular locations of the associated proteins, whilst the plasma membrane (8%), extracellular space (4%), and undefined (other, 3%) accounted for the remainder (**Fig 2D**). The observed distribution of the proteins detected in the CIP2A interactome is consistent with its association in these compartments in Th17 cells.

### Cellular processes regulated by CIP2A interactome

To gain an overview of the biological processes associated with CIP2A, we analyzed the CIP2A interactome data with several Gene Ontology and interaction bioinformatics tools. To collate the known interactions between the proteins, a network was constructed using the STRING database [26]. The resulting network was visualized with Cytoscape [27] revealing “RNA metabolism or splicing” as the process most frequently linked with the associated proteins (**Fig. 3A; Supplementary Table 2 and 3**). The top 20 proteins associated with RNA metabolic process, RNA splicing via transesterification and RNA splicing via spliceosome are represented in the form of heatmaps (**Fig. S3A-C**). In agreement with these results, a recent phosphoproteome screen also identified RNA splicing as one of the most significantly CIP2A-regulated processes [28].

**Figure 3.**
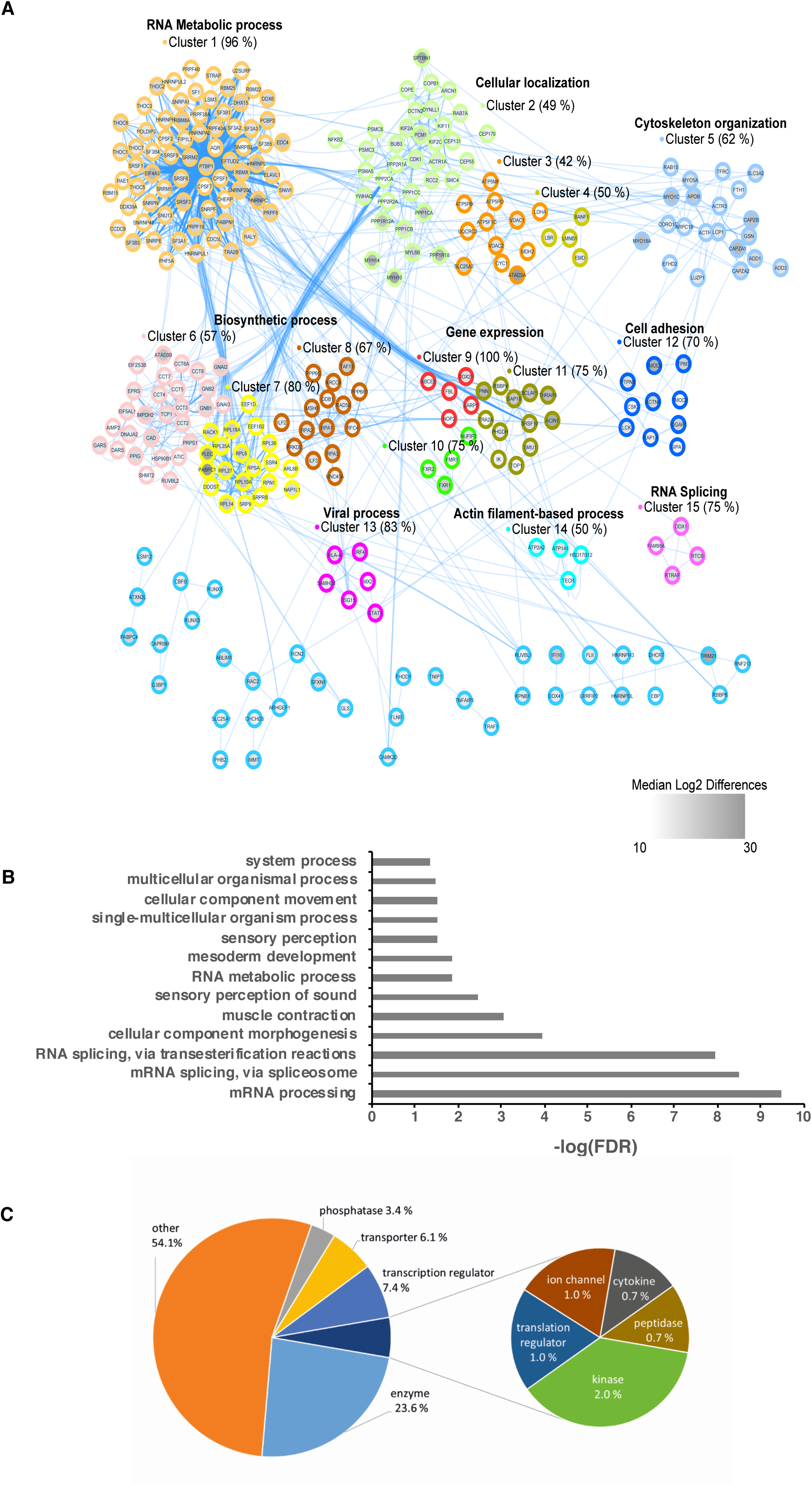
The CIP2A protein interaction network. **(A)** The interaction network among the CIP2A interacting proteins. Protein-protein interactions for the network were downloaded from the STRING database and visualized using the Cytoscape software. Nodes in the network were clustered using the Markov clustering algorithm. The identified clusters with more than two members are indicated by distinct colours. A representative GO biological process term is shown for each cluster with four or more members. The GO enrichment analysis was performed using DAVID against a Th17 proteome reference background (Tripathi et al., 2019). The median MS intensity differences between the CIP2A immuno-precipitates and the IgG controls are shown as node inner colour from white to grey according to increasingly stronger expression in the CIP2A immuno-precipitates compared to the IgG controls. **(B)** Enriched Biological processes associated with the proteins in the CIP2A interactome in human Th17 cells. The enrichment analyses was performed using PANTHER with the enrichment calculated from a background of proteins detected in Th17 proteomic reference background (Tripathi et al., 2019). The scale is –log FDR. **(C)** Functional classification of the CIP2A interactome in Th17 cells. The proportions of different enriched functional classes in the CIP2A interacting proteins. The enrichment analysis was performed using IPA (Qiagen).

To provide additional context and an appropriate milieu, functional enrichment analysis was made relative to a combined list of the proteins detected together with those found in a recent cellular proteomics analysis of human Th17 cells [29] rather than relative to a background of the whole genome. For this comparison the bioinformatics tools DAVID [30], [31] and PANTHER [32], [33] were used. The enriched Th17 pathways are represented in **Fig. 3B**. Again, mRNA processing and splicing were found to be the most significant biological process linked with CIP2A interactors. Collectively, these results suggest the involvement of CIP2A in multiple processes.

IPA analysis was also used to summarize the molecular functions of the CIP2A interacting proteins (**Fig. 3C)**. This revealed enzymes (∼24%) as the major functional class of the proteins. In addition, transcription regulators (∼7.4%) and transporters (6.1%) are the next major class of proteins. A small fraction of proteins was classified as phosphatases (3.4%), kinases (2.0%), translation regulators and ion channels (1.0% each), cytokines and peptidases (<1% each). For a major fraction of the proteins in CIP2A interactome (54%), however, analysis with the IPA database indicated that no specific function could be assigned (marked as “others” in **Fig 3C**). Overall the IPA analysis indicated that CIP2A interacts with proteins with a variety of different functions. It is possible that via its interaction with these proteins CIP2A regulates a range of processes in different cell types, some of which are as of yet undefined.

Thus, the functional and network analysis of the associated proteins revealed how CIP2A and its interacting partners participate in a broad set of essential cellular processes, ranging from RNA metabolic processes and splicing to viral processes.

### Validation of the Th17 cell CIP2A protein interactions

Validation experiments were performed to verify the interaction of selected proteins with CIP2A. The choice of targets focused on those with most significant interactions and nodal positions in CIP2A interactome (**Fig. 1D).** The validation experiments were carried out by confocal microscopy, WB, or selected reaction monitoring (SRM) targeted mass spectrometry (MS) analysis. The choice of method depended on the availability of suitable working antibodies or proteotypic peptide signatures, for confocal microscopy and WB or SRM-MS analysis, respectively.

SRM-MS analysis provided successful validation of twenty CIP2A interaction partners using immunoprecipitates from three independent biological replicates (**Fig 4A; Supplementary Table 4-5**). The results are summarized in **Supplementary Table 5.** We confirmed the known interaction of CIP2A with PP2A regulatory subunit A (PPP2R1A) **(Fig. 4A).** In addition, we identified and validated among the CIP2A interactors, PP2A catalytic subunits PPP2CA and PPP2CB **(Fig. 4A; S4A and Supplementary Table 5).**

**Figure 4.**
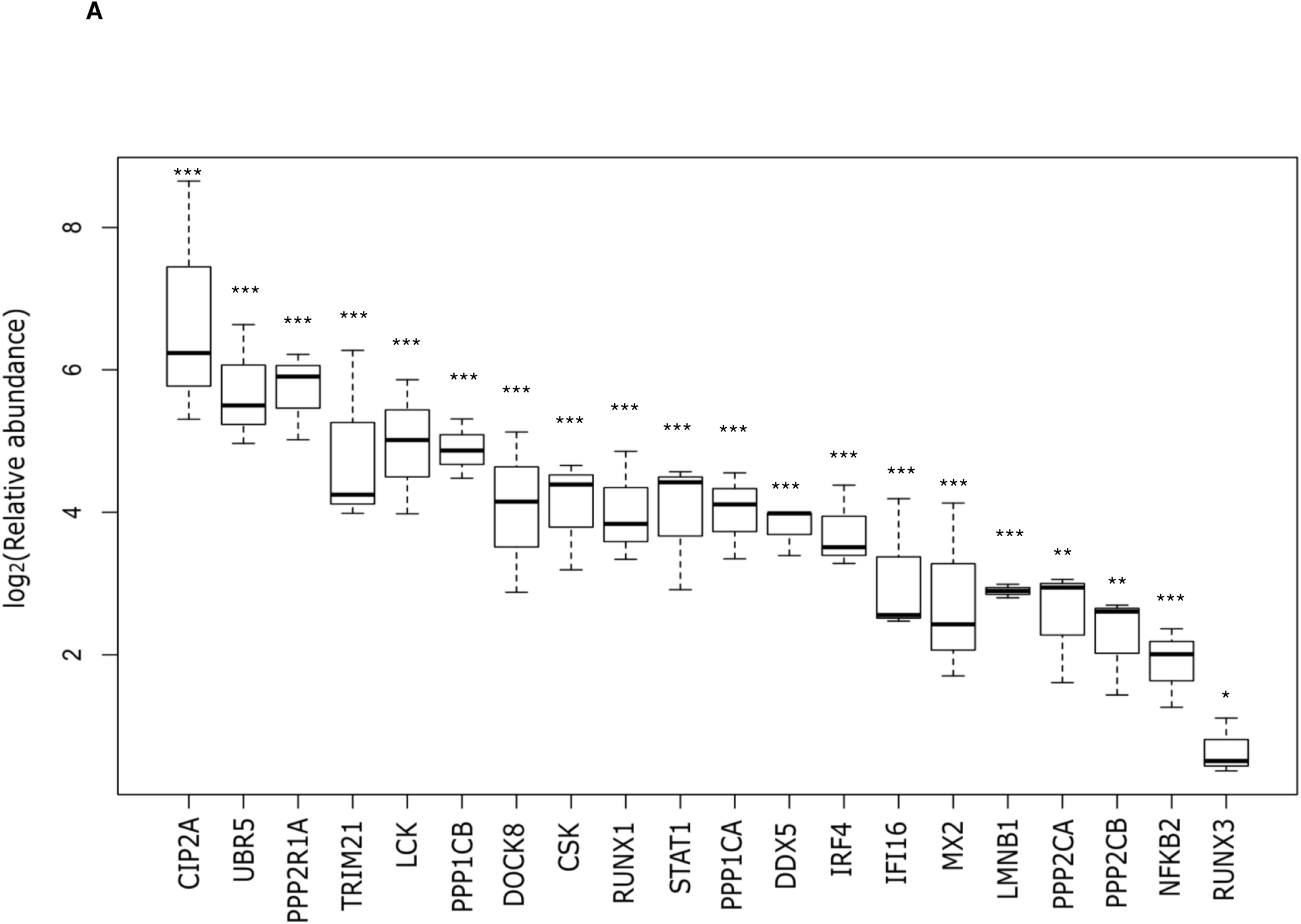
CIP2A interactome validation by SRM mass spectrometry. Selected reaction monitoring targeted mass spectrometry (SRM-MS) validation of the CIP2A interactome. CIP2A IP was performed using antibody Ab2 in 72h polarised Th17 cells and twenty proteins were validated by targeted SRM-MS. Averaged results from three replicates are presented in the form of a box plot. Statistical analysis was made by a two-tailed paired Student’s T-test, where * p-value <0.05; and *** p-value <0.001. The error bars represent 95% confidence interval.

Interestingly, we identified PP1 regulatory (PPP1R18 and PPP1R12A) and catalytic subunits (PPP1CA) as the top CIP2A interacting proteins (**Fig 1D)**. By SRM-MS analysis, further confirmed CIP2A interaction with the catalytic subunits (PPP1CA and PPP1CB) of phosphatase PP1 **(Fig. 4A; S4A and Supplementary Table 5)**. Further, in a recent phospho-proteome analysis of CIP2A-regulated proteins, LMNB1 was found to be dephosphorylated in CIP2A silenced cells [28] and here its interaction was identified and validated in CIP2A interactome by SRM MS analysis **(Fig. 4A).** Previous studies have revealed the interaction of with phosphatase PP2A regulatory A subunit [8]. Accordingly, we observed and validated this interaction with PP2A-A in Th17 cells (**Fig 5A).** Additionally, we validated the interaction with the PP2A catalytic subunit PPP2ACA. Notably, we were also able to validate interaction between CIP2A and PP1 catalytic subunit by co-immunoprecipitation followed by WB **(Fig. 5B).** Interaction of CIP2A with both PPP2ACA, and PPP1CA could be because of high structural similarity of the two proteins [34]. Further, PPP1CA interaction was independently validated by Proximity ligation assay (PLA) **(Fig. 5C)**. The negative control, PLA conducted from same samples without primary antibodies, did not show background signals (**Fig. 5C**).

**Figure 5.**
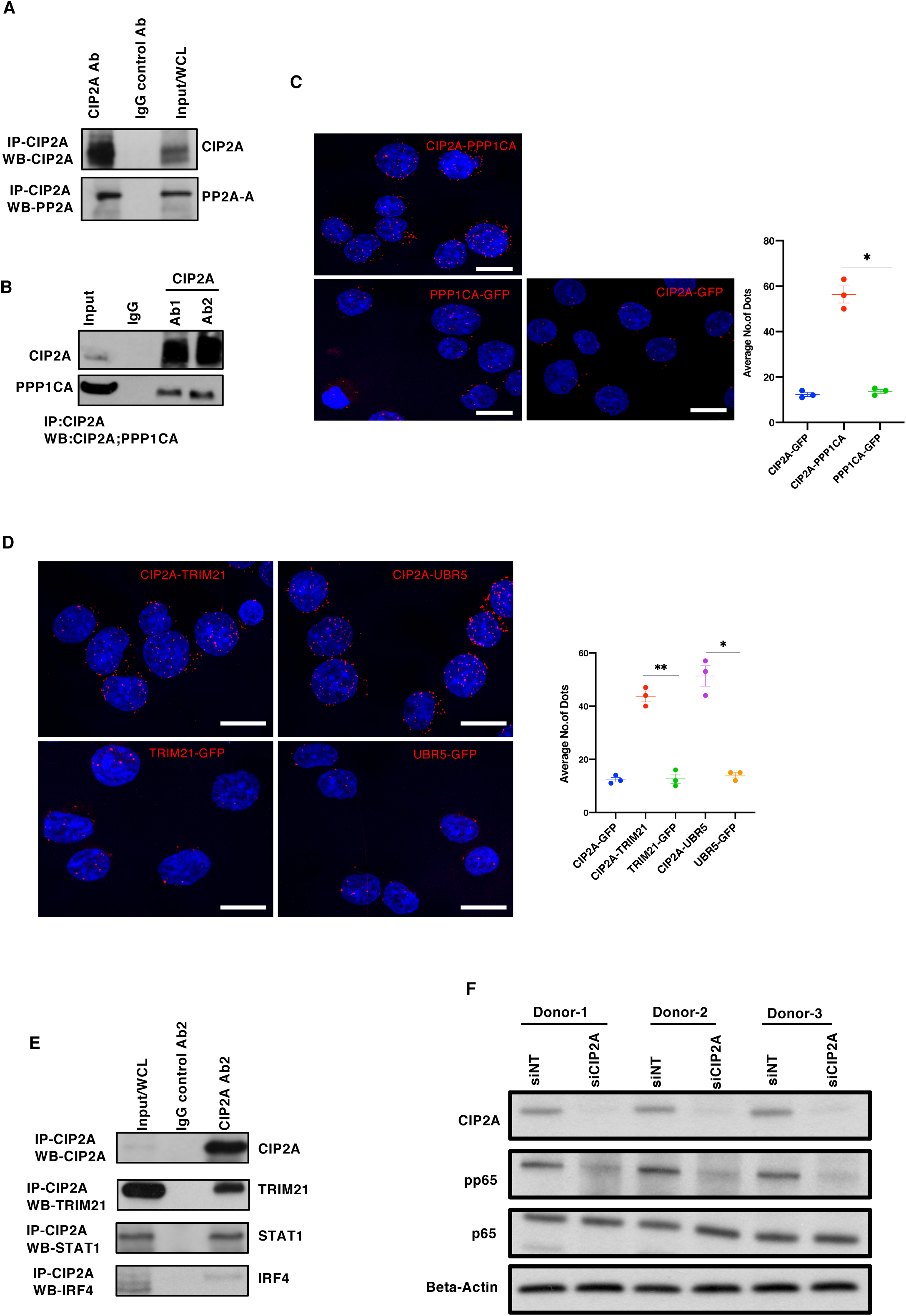
Validation of CIP2A known interaction by Western Blot and Confocal microscopy. **(A)** CIP2A IP performed in Th17 cells and WB detection to validates CIP2A known interaction with phosphatase PP2A-A. Total lysate (Input) and control IP (IgG) and CIP2A IP using Ab2 are shown on the blot. A conformational specific secondary antibody was used to probe proteins without interference from the denatured IgG heavy (50 kDa) and light chains (25 kDa). **(B)** WB validation of the CIP2A-PPP1CA interaction. CIP2A immunoprecipitated with two different antibodies (Ab1 and Ab2) in Th17 cells (72h) and detected as in A. **(C-D)** Proximity ligation assay (PLA) assay validation of the CIP2A-PPP1CA interaction (C) or selected proteins (TRIM21 and UBR5) interaction with CIP2A in Th17 cells (D). The GFP antibody was used as a negative control in PLA. DAPI was used to stain the nuclei. Bar is 7μm. Representation of the interaction between CIP2A-PPP1CA (C) or TRIM21 and UBR5 with CIP2A (D) by PLA as a univariate dot plot. Average number of PLA signals (spots) per cell (n=10 images from a representative of three experiments) was plotted. Statistics by paired two tailed T-test, where * represents P< 0.05. and **< 0.01. Scale bar is 7μm (**E**) WB validation represent interactions between TRIM21, IRF4, STAT1 and PP2A with CIP2A in Th17 cells at 72h and CIP2A. IP was performed using Ab2 and conformational specific secondary antibody was used as in Fig. 2C. **(F)** CIP2A inhibition leads to decreased TNFα-elicited Nf-κB activation. Total and phospho-p65 (pp65) Nf-κB subunit expression in CIP2A silenced and control Th17 cells. CIP2A is silenced in three different individual donor cell pools, and CIP2A, pp65, p65 and Actin are shown on the blot.

TRIM21 was amongst the most enriched proteins in this interactome, and the interaction was validated by both SRM-MS and PLA confocal microscopy **(Fig 5D).** Interaction with the ubiquitin ligase UBR5 was validated by PLA confocal microscopy, both in Th17 cells and Hela cells (**Fig 5D; Fig S5A**). In addition, WB analysis were made for STAT1, IRF4 and TRIM21, which were consistent with the targeted SRM-MS analysis (**Fig 4A; Fig 5E; Fig S5B-C**). Further, NFKB2 was also confirmed to be a CIP2A interacting protein (**Fig 4A, Fig S4A**) and CIP2A inhibition decreased NF-KB activation (**Fig 5F**). In CIP2A silenced Th17 cells, reduced phosphorylation of P65 was detected by WB analysis (**Fig 5F).**

## DISCUSSION

With these analyses of the CIP2A interactome in the Th17 milieu, and in relation to phosphatase biology, we verified the well-known interaction with PP2A-A (PPP2R1A) and PP2A regulatory B subunit **(Supplementary Table 1A-C)**. Previously, Wang et al. (2017) demonstrated direct interaction between CIP2A and PP2A regulatory B subunits B56α and B56γ by yeast two-hybrid analysis (Y2H), and Junttila et al. (2007) discovered PP2A-A subunit interaction with CIP2A. In the present report, we detected and validated for the first time the interaction of CIP2A with the PP2A catalytic subunit (PPPCA and PPPCB). All of these interactions were validated with SRM-MS analysis **(Fig. 4A; and Supplementary Table 4-5).**

The regulatory (PPP1R18 and PPP1R12A) and catalytic (PPP1CA) subunits of phosphatase PP1 were among the top CIP2A interacting partners. Interaction between CIP2A and PP1 has not been previously shown and further studies are required to demonstrate the importance of PP1 association with CIP2A. Notably, the catalytic subunits of both phosphatases PP2A and PP1 were detected and confirmed in the CIP2A interactome. This interaction could arise due to the structural similarity between catalytic subunits of phosphatases PP2A and PP1. It will be an interesting topic for future studies to determine whether CIP2A engages other catalytic subunits of serine/threonine phosphatases.

Previously in studies with T regulatory (Treg) cells, PP2A enhanced catalytic activity was identified as an indispensable element of the suppressive function of Treg cells [1]. Treg cells maintain tolerance and prevent autoimmune diseases by suppressing effector cells and modulating the immune system. It remains to be studied whether the interaction between CIP2A and catalytic subunit of PP2A regulates the suppressive function of Treg cells.

In relation to CIP2A and PP1 in autoimmune disease, in rheumatoid arthritis (RA) inflamed synovium, increased expression and activity of phosphatase PP1 and reduced transcriptional activity of FOXP3 through specific dephosphorylation of the Ser418 site has been observed [35]. Intriguingly, CIP2A expression in RA has been associated with the histopathological score of synovitis as well as apoptotic resistance and invasive function of fibroblast-like synoviocytes [36], [37]. Thus, interaction between CIP2A and PP1 may play a role in regulatory T cell function and inflammation in RA.

Several kinases related to T cell receptor (TCR) signaling (i.e. CSK and LCK) [38], [39] were identified as CIP2A interactors. The initiation of TCR signalling is regulated by the SRC-related protein tyrosine kinase family, which includes LCK, whereas CSK is a potent inhibitor of TCR signalling [38]. Notably, we have observed reduced T cell activation in CIP2A depleted T cells [21]. It will be interesting to study if the loss of CIP2A binding to LCK or CSK contributes to impaired T cell activation

The adaptor protein DOCK8 was amongst the most highly enriched proteins detected in this interactome. Combined immunodeficiency in humans is associated with loss-of-function mutations in the gene encoding DOCK8 [40]. Notably, the phenotype in immunodeficiency due to mutations in the gene encoding the transcription factor STAT3 is very similar to that caused by the loss of DOCK8 [41]. Jabara et al. reported that in B cells DOCK8 and STAT3 are components of the same signalling cascade [42]. In Th17 cells, CIP2A interacts with both DOCK8 and Lyn, and regulates STAT3 phosphorylation (Khan MM et al., submitted). It could be revealing to study whether CIP2A regulation of STAT3 phosphorylation is dependent on its interaction with Lyn and DOCK8.

Several of the proteins of this interactome have functions related to viral response and interferon signalling, i.e. IFI16, IRF4, MX2, STAT1 and TRIM21. Notably amongst these, the increased activity and DNA binding of IRF4 has previously been found in association with the increased capacity of PP2Ac transgenic T cells to produce IL-17 in response to TCR stimulation [2]. CIP2A directly interacts and inhibits PP2A in cancer cells and depletion of CIP2A in T cells results in enhanced IL17 (Khan MM et al., submitted manuscript). On the basis of these associations, the influence of CIP2A interaction on the DNA binding of IRF4 in T cells could be an important focus of future studies.

Amongst the CIP2A interacting transcription factors were RUNX1 and STAT1, which were validated by WB and SRM. Future studies could address the relationship between CIP2A and STAT1 in Th cells. RUNX1 is important for the development of Th17 cells as it directly binds the promoter of RORC and mediates transactivation of RORC, a transcription factor important for Th17 cell development [43]. In addition, Runx1 has been reported to bind to the IFNγ locus and induce IFNγ in Th17 cells resulting in development of pathogenic IL17^+^ IFNγ^+^ Th17 Cells [44]. In view of the latter and regarding the detected interaction between CIP2A and RUNX1, it will be of interest to find out if this interaction plays a role in RUNX1 mediated RORC and IFNγ induction in Th17 cells [43], [44].

TRIM21, an E3 ubiquitin ligase, was prominent in the CIP2A interactome. TRIM21 has been associated with the pathogenesis of several autoimmune diseases, including systemic lupus erythematosus (SLE) and inflammatory bowel diseases (IBDs). TRIM21 plays a protective role in IBD pathogenesis, possibly through inhibition of Th1 and Th17 cell immune responses in the intestinal mucosa [45] – [47]. In view of the detected interaction, it is possible that pathways involving CIP2A are also active in the pathogenesis of autoimmune diseases like SLE and IBD.

Beta-spectrin (SPTBN1) was amongst the top most proteins enriched in CIP2A interactome. Beta spectrin is a molecular scaffold protein that is involved in cell shape and the organisation of organelles in the cell [48]. Beta spectrin has been identified as an integral component of beta-amyloid plaques in Alzheimer’s disease (AD) and spectrin breakdown products (SBDPs) have been implicated as potential biomarkers for neurodegenerative diseases including AD [49], [50]. In a recent study of AD, CIP2A was found to be overexpressed and associated with tau hyperphosphorylation, mis-localization and AD-related cellular pathology [14]. In mice, overexpression of CIP2A in the brain induced AD-like cognitive deficits [14]. Future studies may determine whether CIP2A interaction with beta-spectrin modulates protein aggregation in AD pathology.

In summary, the results of this study present the first characterisation of CIP2A interactome. In addition to the known interactions with the PP2A protein A and B regulatory subunits, our results detail several novel CIP2A interacting proteins. These include the catalytic subunit of PP2A, as well as regulatory and catalytic subunits of PP1. Gene ontology and pathway analysis of the associated proteins suggest CIP2A to be involved in several cellular functions that it has not been previously linked to. Taken together, we anticipate the new information of proteins depicted in this CIP2A interactome and their associated networks, could be useful in understanding the role of CIP2A in regulating cell signalling and its role in various human disorders.

## Methods

### Human CD4+ T cell isolation, activation, and differentiation

Umbilical cord blood was layered on Ficol (GE health Care # 17-1440-03) for isolation of white blood cells. CD4+ T cells were then isolated using the Dynal bead CD4+ isolation kit from Invitrogen (Cat #11331D). For activation of T cells, a combination of plate-bound anti-CD3 (Beckman Coulter REF # IM-1304) and soluble anti-CD28 (Beckman coulter REF # IM1376) antibodies was used. The Th17 cultured cells were stimulated in presence of IL-6 (20ng/ml, Roche, Cat# 11138600 001); IL-1β (10ng/ml, R&D Systems Cat# 201 LB); TGF-β1 (10ng/ml, R&D Systems Cat# 240); anti-IL-4 (1 ug/ml) R&D Systems Cat# MAB204); anti-IFN-γ (1 μg/ml R&D Systems Cat#MAB-285).

### Flow cytometry

For Th17 and activated T cells, CCR6 surface staining was performed for Th17 cell culture polarization control. Cells were stained with the antibody (eBioscience) in FACS I buffer (1%FBS in PBS) at +4 °C in darkness for 30min. Cells were then washed two times with FACS I, and finally re-suspended in FACS I Buffer or 1% Formalin before acquisition in BD LSR II/ Caliber. Data analysis was performed using FlowJo (BD) or Flowing Software (Version 2.5.1).

### Luminex assay

As a polarization control, IL17 cytokine protein secretion was detected in 72h polarized human Th17 cells and TCR activated control cells. Luminex assay (Milliplex MAP human cytokine, Luminex 200 by Luminex xMAP technology) was used to detect the IL17 cytokine in the supernatant and measured according to the manufacturer’s instructions. To avoid cell proliferation bias, the cytokine concentrations were normalized by the number of cells from each respective culture, as counted by flow cytometry.

### Immunoprecipitation

CIP2A immunoprecipitation (IP) was carried out with two specific anti-CIP2A antibodies that recognize distinct regions of CIP2A protein and the respective Immunoglobulin G (IgG) were used in control IPs. The Pierce™ MS-Compatible Magnetic IP Kit, (Catalog number: 90409) was used to perform IP. Briefly, cells were first harvested on ice and the culture medium removed. This was followed by washing the cells twice with PBS. Cells were then re-suspended on ice-cold IP-MS Cell Lysis Buffer, as recommended by the manufacturer, and incubated on ice for 10 minutes with periodic mixing. The lysate was centrifuged at ∼13,000 × g for 10 minutes to pellet the cell debris. The supernatant was transferred to a new tube for protein concentration determination using a NanoDrop™ spectrophotometer (Thermo Scientific). Cell lysates (500μg) were incubated with the IP antibody (1:50 dilution). The antibody/lysate solution was diluted with the manufacturer’s proprietary IP-MS cell lysis buffer and incubated with mixing overnight at +4°C to form the immune complex. On the next day the immune complex was incubated with Protein A/G magnetic beads for 1h at room temperature. The beads were then washed and the target antigen eluted with the elution buffer provided and transferred to a new low protein-binding 1.5mL microcentrifuge tube. A vacuum centrifuge was used to dry the eluate prior to preparation for MS or SDS-PAGE and Western blot.

### Western Blotting

T cells were first resuspended in Triton-X-100 lysis buffer (TXLB). The composition of TXLB used was 50 mM Tris-HCl (pH 7.5), 150 mM NaCl, 0.5% Triton-X-100, 5% Glycerol, 1% SDS. For cell lysis, T cells in TXLB were then sonicated in ice for 5 minutes using a Bioruptor® sonicator. After cell lysis, the lysate was centrifuged at high speed (16,000 g) and the supernatant was transferred to a new tube to remove cell debris. Protein concentration was estimated using a DC Protein assay (Bio-Rad Cat#500-0111). Equal amounts of protein were loaded onto the acrylamide gel (Bio-Rad Mini or Midi PROTEAN® TGX precast gels). For the transfer of proteins to the PVDF membrane, mini or a midi transfer packs from Bio-Rad were used depending on the gel size. Primary/ secondary antibody incubations were performed in 5% non-fat milk or Bovine serum albumin (BSA) for phosphoproteins in TBST buffer (0.1%Tween 20 in Tris-buffered saline). Quantification of the bands detected was performed using Image J software after normalization relative to the loading control intensities. The following antibodies were used for western blotting, CIP2A antibody 1(Ab1) (Cell Signaling Technology; cat#14805) CIP2A antibody 2 (Ab2) provided by Prof. Jukka Westermarck [8], [22]; PP2A-A/B (sc-15355; Santa Cruz); Trim21 antibody (Santa Cruz); Irf4 antibody (Santa Cruz); Beta-actin antibody (Sigma Cat #. A5441). PPP1A/PPP1CA antibody (ab137512; abcam), PP1α Antibody (G-4): (sc-271762; Santa Cruz).

### Immunofluorescence

Th17 cells were plated onto acid-washed glass coverslips. The cells were then fixed at room temperature (RT) using 4 % PFA for 10 mins. After fixing, the cells were permeabilized in PBS containing 30% horse serum and 0.1% Triton X-100 for 10 min at RT, washed in PBS and then blocked in PBS-30% horse serum for 1h at RT. The cells were then incubated with the CIP2A primary antibody overnight at 4°C. On the next day, the cells were washed three times using PBS and incubated for 1h at RT with AlexaFluor-conjugated secondary antibodies (Life Technologies). Cells were then mounted with Mowiol® for 30 min at 37°C and Images taken with a Carl Zeiss LSM780 laser scanning confocal microscope equipped with 100×1.40 oil plan-Apochromat objective (Zeiss). Corrected total cell fluorescence was calculated from confocal images to quantify the intensities in Th0 and Th17 cells. [51], [52].

Immunofluorescence staining of UBR5 in Hela cells was performed similarly, the cells were cultured on chambered coverslip (80826, Ibidi). 4% paraformaldehyde was used to fix the cells for 15 minutes under room temperature, and then cells were permeabilized with 0.5% Triton X-100 in PBS on ice for 5 minutes. Next, the cells were blocked with 10% normal goat serum (ab7481, Abcam) diluted in PBS for 30 minutes, and followed by incubating with the primary antibodies anti-UBR5 (ab70311, Abcam) overnight at 4°C. Subsequently, cells were washed with PBS and incubated with the secondary antibody, Alex Fluor 488 goat anti-rabbit IgG (A-11008, Invitrogen) for 1h under room temperature. After secondary antibody incubation, the cells were washed with PBS and nuclei were stained with DAPI (D1306, Invitrogen) in PBS at RT for 10 min. Images were acquired with confocal microscope (LSM780, Carl Zeiss).

### Proximity ligation assay

The PLA assay was performed according to the manufacturer’s protocol (Duolink® PLA, Sigma). Briefly, Th17 or HeLa cells were plated on coverslips. The cells were fixed with 4% paraformaldehyde for 15 minutes at room temperature, and then permeabilized with 0.5% Triton X-100 in PBS on ice for 5 minutes. Next, cells were blocked with blocking solution, and incubated in a pre-heated humidity chamber for 30 min at 37°C, followed by incubating with the primary antibodies (in blocking solution) anti-UBR5 (ab70311, Abcam) and anti-CIP2A (sc-80659, Santa Cruz), anti-PPP1CA (ab137512, Abcam), anti-UBR5 (ab70311, Abcam), anti-TRIM21 (SC25351, Santa Cruz), anti-GFP (Mouse, ab1218, Abcam) and anti-GFP (Rabbit, A11122, Invitrogen) overnight at 4°C. Subsequently, cells were washed with the kit buffer A, and the PLA probe was incubated in a pre-heated humidity chamber for 1h at 37°C, followed by ligase reaction in a pre-heated humidity chamber for 30 minutes at 37°C. Next, amplification polymerase solution for PLA was added, followed by incubating the cells in a pre-heated humidity chamber for 100 minutes at 37°C. After amplification, the coverslips were washed with the kit buffer B, and mounted with DAPI. The PLA signal was detected by using a confocal microscope (LSM780, Carl Zeiss)) and 3i CSU-W1 spinning disk microscope equipped with 100×1.4 O Plan-apochromat objective (Zeiss). PLA signals per cell were calculated by dividing the amount of PLA signal dots in one field of view, determined using Cell Profiler software [53]

### Cell transfection with siRNA

Naïve T cells were isolated form the cord blood and four million cells were transfected in 100 μl volume of Opti-MEM™ (Gibco by Life Technologies, cat # 31985-047). Lonza nucleofector was used for siRNA transfections. After nucleofection cells were rested for 48h in RPMI supplemented with 10% serum.

The siRNAs (synthesized by Sigma) are shown below:

siNT 5’-AAUUCUCCGAACGUGUCACGU-3’ 22;
siCIP2A-1 5’-CUGUGGUUGUGUUUGCACUTT-3’1.

### LC-MS/MS analysis of the proteomic profiles of the immuno-precipitates

The immuno-precipitates were denatured with 8 M urea and cysteine residues reduced with dithiothreitol then alkylated with iodoacetamide. After digestion with trypsin (37 ^0^C, 16 hours), the samples were desalted using Empore C_18_ disks (3M, Cat No 2215). Based on using UV absorption measurements with a NanoDrop-1000 UV spectrophotometer (Thermo Scientific), equivalent aliquots of the digested peptides were analyzed in triplicate with an Easy-nLC 1200 coupled to a Q Exactive HF mass spectrometer (Thermo Fisher Scientific). A 20 x 0.1 mm i.d. pre-column was used in conjunction with a 150 mm x 75 µm i.d. analytical column, both packed with 5 µm Reprosil C_18_-bonded silica (Dr Maisch GmbH). A separation gradient from 8 to 45% B in 78 min was used at a flow rate of 300 nl/min. The mobile phase compositions were, water with 0.1 % formic acid (A) and 80% acetonitrile 0.1% formic acid (B). The MS/MS data were acquired in positive ion mode with a data dependent acquisition setting for higher-energy C-trap dissociation (HCD) of the 10 most intense ions (m/z 300-2000, charge states > 1+). Further details of the preparation and instrument parameters are available as supplementary information.

The LC-MS/MS discovery data are available via the PRIDE [54] partner repository from the ProteomeXchange Consortium with the dataset identifier PXD008983.

### LC-MS/MS validation of selected targets

Selected reaction monitoring (SRM) mass spectrometry was used to validate the relative abundance of twenty of the proteins that were found to be associated with the CIP2A immuno-precipitates (**Supplementary Table 4**). Heavy-labeled synthetic peptides (lysine ^13^C_6_ ^15^N_2_ and arginine ^13^C_6_ ^15^N_4_) analogues for the targets of interest were selected on the basis of their consistency and intensity in the discovery data and obtained from Thermo Scientific. For these validations, three additional cultures were prepared from the cord blood of three donors.

Using Skyline software [55] the top five most intense transitions for each peptide were evaluated from the MS/MS spectra of heavy labelled peptides and the relative performance of the native peptides in the spiked validation samples.

The validation samples were prepared using the same digestion and desalting protocols used for discovery. These were then spiked with synthetic heavy labelled analogues of the peptide targets and a retention time standard (MSRT1, Sigma) for scheduled selected reaction monitoring. The LC-MS/MS analyses were conducted using Easy-nLC 1000 liquid chromatograph (Thermo Scientific) coupled to a TSQ Vantage Triple Quadrupole Mass Spectrometer (Thermo Scientific). The column configuration was the same as used in the discovery experiments and the peptides were separated using a gradient of 8% to 43% B in 27 min, then to 100% B in 3 min, at a flow rate of 300 nl/min (the mobile phase compositions are as indicate above). The raw SRM data are available through PASSEL [56] with the dataset identifier PASS01186

### Data analysis and statistics

The peptide/protein identification was performed in Andromeda search engine [57], incorporated in MaxQuant software [58], with a UniProt human protein sequence database (version June 2016, 20,237 entries). Trypsin digestion with a maximum of two missed cleavages, carbamidomethylation of cysteine as a fixed modification, and methionine oxidation and N-terminal acetylation as variable modifications were used in the search parameters. A false discovery rate (FDR) of 0.01 at the peptide and protein level was applied. MaxQuant’s label-free quantification (LFQ) algorithm [59] was used to calculate the relative protein intensity profiles across the samples. The “match between the runs” option was enabled with a time window of 0.7 min.

The MaxQuant output was further processed using Perseus [60]. The downstream processing included the removal of contaminants, reverse hits and proteins with less than three valid replicate values in the sample groups (IgG or CIP2A pulldown). To identify the potential interacting proteins, Significance Analysis of *INTeractome* (*SAINT*) analysis was performed [24] using the Resource for Evaluation of Protein Interaction Networks (REPRINT) interface (https://reprint-apms.org/). This algorithm uses the distribution, relative intensity and variation of the prey protein detection to calculate probability scores for the certainty of their interaction with the bait. Additionally, the protein lists were compared against a list of common contaminants identified in a catalogue of immunoprecipitation experiments (CRAPome). The thresholds applied to define the putatively interacting proteins included identification on the basis of two or more unique peptides and a SAINT probability score of ≥ 0.95. Although there was no direct match for our sample type and IP conditions in the contaminants repository database, additional filtering was made to exclude proteins that were more than 60% of the datasets. Skyline was used to design the SRM experiments and process the data generated. The MS Stats (3.8.4) plugin included in the Skyline software was used for the group comparison between cases and controls [61].

### Network analysis

Previously reported protein-protein interactions for the detected CIP2A interacting proteins were downloaded from the STRING functional protein association networks database version 10.5 [26]. Both predicted and known high confidence (interaction score ≥ 0.7) interactions were considered. The resulting network was visualized with Cytoscape version 3.6.0 [27]. Clusters in the network were identified using the Markov clustering algorithm implemented in the Cytoscape plug-in clusterMaker v2 algorithm MCL Cluster [62] with the granularity parameter (inflation value) set to 1.8 as suggested by [63]. The MS intensity differences between CIP2A immuno-precipitates and the IgG controls were calculated as median differences across all replicates in the normalized log2-transformed intensities (color coded as a continuous gradient from white to grey as node inner color in Figure 3A).

### Enrichment analysis

The enriched GO biological processes were identified using the Database for Annotation, Visualization and Integrated Discovery (DAVID) version 6.8 [30], [31] and the Protein Annotation Through Evolutionary Relationship (PANTHER) classification system version 13.0 [32], [33]. The enrichment analyses were performed using the detected CIP2A interactome as the input and the whole detected Th17 proteome from a Th17 profiling study [29] as the background reference. The GO FAT terms from DAVID, which filter out the broadest, non-specific terms, were considered for calculating the most enriched processes within each identified cluster with more than four members. The GO SLIM terms from PANTHER, which give only the broadest terms and filter the more specific ones, were considered for a summary of the enriched biological processes in the CIP2A interactome. A biological process was considered enriched if it had FDR ≤ 0.05. To summarize the cellular locations and protein classes the protein list was annotated using Ingenuity Pathway Analysis (Qiagen).

## ACKNOWLEDGEMENTS

For the cord blood collections, we are thankful to the staff of Turku University Hospital, Maternity Ward, Department of Obstetrics and Gynecology. Marjo Hakkarainen and Sarita Heinonen are acknowledge for excellent technical help. We are thankful to the core facilities at the Turku Bioscience centre namely, the Proteomics facility and Cell Imaging Core (CIC) facility. Biocenter Finland are acknowledged for their support of the latter facilities. The Finnish Centre for Scientific Computing (CSC) is duly acknowledged for their efficient servers and their resources in data analysis. MMK was supported by University of Turku graduate school on Turku Doctoral Programme of Molecular Medicine (TuDMM) as well as a central grant from Finnish Cultural Foundation and T.V. was supported by the Doctoral Programme in Mathematics and Computer Sciences (MATTI) of University of Turku. R.L. was supported by the Academy of Finland, AoF, Centre of Excellence in Molecular Systems Immunology and Physiology Research (2012-2017) grant 250114; by the AoF grants 292335, 294337, 292482, 31444, by grants from the JDRF, the Sigrid Jusélius Foundation and the Finnish Cancer Foundation. L.L.E. has received grants from the European Research Council (ERC; grant 677943), the European Union’s Horizon 2020 research and innovation program (grant 675395), the AoF (grants 296801 and 304995), the Juvenile Diabetes Research Foundation (JDRF; grant 2-2013-32), the Finnish Funding Agency for Innovation (TEKES; grant 1877/31/2016), and the Sigrid Jusélius Foundation. J.W. was funded by the Sigrid Jusélius Foundation.

## ETHICAL APPROVAL

Usage of the blood of unknown donors is approved by the Ethics Committee, Hospital District of Southwest Finland.

## AUTHOR CONTRIBUTIONS

M.M.K designed and performed the experiments, analyzed data and prepared figures related to WB, TaqMan-PCR and Flowcytometry, and wrote the manuscript. T.V. analyzed data, prepared figures for network analysis and wrote part of the manuscript. M.H.K. designed and performed the experiments, prepared figures, and wrote part of the manuscript. R.M performed SRM-proteomics, analyzed the proteomics data, and contributed to writing the manuscript. U.U. contributed to the pathway and network analyses and writing of the manuscript. S.D.B performed the proteomics experiments data analysis. E.K performed experiments, and analyzed the data. JW provided scientific expertise, and reagents and edited the manuscript. L.E. designed experiments, supervised the study and edited the manuscript. R.L. designed and supervised the study and wrote the manuscript.

## ACCESSION NUMBERS

The mass spectrometry data for the interactome have been deposited with the ProteomeXchange Consortium via the PRIDE [54] partner repository with the dataset identifier PXD008983. The validation data have similarly been submitted to the Peptide Atlas [64] with the identifier PASS01186.

## SUPPLEMENTARY FIGURE LEGENDS

**Figure S1 (Related to Figure 1).**
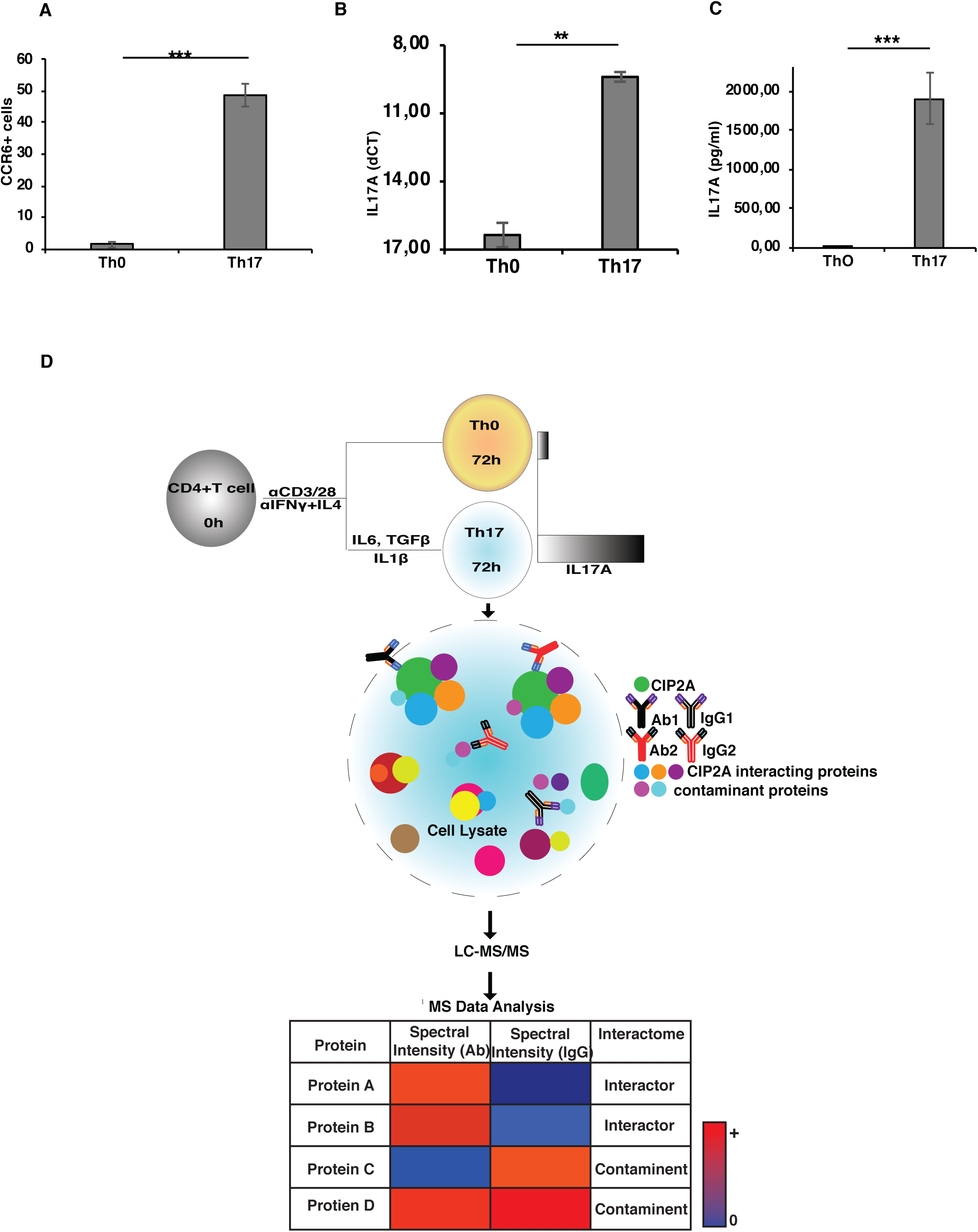
The polarization of the Th17 cultures used in the study and a schematic representation of the interactome study. **(A)** The proportion of CCR6+ cells from the flow-cytometry analysis of CCR6 surface expression at 72h in TCR activated control Th0 cells and polarized Th17 cells. The IL17A mRNA expression (**B**), and IL17A protein secretion (**C**) from the cultures at 72h post initiation of Th17 polarization. The data (**A**-**C**) is from six independent cultures and each culture consisted of cells from more than four individual donors. Error bars represent SEM estimates across biological replicates and P-values calculated using the paired two-tailed student’s t-test. (n = 6; P <0.01(**) and P <0.001(***) for Th0 vs Th17). (**D**) A schematic layout representation of the interactome study.

**Figure S2 (Related to Figure 2).**
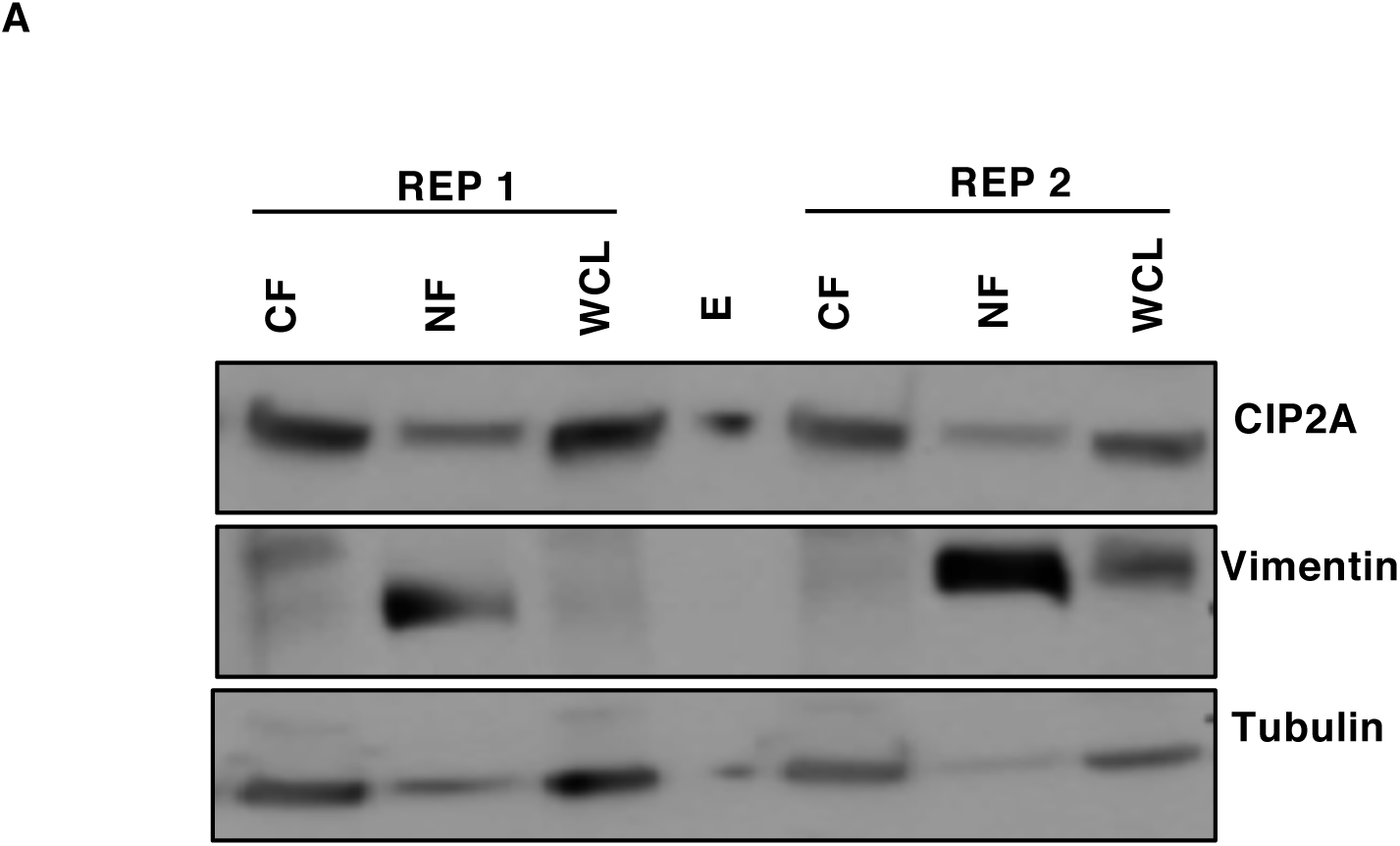
WB analysis of two replicates for the cytoplasmic and nuclear fractions for CIP2A sub-cellular localisation in 72h polarised Th17 cells. WB probes with antibodies as controls for nuclear and cytoplasmic fractions detecting Vimentin and Tubulin proteins, respectively. WCL, NF and CF represent Whole Cell lysate, Nuclear Fraction, and Cytoplasmic Fraction respectively.

**Figure S3 (Related to Figure 3).**
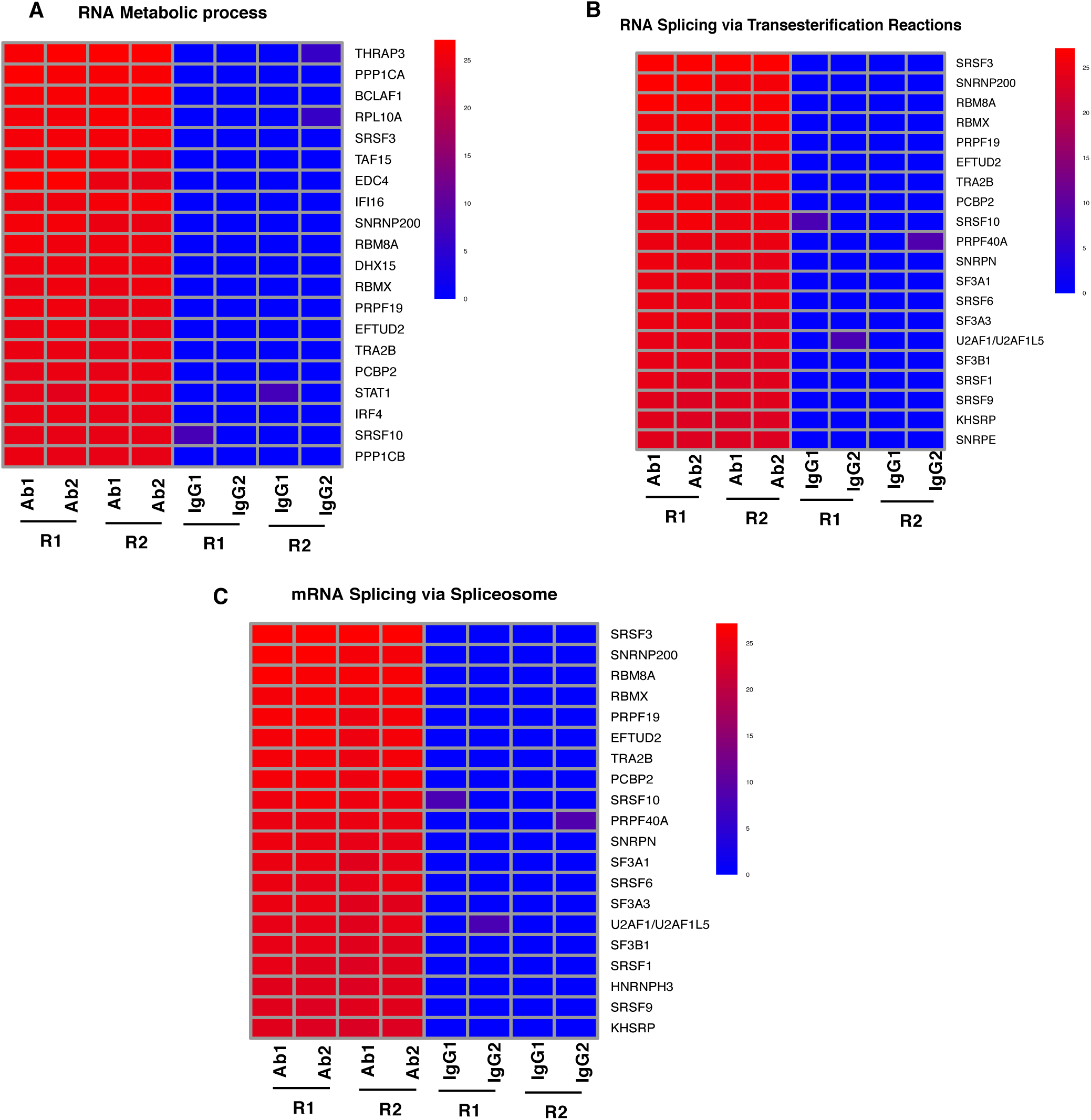
The normalized expression of the top CIP2A interacting proteins in selected highly enriched GO biological processes in the detected CIP2A interactome. The 20 strongest CIP2A interacting proteins in the **A**) RNA metabolic process (**B**) RNA splicing via transesterification and (**C**) RNA splicing via spliceosome enriched GO biological processes. The expression values are normalized log_2_ transformed protein intensities from the MS analysis of the CIP2A immuno-precipitates (IP) and the IgG controls. The expression values for both antibodies (Ab1 and Ab2) specific to different regions for two individual replicates are shown. The proteins in each heatmap are in descending order based on the median expression differences between the CIP2A-IPs and the IgG controls.

**Figure S4 (Related to Figure 4).**
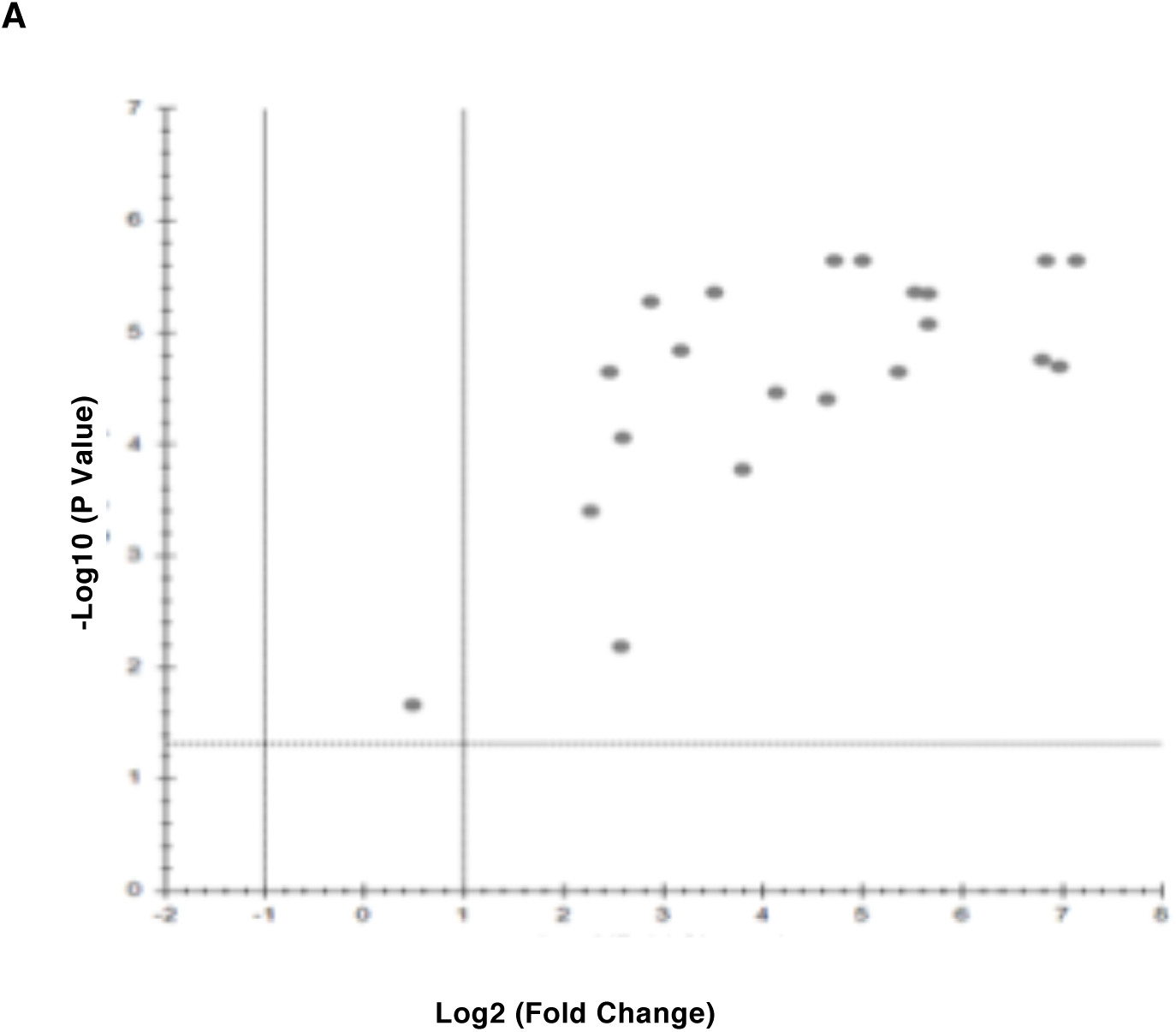
The averaged logarithmic fold changes and logarithmic significance values (log FDR) for the targeted mass spectrometry (SRM-MS) validations analysis of the twenty selected (shown in Figure 4A) proteins from the CIP2A interactome.

**Figure S5 (Related to Figure 5).**
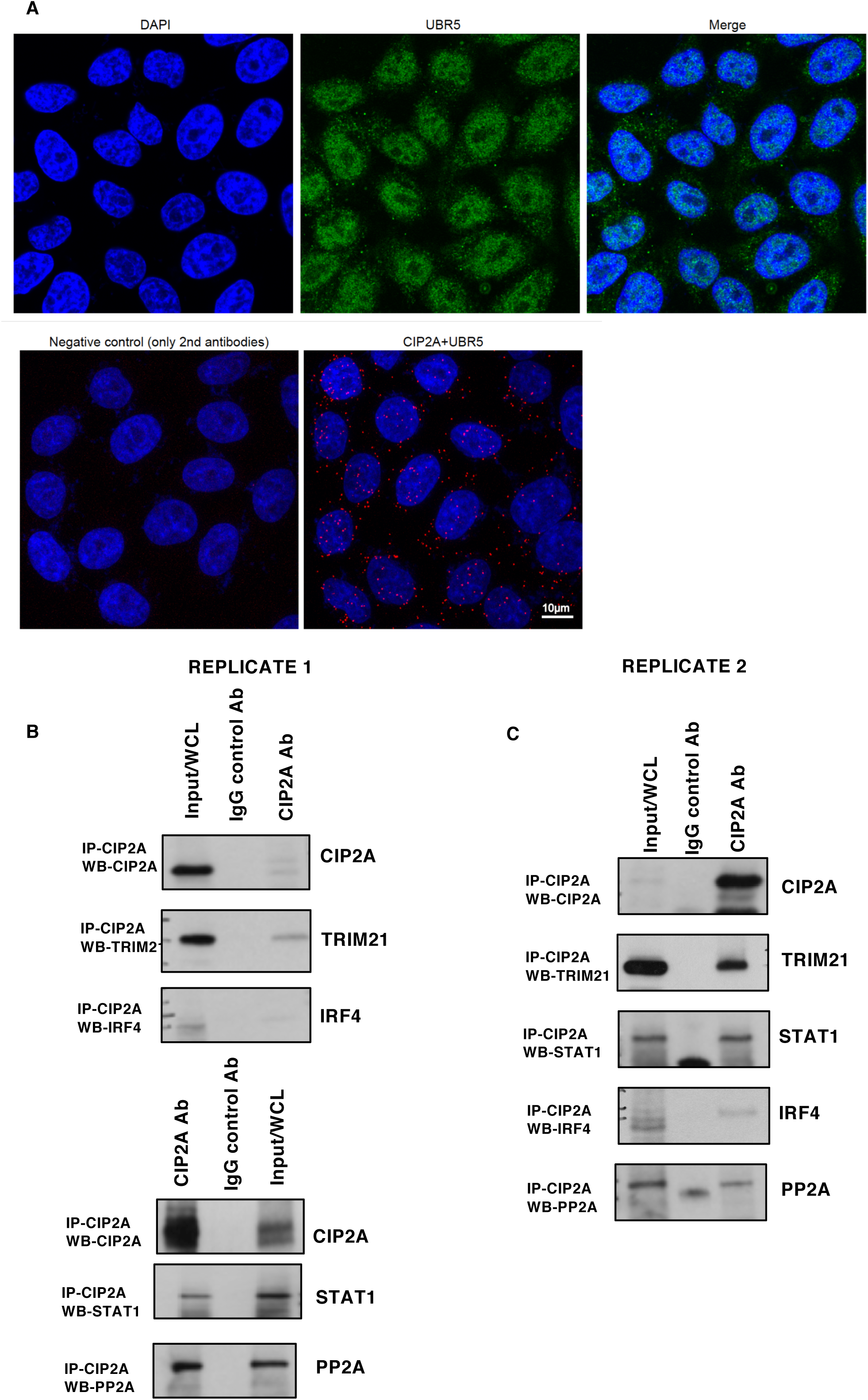
(**A**) UBR5 immunostaining in Hela cells and PLA validation for interaction between CIP2A and UBR5 by confocal microscopy. For PLA, the antibodies were used as negative controls. DAPI was used to stain nuclei. Bar = 10μm **(B-C)** WB validations of selected proteins in CIP2A interactome (Two replicates). (**B**) First replicate showing two separate blots, TRIM21 and IRF4 (first blot) and STAT1 and PP2A (second blot). (**C**) WB validations of the second replicate for the same proteins. CIP2A pull-down in 72h polarised Th17 cells and CIP2A IP, IgG control IP and Input reactions are shown on the blot. CIP2A IP was performed using Ab2 and conformational specific secondary antibody was used to probe proteins without interference from denatured IgG heavy (50 kDa) and light chains (25 kDa).

## SUPPLEMENTRY TABLE LEGENDS

**Supplementary Table S1 (A-C). CIP2A interactome in Th17 cells. (**Related to Fig. 1). Table list proteins associated with the CIP2A interactome (**Supplementary Table S1A)**. The table also list Ingenuity Pathway Analysis (IPA) cellular location and functional class assigned IPA annotation for the proteins detected in the CIP2A interactome (**Supplementary Table S1B)**. The CIP2A interactome associated proteins information are listed together with the data from the MaxQuant results (unique peptides, sequence coverage, intensity and Andromeda score), representation in the CRAPome, and SAINT analysis showing Saint Probability (SP) (**Supplementary Table S1C)**. The data is filtered to only include proteins with a SP values of > 0.95 and less than 60% representation in the CRAPOME.

**Supplementary Table S2.** (Related to Fig. 3). **The 20 most enriched GO FAT biological processes (BP) in each cluster of the CIP2A interactome network in** Figure 3A. The GO enrichment was performed using DAVID (Huang et al., 2009) against a Th17 proteome reference background (Tripathi et al., 2019). The GO FAT BP terms, which filter out the broadest higher level GO terms, were considered for more specific enrichment information. For each cluster with four or more members, 20 of the most frequently occurring GO FAT terms among the cluster members are listed. For each term, the number of cluster members (proteins or nodes) with the term, the proportion of cluster members with the term, cluster number and annotation for the cluster in Figure 3A, is shown.

**Supplementary Table S3. (**Related to Fig. 3). **The enriched GO biological processes in the detected CIP2A interactome.** The enrichment analysis was performed using PANTHER (Mi et al., 2013, 2017) against a Th17 proteome reference background (Tripathi et al., 2019). The broad GO SLIM terms, which include only the higher level GO terms, were considered for a functional summary of the interactome. Fold enrichment, significance value (p-value) and false discovery rate is presented for each term.

**Supplementary Table S4. SRM Targets for the CIP2A interactome validations. (**Associated with Fig. 4). The table lists the peptides, associated transitions, collision energies and retention time windows used for the SRM validations of the CIP2A vs. control IgG pull downs.

**Supplementary Table S5. SRM MS CIP2A interactome validations. (**Associated with Fig. 4). The table lists the log_2_ transformed relative abundance data from the SRM analysis of the CIP2A IP vs. control IgG pull downs from three biological replicates for proteins validated for the CIP2A interactome.

## REFERENCES

1. Apostolidis SA, Rodríguez-Rodríguez N, Suárez-Fueyo A, Dioufa N, Ozcan E, Crispín JC, et al. Phosphatase PP2A is requisite for the function of regulatory T cells. Nat Immunol. 2016;17(5):556–64.

2. Apostolidis SA, Rauen T, Hedrich CM, Tsokos GC, Crispín JC. Protein phosphatase 2A enables expression of interleukin 17 (IL-17) through chromatin remodeling. J Biol Chem. 2013;288(37):26775–84.

3. Eitelhuber AC, Warth S, Schimmack G, Düwel M, Hadian K, Demski K, et al. Dephosphorylation of Carma1 by PP2A negatively regulates T-cell activation. EMBO J. 2011;30(3):594–605.

4. Janssens V, Goris J. Protein phosphatase 2A: a highly regulated family of serine/threonine phosphatases implicated in cell growth and signalling. Biochem J. 2001 Feb 1;353(Pt 3):417–39.

5. Katsiari CG, Kyttaris VC, Juang Y-T, Tsokos GC. Protein phosphatase 2A is a negative regulator of IL-2 production in patients with systemic lupus erythematosus. J Clin Invest. 2005 Nov;115(11):3193–204.

6. Long L, Deng Y, Yao F, Guan D, Feng Y, Jiang H, et al. Recruitment of phosphatase PP2A by RACK1 adaptor protein deactivates transcription factor IRF3 and limits Type I interferon signaling. Immunity. 2014;40(4):515–29.

7. Wlodarchak N, Xing Y. PP2A as a master regulator of the cell cycle. Crit Rev Biochem Mol Biol. 2016;51(3):162–84.

8. Junttila MR, Puustinen P, Niemelä M, Ahola R, Arnold H, Böttzauw T, et al. CIP2A Inhibits PP2A in Human Malignancies. Cell. 2007;130(1):51–62.

9. Laine A, Sihto H, Come C, Rosenfeldt MT, Zwolinska A, Niemelä M, et al. Senescence Sensitivity of Breast Cancer Cells Is Defined by Positive Feedback Loop between CIP2A and E2F1. Cancer Discov. 2013 Feb 1;3(2):182–97.

10. Niemelä M, Kauko O, Sihto H, Mpindi J-P, Nicorici D, Pernilä P, et al. CIP2A signature reveals the MYC dependency of CIP2A-regulated phenotypes and its clinical association with breast cancer subtypes. Oncogene. 2012 Sep 16;31(39):4266–78.

11. Khanna A, Pimanda JE, Westermarck J. Cancerous Inhibitor of Protein Phosphatase 2A, an Emerging Human Oncoprotein and a Potential Cancer Therapy Target. Cancer Res. 2013 Nov 15;73(22):6548–53.

12. Lucas CM, Milani M, Butterworth M, Carmell N, Scott LJ, Clark RE, et al. High CIP2A levels correlate with an antiapoptotic phenotype that can be overcome by targeting BCL-XL in chronic myeloid leukemia. Leukemia. 2016 Jun;30(6):1273–81.

13. Wang J, Okkeri J, Pavic K, Wang Z, Kauko O, Halonen T, et al. Oncoprotein CIP2A is stabilized via interaction with tumor suppressor PP2A/B56. EMBO Rep. 2017 Mar 7;18(3):437–50.

14. Shentu Y-P, Huo Y, Feng X-L, Gilbert J, Zhang Q, Liuyang Z-Y, et al. CIP2A Causes Tau/APP Phosphorylation, Synaptopathy, and Memory Deficits in Alzheimer’s Disease. Cell Rep. 2018 Jul 17;24(3):713–23.

15. Guglani L, Khader SA. Th17 cytokines in mucosal immunity and inflammation. Curr Opin HIV AIDS. 2010 Mar;5(2):120–7.

16. Kleinewietfeld M, Manzel A, Titze J, Kvakan H, Yosef N, Linker RA, et al. Sodium chloride drives autoimmune disease by the induction of pathogenic TH17 cells. Nature. 2013;496(7446):518–22.

17. Kleinewietfeld M, Hafler DA. The plasticity of human Treg and Th17 cells and its role in autoimmunity. Semin Immunol. 2013 Nov 15;25(4):305–12.

18. Romani L. Immunity to fungal infections. Nat Rev Immunol. 2011 Apr 11;11(4):275–88.

19. Bedoya SK, Lam B, Lau K, Larkin J, III. Th17 cells in immunity and autoimmunity. Clin Dev Immunol. 2013;2013:986789.

20. Di Cesare A, Di Meglio P, Nestle FO. The IL-23/Th17 Axis in the Immunopathogenesis of Psoriasis. J Invest Dermatol. 2009 Jun 1;129(6):1339–50.

21. Côme C, Cvrljevic A, Khan MM, Treise I, Adler T, Aguilar-Pimentel JA, et al. CIP2A promotes T-cell activation and immune response to Listeria monocytogenes infection. PLoS One. 2016;11(4):1–18.

22. Hoo LS, Zhang JY, Chan EK. Cloning and characterization of a novel 90 kDa ‘companion’ auto-antigen of p62 overexpressed in cancer. Oncogene. 2002 Jul;21(32):5006–15.

23. Choi H, Larsen B, Lin Z-Y, Breitkreutz A, Mellacheruvu D, Fermin D, et al. SAINT: probabilistic scoring of affinity purification-mass spectrometry data. Nat Methods. 2011 Jan;8(1):70–3.

24. Mellacheruvu D, Wright Z, Couzens AL, Lambert J-P, St-Denis NA, Li T, et al. The CRAPome: a contaminant repository for affinity purification–mass spectrometry data. Nat Methods. 2013 Aug 7;10(8):730–6.

25. Myant K, Qiao X, Halonen T, Come C, Laine A, Janghorban M, et al. Serine 62-Phosphorylated MYC Associates with Nuclear Lamins and Its Regulation by CIP2A Is Essential for Regenerative Proliferation. Cell Rep. 2015 Aug 11;12(6):1019–31.

26. Szklarczyk D, Morris JH, Cook H, Kuhn M, Wyder S, Simonovic M, et al. The STRING database in 2017: quality-controlled protein-protein association networks, made broadly accessible. Nucleic Acids Res. 2017 Jan 4;45(D1):D362–8.

27. Shannon P, Markiel A, Ozier O, Baliga NS, Wang JT, Ramage D, et al. Cytoscape: a software environment for integrated models of biomolecular interaction networks. Genome Res. 2003 Nov;13(11):2498–504.

28. Kauko O, Imanishi SY, Kulesskiy E, Laajala TD, Yetukuri L, Laine A, et al. Rules for PP2A-controlled phosphosignalling and drug responses. bioRxiv. 2018 Feb 26;271841.

29. Tripathi SK, Välikangas T, Shetty A, Khan MM, Moulder R, Bhosale SD, et al. Quantitative Proteomics Reveals the Dynamic Protein Landscape during Initiation of Human Th17 Cell Polarization. iScience. 2019 Jan 26;11:334–55.

30. Huang DW, Sherman BT, Lempicki RA. Systematic and integrative analysis of large gene lists using DAVID bioinformatics resources. Nat Protoc. 2009 Jan 1;4(1):44–57.

31. Huang DW, Sherman BT, Lempicki RA. Bioinformatics enrichment tools: paths toward the comprehensive functional analysis of large gene lists. Nucleic Acids Res. 2009 Jan;37(1):1–13.

32. Mi H, Huang X, Muruganujan A, Tang H, Mills C, Kang D, et al. PANTHER version 11: expanded annotation data from Gene Ontology and Reactome pathways, and data analysis tool enhancements. Nucleic Acids Res. 2017 Jan 4;45(D1):D183–9.

33. Mi H, Muruganujan A, Casagrande JT, Thomas PD. Large-scale gene function analysis with the PANTHER classification system. Nat Protoc. 2013 Jul 18;8(8):1551–66.

34. Shi Y. Serine/Threonine Phosphatases: Mechanism through Structure. Cell. 2009 Oct 30;139(3):468–84.

35. Nie H, Zheng Y, Li R, Guo TB, He D, Fang L, et al. Phosphorylation of FOXP3 controls regulatory T cell function and is inhibited by TNF-α in rheumatoid arthritis. Nat Med. 2013 Mar 10;19(3):322–8.

36. Lee J, Park E-J, Hwang JW, Oh J-M, Kim H, Bae E-K, et al. CIP2A expression is associated with synovial hyperplasia and invasive function of fibroblast-like synoviocytes in rheumatoid arthritis. Rheumatol Int. 2012 Jul 9;32(7):2023–30.

37. Lee J, Jeong H, Park E-J, Hwang JW, Huang B, Bae E-K, et al. CIP2A facilitates apoptotic resistance of fibroblast-like synoviocytes in rheumatoid arthritis independent of c-Myc expression. Rheumatol Int. 2013 Sep 2;33(9):2241–8.

38. Chow LML, Fournel M, Davidson D, Veillette A. Negative regulation of T-cell receptor signalling by tyrosine protein kinase p50 csk. Nature. 1993 Sep 9;365(6442):156–60.

39. Veillette A, Latour S, Davidson D. N EGATIVE R EGULATION OF I MMUNORECEPTOR S IGNALING. Annu Rev Immunol. 2002 Apr;20(1):669–707.

40. Zhang Q, Davis JJC, Lamborn IT, Freeman AF, Jing H, Favreau AJ, et al. Combined Immunodeficiency Associated with *DOCK8* Mutations. N Engl J Med. 2009 Nov 19;361(21):2046–55.

41. Su HC. Dedicator of cytokinesis 8 (DOCK8) deficiency. Curr Opin Allergy Clin Immunol. 2010 Dec;10(6):515–20.

42. Jabara HH, McDonald DR, Janssen E, Massaad MJ, Ramesh N, Borzutzky A, et al. DOCK8 functions as an adaptor that links TLR-MyD88 signaling to B cell activation. Nat Immunol. 2012 May 13;13(6):612–20.

43. Lazarevic V, Chen X, Shim J-H, Hwang E-S, Jang E, Bolm AN, et al. T-bet represses TH17 differentiation by preventing Runx1-mediated activation of the gene encoding RORγt. Nat Publ Gr. 2010;12.

44. Wang Y, Godec J, Ben-Aissa K, Cui K, Zhao K, Pucsek AB, et al. The transcription factors T-bet and runx are required for the ontogeny of pathogenic interferon-γ-producing T helper 17 cells. Immunity. 2014;40(3):355–66.

45. Espinosa A, Dardalhon V, Brauner S, Ambrosi A, Higgs R, Quintana FJ, et al. Loss of the lupus autoantigen Ro52/Trim21 induces tissue inflammation and systemic autoimmunity by disregulating the IL-23–Th17 pathway. J Exp Med. 2009 Aug 3;206(8):1661–71.

46. Yoshimi R, Ishigatsubo Y, Ozato K. Autoantigen TRIM21/Ro52 as a Possible Target for Treatment of Systemic Lupus Erythematosus. Int J Rheumatol. 2012;2012:718237.

47. Zhou G, Wu W, Yu L, Yu T, Yang W, Wang P, et al. Tripartite motif-containing (TRIM) 21 negatively regulates intestinal mucosal inflammation through inhibiting TH1/TH17 cell differentiation in patients with inflammatory bowel diseases. J Allergy Clin Immunol. 2017 Nov 4;

48. Machnicka B, Grochowalska R, Bogusławska DM, Sikorski AF, Lecomte MC. Spectrin-based skeleton as an actor in cell signaling. Cell Mol Life Sci. 2012 Jan;69(2):191–201.

49. Sihag RK, Cataldo AM. Brain beta-spectrin is a component of senile plaques in Alzheimer’s disease. Brain Res. 1996 Dec 16;743(1–2):249–57.

50. Yan X-X, Jeromin A, Jeromin A. Spectrin Breakdown Products (SBDPs) as Potential Biomarkers for Neurodegenerative Diseases. Curr Transl Geriatr Exp Gerontol Rep. 2012 Jun;1(2):85–93.

51. Burgess A, Vigneron S, Brioudes E, Labbé J-C, Lorca T, Castro A. Loss of human Greatwall results in G2 arrest and multiple mitotic defects due to deregulation of the cyclin B-Cdc2/PP2A balance. Proc Natl Acad Sci U S A. 2010 Jul 13;107(28):12564–9.

52. McCloy RA, Rogers S, Caldon CE, Lorca T, Castro A, Burgess A. Partial inhibition of Cdk1 in G _2_ phase overrides the SAC and decouples mitotic events. Cell Cycle. 2014 May 6;13(9):1400–12.

53. Carpenter AE, Jones TR, Lamprecht MR, Clarke C, Kang I, Friman O, et al. CellProfiler: image analysis software for identifying and quantifying cell phenotypes. Genome Biol. 2006;7(10):R100.

54. Vizcaíno JA, Csordas A, del-Toro N, Dianes JA, Griss J, Lavidas I, et al. 2016 update of the PRIDE database and its related tools. Nucleic Acids Res. 2016 Jan 4;44(D1):D447–56.

55. MacLean B, Tomazela DM, Shulman N, Chambers M, Finney GL, Frewen B, et al. Skyline: an open source document editor for creating and analyzing targeted proteomics experiments. Bioinformatics. 2010 Apr 1;26(7):966–8.

56. Farrah T, Deutsch EW, Kreisberg R, Sun Z, Campbell DS, Mendoza L, et al. PASSEL: the PeptideAtlas SRMexperiment library. Proteomics. 2012 Apr;12(8):1170–5.

57. Cox J, Neuhauser N, Michalski A, Scheltema RA, Olsen J V., Mann M. Andromeda: A Peptide Search Engine Integrated into the MaxQuant Environment. J Proteome Res. 2011 Apr 1;10(4):1794–805.

58. Cox J, Mann M. MaxQuant enables high peptide identification rates, individualized p.p.b.-range mass accuracies and proteome-wide protein quantification. Nat Biotechnol. 2008 Dec 30;26(12):1367–72.

59. Cox J, Hein MY, Luber CA, Paron I, Nagaraj N, Mann M. Accurate Proteome-wide Label-free Quantification by Delayed Normalization and Maximal Peptide Ratio Extraction, Termed MaxLFQ. Mol Cell Proteomics. 2014 Sep;13(9):2513–26.

60. Tyanova S, Temu T, Sinitcyn P, Carlson A, Hein MY, Geiger T, et al. The Perseus computational platform for comprehensive analysis of (prote)omics data. Nat Methods. 2016 Sep 27;13(9):731–40.

61. Choi M, Chang C-Y, Clough T, Broudy D, Killeen T, MacLean B, et al. MSstats: an R package for statistical analysis of quantitative mass spectrometry-based proteomic experiments. Bioinformatics. 2014 Sep 1;30(17):2524–6.

62. Morris JH, Apeltsin L, Newman AM, Baumbach J, Wittkop T, Su G, et al. clusterMaker: a multi-algorithm clustering plugin for Cytoscape. BMC Bioinformatics. 2011 Nov 9;12(1):436.

63. Brohée S, van Helden J. Evaluation of clustering algorithms for protein-protein interaction networks. BMC Bioinformatics. 2006 Nov 6;7(1):488.

64. Desiere F, Deutsch EW, King NL, Nesvizhskii AI, Mallick P, Eng J, et al. The PeptideAtlas project. Nucleic Acids Res. 2006 Jan 1;34(Database issue):D655–8.

