## Supplementary Table 1A for "Protein Interactome of the Cancerous Inhibitor of Protein Phosphatase 2A (CIP2A) in Th17 Cells"

### Supplementry Table 1A: Scoring from SAINT analysis

The data is filtered to only include proteins with a SP values of > 0.95 and less than 60% representation in the CRAPOME

The listed abundance and intensity values are from the SAINT processed MaxQuant intensity values

| Prey | Abundance | FCA | log.FCA. | SP | Prey Gene | Intensity | SAINT Intensity Sum | ctrlIntensity | Num Rep | Prob | iProb | AvgP | MaxP | FDR | % CRAPome |
|---|---|---|---|---|---|---|---|---|---|---|---|---|---|---|---|
| B0I1T2 | 21.7 | 0.78 | 0.83 | 1 | MYO1G | 6.307\|5.544\|6.255\|5.740 | 23.85 | 0.05\|.\|0.92\|0.14 | 4 | 1 | 1.00\|1.00\|1.00\|1.00 | 1.00 | 1 | 8.00E-04 | 3 |
| Q8TCG1 | 31.6 | 8.37 | 3.23 | 1 | KIAA1524 | 5.548\|6.011\|4.460\|4.254 | 20.27 | .\|.\|.\|. | 4 | 1 | 1.00\|1.00\|1.00\|1.00 | 1.00 | 0.9995 | 5.00E-04 | 2 |
| P11940 | 23.9 | 1.43 | 1.28 | 0.99 | PABPC1 | 3.495\|3.308\|3.450\|3.423 | 13.68 | .\|.\|0.90\|. | 4 | 0.99 | 0.99\|0.98\|0.99\|0.99 | 0.99 | 0.991 | 0.0048 | 58 |
| Q9NVI7 | 27.3 | 2.92 | 1.97 | 1 | ATAD3A | 3.513\|3.161\|3.418\|3.164 | 13.26 | .\|.\|0.06\|. | 4 | 1 | 1.00\|1.00\|1.00\|1.00 | 1.00 | 1 | 1.00E-04 | 29 |
| Q9NVJ2 | 21.5 | 6.03 | 2.81 | 0.96 | ARL8B | 0.753\|0.785\|0.692\|0.698 | 2.93 | .\|.\|.\|. | 4 | 0.96 | 0.97\|0.97\|0.97\|0.96 | 0.96 | 0.9685 | 0.0087 | 2 |
| P19474 | 28.8 | 7.73 | 3.13 | 1 | TRIM21 | 3.236\|4.121\|2.318\|2.449 | 12.13 | .\|.\|.\|. | 4 | 1 | 1.00\|1.00\|1.00\|1.00 | 1.00 | 0.998 | 0.0014 | 54 |
| P35580 | 27.1 | 3.12 | 2.04 | 1 | MYH10 | 3.116\|3.029\|3.063\|2.960 | 12.17 | .\|.\|0.05\|. | 4 | 1 | 1.00\|1.00\|1.00\|1.00 | 1.00 | 0.9995 | 7.00E-04 | 52 |
| Q01082 | 28.9 | 7.75 | 3.13 | 1 | SPTBN1 | 2.930\|2.846\|2.942\|3.293 | 12.01 | .\|.\|.\|. | 4 | 1 | 1.00\|1.00\|1.00\|1.00 | 1.00 | 1 | 8.00E-04 | 49 |
| P07910 | 13.4 | 0.47 | 0.56 | 0.99 | HNRNPC | 2.894\|2.626\|3.166\|3.149 | 11.83 | 0.12\|0.12\|0.48\|0.40 | 4 | 1 | 1.00\|1.00\|0.99\|0.99 | 0.99 | 0.998 | 0.0025 | 59 |
| Q8NF50 | 26.7 | 2.30 | 1.72 | 1 | DOCK8 | 3.086\|2.856\|2.961\|2.922 | 11.83 | .\|0.06\|.\|. | 4 | 1 | 1.00\|1.00\|1.00\|1.00 | 1.00 | 1 | 4.00E-04 | 1 |
| Q15149 | 26.7 | 2.40 | 1.77 | 1 | PLEC | 3.119\|2.650\|3.053\|2.932 | 11.76 | .\|.\|.\|0.06 | 4 | 1 | 1.00\|1.00\|1.00\|1.00 | 1.00 | 1 | 8.00E-04 | 32 |
| Q9UKV3 | 28.6 | 7.68 | 3.12 | 1 | ACIN1 | 2.807\|2.679\|2.843\|2.965 | 11.29 | .\|.\|.\|. | 4 | 1 | 1.00\|1.00\|1.00\|1.00 | 1.00 | 1 | 4.00E-04 | 13 |
| Q9NYL9 | 28.4 | 7.64 | 3.11 | 1 | TMOD3 | 2.772\|2.827\|2.768\|2.647 | 11.01 | .\|.\|.\|. | 4 | 1 | 1.00\|1.00\|1.00\|1.00 | 1.00 | 1 | 4.00E-04 | 16 |
| P0DN76 | 22.4 | 2.58 | 1.84 | 1 | U2AF1L5 | 1.237\|1.195\|1.284\|1.302 | 5.02 | .\|.\|0.05\|. | 4 | 1 | 1.00\|1.00\|0.99\|1.00 | 1.00 | 0.9965 | 0.002 | 40 |
| Q6NYC8 | 28.1 | 7.57 | 3.1 | 1 | PPP1R18 | 2.542\|2.465\|2.530\|2.850 | 10.39 | .\|.\|.\|. | 4 | 1 | 1.00\|1.00\|1.00\|1.00 | 1.00 | 0.999 | 0.0013 | 2 |
| Q9H307 | 28.1 | 7.56 | 3.1 | 1 | PNN | 2.599\|2.437\|2.594\|2.655 | 10.29 | .\|.\|.\|. | 4 | 1 | 1.00\|1.00\|1.00\|1.00 | 1.00 | 1 | 4.00E-04 | 10 |
| Q7Z2W4 | 27.9 | 7.52 | 3.09 | 1 | ZC3HAV1 | 2.627\|3.011\|2.132\|2.248 | 10.02 | .\|.\|.\|. | 4 | 1 | 1.00\|1.00\|1.00\|1.00 | 1.00 | 0.9995 | 6.00E-04 | 14 |
| Q13428 | 28 | 7.55 | 3.1 | 1 | TCOF1 | 2.700\|2.373\|2.720\|2.412 | 10.21 | .\|.\|.\|. | 4 | 1 | 1.00\|1.00\|1.00\|1.00 | 1.00 | 0.9995 | 0.0011 | 34 |
| P52907 | 24.4 | 1.91 | 1.54 | 1 | CAPZA1 | 2.414\|2.556\|2.595\|2.647 | 10.21 | .\|.\|0.20\|. | 4 | 1 | 1.00\|1.00\|1.00\|1.00 | 1.00 | 1 | 3.00E-04 | 42 |
| Q7Z406 | 22.2 | 1.23 | 1.16 | 1 | MYH14 | 2.309\|2.656\|2.450\|2.580 | 10.00 | .\|.\|0.21\|0.05 | 4 | 1 | 1.00\|1.00\|1.00\|1.00 | 1.00 | 1 | 4.00E-04 | 40 |
| O43795 | 27.8 | 7.51 | 3.09 | 1 | MYO1B | 2.428\|2.436\|2.468\|2.491 | 9.82 | .\|.\|.\|. | 4 | 1 | 1.00\|1.00\|1.00\|1.00 | 1.00 | 0.9995 | 6.00E-04 | 14 |
| P38919 | 27.8 | 7.49 | 3.09 | 1 | EIF4A3 | 2.485\|2.281\|2.458\|2.498 | 9.72 | .\|.\|.\|. | 4 | 1 | 1.00\|1.00\|1.00\|1.00 | 1.00 | 1 | 4.00E-04 | 49 |
| O00159 | 25.9 | 2.49 | 1.8 | 1 | MYO1C | 2.385\|2.559\|2.345\|2.388 | 9.68 | .\|.\|.\|0.05 | 4 | 1 | 1.00\|1.00\|1.00\|1.00 | 1.00 | 0.9995 | 6.00E-04 | 20 |
| Q9Y2W1 | 26.2 | 2.86 | 1.95 | 1 | THRAP3 | 2.518\|2.406\|2.410\|2.246 | 9.58 | 0.04\|.\|.\|. | 4 | 1 | 1.00\|1.00\|1.00\|1.00 | 1.00 | 1 | 3.00E-04 | 51 |
| Q92614 | 27.7 | 7.47 | 3.08 | 1 | MYO18A | 2.401\|2.375\|2.417\|2.350 | 9.54 | .\|.\|.\|. | 4 | 1 | 1.00\|1.00\|1.00\|1.00 | 1.00 | 0.999 | 0.001 | 2 |
| P62136 | 27.5 | 7.44 | 3.08 | 1 | PPP1CA | 2.240\|2.240\|2.274\|2.526 | 9.28 | .\|.\|.\|. | 4 | 1 | 1.00\|1.00\|1.00\|1.00 | 1.00 | 0.999 | 9.00E-04 | 40 |

| | | | | | | | | | | | | | | | |
|---|---|---|---|---|---|---|---|---|---|---|---|---|---|---|---|
| P14866 | 27.5 | 7.43 | 3.08 | 0.99 | HNRNPL | 2.266\|2.169\|2.433\|2.371 | 9.24 | .\|.\|.\|. | 4 | 0.99 | 0.98\|0.98\|0.99\|0.99 | 0.99 | 0.987 | 0.0058 | 58 |
| P47756 | 27.5 | 7.42 | 3.07 | 1 | CAPZB | 2.235\|2.239\|2.332\|2.338 | 9.14 | .\|.\|.\|. | 4 | 1 | 1.00\|1.00\|1.00\|1.00 | 1.00 | 1 | 6.00E-04 | 37 |
| Q9NYF8 | 27.4 | 7.41 | 3.07 | 1 | BCLAF1 | 2.378\|2.271\|2.243\|2.204 | 9.10 | .\|.\|.\|. | 4 | 1 | 1.00\|1.00\|1.00\|1.00 | 1.00 | 0.9995 | 6.00E-04 | 48 |
| O14974 | 27.3 | 7.37 | 3.06 | 1 | PPP1R12A | 2.146\|2.172\|2.156\|2.295 | 8.77 | .\|.\|.\|. | 4 | 1 | 1.00\|1.00\|1.00\|1.00 | 1.00 | 0.9995 | 7.00E-04 | 10 |
| Q6P2E9 | 26.8 | 7.27 | 3.05 | 1 | EDC4 | 1.616\|1.648\|2.402\|2.603 | 8.27 | .\|.\|.\|. | 4 | 1 | 1.00\|1.00\|1.00\|1.00 | 1.00 | 0.9985 | 0.0012 | 13 |
| P12268 | 24.6 | 2.93 | 1.98 | 1 | IMPDH2 | 2.311\|2.719\|1.515\|1.299 | 7.85 | .\|.\|0.05\|. | 4 | 1 | 1.00\|1.00\|1.00\|0.99 | 1.00 | 0.9995 | 0.0019 | 32 |
| Q5T9A4 | 27.1 | 7.34 | 3.06 | 1 | ATAD3B | 2.265\|1.973\|2.216\|2.134 | 8.59 | .\|.\|.\|. | 4 | 1 | 1.00\|1.00\|1.00\|1.00 | 1.00 | 0.9995 | 8.00E-04 | 23 |
| Q96HS1 | 24.8 | 2.15 | 1.65 | 1 | PGAM5 | 2.088\|1.921\|2.158\|2.193 | 8.36 | .\|0.06\|.\|. | 4 | 1 | 1.00\|1.00\|1.00\|1.00 | 1.00 | 0.9995 | 5.00E-04 | 26 |
| P62906 | 25.4 | 2.74 | 1.9 | 1 | RPL10A | 2.109\|2.157\|2.027\|2.007 | 8.30 | 0.04\|.\|.\|. | 4 | 1 | 1.00\|1.00\|1.00\|1.00 | 1.00 | 1 | 0.0012 | 43 |
| O00422 | 23.4 | 1.69 | 1.43 | 1 | SAP18 | 2.097\|1.963\|2.135\|2.103 | 8.30 | 0.04\|.\|0.05\|. | 4 | 1 | 1.00\|1.00\|1.00\|1.00 | 1.00 | 1 | 4.00E-04 | 14 |
| Q02878 | 11.3 | 0.43 | 0.52 | 0.99 | RPL6 | 2.355\|2.071\|2.082\|1.624 | 8.13 | 0.38\|0.38\|0.17\|0.12 | 4 | 1 | 0.99\|1.00\|0.99\|0.99 | 0.99 | 0.998 | 0.0025 | 57 |
| Q9Y6N5 | 26.9 | 7.29 | 3.05 | 1 | SQOR | 2.175\|2.136\|2.001\|1.965 | 8.28 | .\|.\|.\|. | 4 | 1 | 1.00\|1.00\|1.00\|1.00 | 1.00 | 0.999 | 0.001 | 1 |
| P43243 | 25.2 | 2.56 | 1.83 | 1 | MATR3 | 2.139\|1.852\|2.105\|2.138 | 8.23 | 0.04\|.\|.\|. | 4 | 1 | 1.00\|1.00\|1.00\|1.00 | 1.00 | 0.9995 | 8.00E-04 | 55 |
| P04114 | 24.9 | 2.76 | 1.91 | 1 | APOB | 2.147\|2.063\|1.977\|2.048 | 8.24 | .\|.\|0.06\|. | 4 | 1 | 1.00\|1.00\|1.00\|1.00 | 1.00 | 1 | 3.00E-04 | 1 |
| P84103 | 26.9 | 7.29 | 3.05 | 1 | SRSF3 | 2.067\|1.969\|2.040\|2.142 | 8.22 | .\|.\|.\|. | 4 | 1 | 1.00\|1.00\|1.00\|1.00 | 1.00 | 1 | 0.001 | 57 |
| Q92804 | 26.9 | 7.28 | 3.05 | 1 | TAF15 | 1.938\|1.964\|2.229\|2.031 | 8.16 | .\|.\|.\|. | 4 | 1 | 1.00\|1.00\|1.00\|1.00 | 1.00 | 0.9985 | 0.0014 | 56 |
| P50914 | 20.4 | 1.12 | 1.08 | 0.97 | RPL14 | 2.242\|2.253\|1.760\|1.722 | 7.98 | .\|.\|0.51\|0.04 | 4 | 0.98 | 0.99\|0.99\|0.96\|0.95 | 0.97 | 0.994 | 0.0083 | 58 |
| P62834 | 23.5 | 6.48 | 2.9 | 1 | RAP1A | 1.121\|1.133\|1.011\|0.996 | 4.26 | .\|.\|.\|. | 4 | 0.99 | 1.00\|1.00\|0.99\|0.99 | 1.00 | 0.997 | 0.0023 | 11 |
| Q15154 | 26.5 | 7.20 | 3.04 | 1 | PCM1 | 1.573\|1.644\|2.197\|2.304 | 7.72 | .\|.\|.\|. | 4 | 1 | 0.99\|1.00\|1.00\|1.00 | 1.00 | 0.9995 | 0.0014 | 4 |
| P27694 | 26.7 | 7.24 | 3.04 | 1 | RPA1 | 1.987\|2.221\|1.812\|1.880 | 7.90 | .\|.\|.\|. | 4 | 1 | 1.00\|1.00\|1.00\|1.00 | 1.00 | 0.999 | 9.00E-04 | 30 |
| Q16666 | 26.7 | 7.24 | 3.04 | 1 | IFI16 | 2.037\|2.049\|1.896\|1.899 | 7.88 | .\|.\|.\|. | 4 | 1 | 1.00\|1.00\|1.00\|1.00 | 1.00 | 0.999 | 9.00E-04 | 2 |
| P06753 | 21.3 | 1.25 | 1.17 | 1 | TPM3 | 1.880\|2.158\|1.813\|1.960 | 7.81 | 0.05\|.\|0.15\|. | 4 | 1 | 1.00\|1.00\|1.00\|1.00 | 1.00 | 1 | 8.00E-04 | 50 |
| O75643 | 26.6 | 7.21 | 3.04 | 1 | SNRNP200 | 1.761\|1.716\|2.134\|2.134 | 7.74 | .\|.\|.\|. | 4 | 1 | 1.00\|1.00\|1.00\|1.00 | 1.00 | 0.9995 | 0.0012 | 41 |
| P26599 | 20.5 | 1.11 | 1.08 | 0.98 | PTBP1 | 1.903\|1.777\|2.053\|2.068 | 7.80 | .\|0.05\|0.31\|. | 4 | 0.98 | 0.99\|0.99\|0.97\|0.97 | 0.98 | 0.9935 | 0.0076 | 55 |
| P35244 | 26.4 | 7.15 | 3.03 | 1 | RPA3 | 1.945\|2.289\|1.629\|1.597 | 7.46 | .\|.\|.\|. | 4 | 1 | 1.00\|1.00\|1.00\|1.00 | 1.00 | 0.9995 | 9.00E-04 | 15 |
| Q9UHB6 | 26.5 | 7.20 | 3.04 | 1 | LIMA1 | 1.901\|1.832\|1.896\|2.032 | 7.66 | .\|.\|.\|. | 4 | 1 | 1.00\|1.00\|1.00\|1.00 | 1.00 | 0.999 | 0.001 | 15 |
| P51114 | 26.4 | 7.17 | 3.03 | 1 | FXR1 | 2.064\|2.078\|1.669\|1.697 | 7.51 | .\|.\|.\|. | 4 | 1 | 1.00\|1.00\|1.00\|1.00 | 1.00 | 0.999 | 0.0011 | 13 |
| Q00325 | 23 | 1.67 | 1.42 | 1 | SLC25A3 | 2.012\|2.065\|1.773\|1.686 | 7.54 | .\|0.04\|0.04\|. | 4 | 1 | 1.00\|1.00\|1.00\|1.00 | 1.00 | 0.999 | 0.0012 | 37 |
| Q9Y5S9 | 26.5 | 7.19 | 3.03 | 1 | RBM8A | 1.824\|1.810\|1.938\|2.010 | 7.58 | .\|.\|.\|. | 4 | 1 | 1.00\|1.00\|1.00\|1.00 | 1.00 | 0.9995 | 0.001 | 23 |
| Q02543 | 26.5 | 7.18 | 3.03 | 1 | RPL18A | 1.934\|1.976\|1.863\|1.771 | 7.54 | .\|.\|.\|. | 4 | 1 | 1.00\|1.00\|1.00\|1.00 | 1.00 | 0.997 | 0.0018 | 40 |
| Q09028 | 22.6 | 6.28 | 2.86 | 0.98 | RBBP4 | 0.944\|0.931\|0.887\|0.848 | 3.61 | .\|.\|.\|. | 4 | 0.98 | 0.98\|0.98\|0.98\|0.97 | 0.98 | 0.982 | 0.0077 | 58 |
| P61353 | 20.9 | 1.11 | 1.08 | 1 | RPL27 | 1.988\|1.827\|1.856\|1.743 | 7.41 | .\|0.05\|0.05\|0.05 | 4 | 1 | 1.00\|1.00\|1.00\|1.00 | 1.00 | 0.999 | 0.001 | 55 |
| P46940 | 24.3 | 2.21 | 1.68 | 1 | IQGAP1 | 1.846\|1.776\|1.814\|1.931 | 7.37 | 0.06\|.\|.\|. | 4 | 1 | 1.00\|1.00\|1.00\|1.00 | 1.00 | 0.999 | 0.0012 | 28 |

| | | | | | | | | | | | | | | | |
|---|---|---|---|---|---|---|---|---|---|---|---|---|---|---|---|
| O43143 | 26.3 | 7.15 | 3.03 | 1 | DHX15 | 1.800\|1.802\|1.864\|1.852 | 7.32 | .\|.\|.\|. | 4 | 1 | 1.00\|1.00\|1.00\|1.00 | 1.00 | 0.9975 | 0.0017 | 54 |
| Q6P2Q9 | 20.7 | 1.10 | 1.07 | 1 | PRPF8 | 1.592\|1.474\|2.010\|2.022 | 7.10 | .\|0.05\|0.05\|0.04 | 4 | 1 | 1.00\|1.00\|1.00\|1.00 | 1.00 | 0.9995 | 0.0013 | 43 |
| P04439 | 21.6 | 1.47 | 1.31 | 1 | HLA-A | 1.632\|1.537\|1.598\|1.506 | 6.27 | .\|0.05\|0.06\|. | 4 | 1 | 1.00\|1.00\|1.00\|1.00 | 1.00 | 0.998 | 0.0015 | 7 |
| P06396 | 25.4 | 6.93 | 2.99 | 1 | GSN | 2.080\|2.112\|1.285\|1.013 | 6.49 | .\|.\|.\|. | 4 | 1 | 1.00\|1.00\|1.00\|0.99 | 1.00 | 1 | 0.0018 | 9 |
| Q9UMS4 | 26 | 7.09 | 3.02 | 1 | PRPF19 | 1.626\|1.561\|1.855\|1.942 | 6.99 | .\|.\|.\|. | 4 | 1 | 0.99\|0.99\|1.00\|1.00 | 1.00 | 0.9975 | 0.0023 | 39 |
| P04899 | 24.2 | 2.81 | 1.93 | 1 | GNAI2 | 1.877\|1.689\|1.744\|1.702 | 7.01 | .\|.\|0.05\|. | 4 | 0.99 | 1.00\|0.99\|1.00\|1.00 | 1.00 | 0.9975 | 0.0019 | 15 |
| Q9UQ35 | 26.1 | 7.09 | 3.02 | 1 | SRRM2 | 1.732\|1.680\|1.750\|1.810 | 6.97 | .\|.\|.\|. | 4 | 1 | 1.00\|1.00\|1.00\|1.00 | 1.00 | 0.999 | 0.0013 | 48 |
| Q86U42 | 26 | 7.07 | 3.01 | 1 | PABPN1 | 1.654\|1.505\|1.812\|1.925 | 6.90 | .\|.\|.\|. | 4 | 1 | 1.00\|1.00\|1.00\|1.00 | 1.00 | 0.9985 | 0.0012 | 36 |
| P14649 | 26 | 7.07 | 3.01 | 1 | MYL6B | 1.713\|1.819\|1.679\|1.649 | 6.86 | .\|.\|.\|. | 4 | 1 | 1.00\|1.00\|1.00\|1.00 | 1.00 | 1 | 6.00E-04 | 47 |
| Q13310 | 25.9 | 7.06 | 3.01 | 1 | PABPC4 | 1.729\|1.650\|1.690\|1.736 | 6.81 | .\|.\|.\|. | 4 | 1 | 1.00\|1.00\|1.00\|1.00 | 1.00 | 0.9975 | 0.0016 | 50 |
| Q15029 | 25.9 | 7.05 | 3.01 | 1 | EFTUD2 | 1.622\|1.570\|1.803\|1.767 | 6.76 | .\|.\|.\|. | 4 | 1 | 1.00\|1.00\|0.99\|1.00 | 1.00 | 0.999 | 0.0016 | 49 |
| O43707 | 25.9 | 7.05 | 3.01 | 0.99 | ACTN4 | 1.667\|1.662\|1.630\|1.789 | 6.75 | .\|.\|.\|. | 4 | 1 | 0.99\|0.99\|0.99\|1.00 | 0.99 | 0.998 | 0.0025 | 46 |
| P42224 | 23.8 | 2.19 | 1.67 | 1 | STAT1 | 1.681\|1.881\|1.471\|1.557 | 6.59 | .\|0.05\|.\|. | 4 | 1 | 1.00\|1.00\|1.00\|1.00 | 1.00 | 0.999 | 0.0015 | 2 |
| P62995 | 25.8 | 7.03 | 3 | 1 | TRA2B | 1.681\|1.584\|1.716\|1.657 | 6.64 | .\|.\|.\|. | 4 | 1 | 1.00\|0.99\|1.00\|1.00 | 1.00 | 0.998 | 0.0015 | 22 |
| Q15366 | 25.8 | 7.02 | 3 | 1 | PCBP2 | 1.684\|1.571\|1.715\|1.621 | 6.59 | .\|.\|.\|. | 4 | 1 | 1.00\|1.00\|1.00\|1.00 | 1.00 | 0.999 | 0.0013 | 57 |
| P26196 | 25.6 | 6.99 | 3 | 1 | DDX6 | 1.486\|1.370\|1.811\|1.773 | 6.44 | .\|.\|.\|. | 4 | 1 | 0.99\|1.00\|1.00\|1.00 | 1.00 | 0.9995 | 0.0016 | 39 |
| Q15393 | 21.7 | 1.46 | 1.3 | 1 | SF3B3 | 1.565\|1.436\|1.721\|1.743 | 6.47 | .\|0.05\|0.05\|. | 4 | 1 | 1.00\|1.00\|1.00\|1.00 | 1.00 | 0.9995 | 5.00E-04 | 51 |
| Q96II8 | 25.7 | 7.00 | 3 | 1 | LRCH3 | 1.680\|1.552\|1.680\|1.587 | 6.50 | .\|.\|.\|. | 4 | 1 | 1.00\|1.00\|1.00\|1.00 | 1.00 | 0.999 | 0.0012 | 0 |
| Q15717 | 25.6 | 6.99 | 3 | 0.99 | ELAVL1 | 1.595\|1.486\|1.663\|1.715 | 6.46 | .\|.\|.\|. | 4 | 0.99 | 0.99\|0.99\|0.99\|1.00 | 0.99 | 0.9955 | 0.003 | 38 |
| P35611 | 25.7 | 6.99 | 3 | 1 | ADD1 | 1.617\|1.484\|1.701\|1.655 | 6.46 | .\|.\|.\|. | 4 | 1 | 1.00\|1.00\|1.00\|1.00 | 1.00 | 0.998 | 0.0016 | 9 |
| P42167 | 25.6 | 6.99 | 3 | 0.99 | TMPO | 1.690\|1.697\|1.509\|1.536 | 6.43 | .\|.\|.\|. | 4 | 0.99 | 0.99\|1.00\|0.99\|0.99 | 0.99 | 0.995 | 0.0035 | 44 |
| P62873 | 25.6 | 6.98 | 3 | 1 | GNB1 | 1.746\|1.705\|1.469\|1.482 | 6.40 | .\|.\|.\|. | 4 | 1 | 1.00\|1.00\|1.00\|1.00 | 1.00 | 0.998 | 0.0015 | 16 |
| Q15306 | 25.6 | 6.99 | 3 | 0.99 | IRF4 | 1.572\|1.564\|1.656\|1.629 | 6.42 | .\|.\|.\|. | 4 | 0.99 | 0.99\|0.99\|1.00\|0.99 | 0.993 | 0.9965 | 0.0031 | 0 |
| P27708 | 25.6 | 6.98 | 3 | 1 | CAD | 1.681\|1.652\|1.562\|1.500 | 6.39 | .\|.\|.\|. | 4 | 1 | 1.00\|1.00\|1.00\|1.00 | 1.00 | 0.9985 | 0.0015 | 49 |
| P15927 | 25.6 | 6.97 | 2.99 | 1 | RPA2 | 1.602\|1.786\|1.474\|1.495 | 6.36 | .\|.\|.\|. | 4 | 0.99 | 1.00\|1.00\|0.99\|1.00 | 1.00 | 0.997 | 0.0021 | 20 |
| O94832 | 25.6 | 6.97 | 2.99 | 0.99 | MYO1D | 1.667\|1.491\|1.704\|1.492 | 6.35 | .\|.\|.\|. | 4 | 0.99 | 1.00\|0.99\|1.00\|0.99 | 0.99 | 0.995 | 0.0032 | 9 |
| P63244 | 25.6 | 6.98 | 3 | 1 | RACK1 | 1.655\|1.633\|1.552\|1.529 | 6.37 | .\|.\|.\|. | 4 | 1 | 1.00\|1.00\|1.00\|1.00 | 1.00 | 0.998 | 0.0014 | 43 |
| P47755 | 25.6 | 6.98 | 3 | 1 | CAPZA2 | 1.536\|1.525\|1.636\|1.666 | 6.36 | .\|.\|.\|. | 4 | 1 | 1.00\|0.99\|1.00\|1.00 | 1.00 | 0.9965 | 0.0022 | 27 |
| O75494 | 23.5 | 2.17 | 1.66 | 1 | SRSF10 | 1.565\|1.515\|1.617\|1.575 | 6.27 | .\|.\|.\|0.06 | 4 | 1 | 1.00\|1.00\|1.00\|1.00 | 1.00 | 0.997 | 0.0019 | 17 |
| P62140 | 25.5 | 6.96 | 2.99 | 0.99 | PPP1CB | 1.550\|1.532\|1.519\|1.657 | 6.26 | .\|.\|.\|. | 4 | 0.99 | 0.99\|0.99\|0.99\|0.99 | 0.99 | 0.9945 | 0.0035 | 40 |
| O75531 | 25.4 | 6.92 | 2.99 | 1 | BANF1 | 1.696\|1.752\|1.347\|1.347 | 6.14 | .\|.\|.\|. | 4 | 1 | 1.00\|1.00\|1.00\|1.00 | 1.00 | 1 | 8.00E-04 | 22 |
| Q9Y224 | 25.5 | 6.95 | 2.99 | 1 | C14ORF166 | 1.608\|1.540\|1.574\|1.527 | 6.25 | .\|.\|.\|. | 4 | 1 | 1.00\|1.00\|1.00\|0.99 | 1.00 | 0.997 | 0.002 | 36 |
| Q8NI27 | 25.5 | 6.95 | 2.99 | 1 | THOC2 | 1.568\|1.451\|1.608\|1.602 | 6.23 | .\|.\|.\|. | 4 | 1 | 1.00\|1.00\|1.00\|1.00 | 1.00 | 0.998 | 0.0014 | 18 |

| | | | | | | | | | | | | | | | |
|---|---|---|---|---|---|---|---|---|---|---|---|---|---|---|---|
| O00299 | 25.4 | 6.94 | 2.99 | 1 | CLIC1 | 1.504\|1.561\|1.536\|1.596 | 6.20 | .\|.\|.\|. | 4 | 1 | 1.00\|1.00\|1.00\|1.00 | 1.00 | 0.999 | 0.0013 | 16 |
| P49756 | 25.4 | 6.94 | 2.99 | 1 | RBM25 | 1.574\|1.491\|1.503\|1.592 | 6.16 | .\|.\|.\|. | 4 | 1 | 0.99\|1.00\|1.00\|1.00 | 1.00 | 0.998 | 0.0017 | 22 |
| P60891 | 24.8 | 6.79 | 2.96 | 1 | PRPS1 | 1.296\|1.297\|1.378\|1.480 | 5.45 | .\|.\|.\|. | 4 | 1 | 1.00\|1.00\|1.00\|1.00 | 1.00 | 0.9985 | 0.0017 | 33 |
| Q9H9B4 | 25.4 | 6.92 | 2.99 | 1 | SFXN1 | 1.599\|1.568\|1.450\|1.477 | 6.09 | .\|.\|.\|. | 4 | 1 | 1.00\|1.00\|1.00\|0.99 | 1.00 | 0.9965 | 0.002 | 6 |
| O14979 | 23.4 | 2.69 | 1.88 | 1 | HNRNPDL | 1.543\|1.389\|1.593\|1.560 | 6.09 | .\|.\|0.05\|. | 4 | 1 | 1.00\|1.00\|1.00\|1.00 | 1.00 | 0.999 | 0.0014 | 48 |
| P15153 | 23.3 | 2.66 | 1.87 | 0.99 | RAC2 | 1.535\|1.626\|1.421\|1.437 | 6.02 | .\|.\|0.05\|. | 4 | 0.99 | 1.00\|0.99\|0.99\|0.99 | 0.99 | 0.995 | 0.0036 | 6 |
| Q86W42 | 25.3 | 6.91 | 2.98 | 0.99 | THOC6 | 1.551\|1.435\|1.551\|1.496 | 6.03 | .\|.\|.\|. | 4 | 0.99 | 0.99\|0.99\|0.99\|0.99 | 0.99 | 0.9915 | 0.0044 | 11 |
| P49327 | 25.3 | 6.90 | 2.98 | 1 | FASN | 1.540\|1.513\|1.501\|1.436 | 5.99 | .\|.\|.\|. | 4 | 1 | 1.00\|1.00\|0.99\|1.00 | 1.00 | 0.997 | 0.002 | 52 |
| Q13427 | 25.2 | 6.90 | 2.98 | 0.99 | PPIG | 1.529\|1.447\|1.468\|1.514 | 5.96 | .\|.\|.\|. | 4 | 0.99 | 0.99\|0.99\|0.99\|0.99 | 0.99 | 0.994 | 0.0033 | 10 |
| O14639 | 25.2 | 6.88 | 2.98 | 0.99 | ABLIM1 | 1.482\|1.540\|1.330\|1.538 | 5.89 | .\|.\|.\|. | 4 | 0.99 | 1.00\|1.00\|0.99\|1.00 | 0.99 | 0.9965 | 0.0024 | 6 |
| Q9Y3I0 | 25.1 | 6.86 | 2.98 | 1 | RTCB | 1.500\|1.434\|1.451\|1.416 | 5.80 | .\|.\|.\|. | 4 | 0.99 | 1.00\|1.00\|0.99\|1.00 | 1.00 | 0.998 | 0.0022 | 44 |
| O75400 | 22.9 | 2.10 | 1.63 | 1 | PRPF40A | 1.465\|1.396\|1.422\|1.462 | 5.75 | 0.06\|.\|.\|. | 4 | 1 | 1.00\|1.00\|1.00\|0.99 | 1.00 | 0.9975 | 0.0019 | 21 |
| P29350 | 23 | 2.66 | 1.87 | 1 | PTPN6 | 1.490\|1.475\|1.384\|1.326 | 5.68 | .\|.\|0.05\|. | 4 | 1 | 1.00\|1.00\|1.00\|1.00 | 1.00 | 0.9975 | 0.0016 | 0 |
| Q8WWM7 | 25 | 6.83 | 2.97 | 1 | ATXN2L | 1.461\|1.445\|1.412\|1.333 | 5.65 | .\|.\|.\|. | 4 | 1 | 1.00\|1.00\|1.00\|0.99 | 1.00 | 0.9965 | 0.0022 | 38 |
| P55209 | 24.9 | 6.82 | 2.97 | 0.99 | NAP1L1 | 1.507\|1.396\|1.404\|1.286 | 5.59 | .\|.\|.\|. | 4 | 0.99 | 1.00\|0.99\|1.00\|0.99 | 0.99 | 0.9965 | 0.0025 | 50 |
| Q9Y3U8 | 21.4 | 1.58 | 1.37 | 1 | RPL36 | 1.390\|1.534\|1.346\|1.312 | 5.58 | .\|.\|0.05\|0.04 | 4 | 0.99 | 1.00\|1.00\|1.00\|1.00 | 1.00 | 0.998 | 0.0015 | 39 |
| Q6WKZ4 | 24.8 | 6.79 | 2.96 | 0.99 | RAB11FIP1 | 1.478\|1.151\|1.555\|1.318 | 5.50 | .\|.\|.\|. | 4 | 0.99 | 0.99\|0.99\|0.99\|0.99 | 0.99 | 0.9945 | 0.0029 | 1 |
| P06239 | 24.9 | 6.81 | 2.97 | 0.99 | LCK | 1.458\|1.358\|1.398\|1.351 | 5.57 | .\|.\|.\|. | 4 | 0.99 | 0.99\|0.99\|1.00\|0.99 | 0.99 | 0.995 | 0.0028 | 13 |
| Q9Y4I1 | 24.9 | 6.81 | 2.97 | 0.99 | MYO5A | 1.349\|1.433\|1.446\|1.324 | 5.55 | .\|.\|.\|. | 4 | 1 | 0.99\|0.99\|1.00\|1.00 | 0.99 | 0.996 | 0.0025 | 6 |
| Q86V48 | 14.2 | 0.60 | 0.68 | 0.99 | LUZP1 | 1.321\|1.319\|1.389\|1.504 | 5.53 | 0.31\|0.06\|0.30\|. | 4 | 0.99 | 1.00\|0.99\|0.99\|1.00 | 0.99 | 0.9955 | 0.0025 | 3 |
| Q92499 | 24.8 | 6.80 | 2.96 | 0.99 | DDX1 | 1.444\|1.382\|1.337\|1.340 | 5.50 | .\|.\|.\|. | 4 | 0.99 | 0.99\|0.99\|0.99\|0.99 | 0.99 | 0.9915 | 0.0041 | 46 |
| P63167 | 23.4 | 6.48 | 2.9 | 0.99 | DYNLL1 | 1.071\|1.049\|1.058\|1.047 | 4.23 | .\|.\|.\|. | 4 | 0.99 | 0.98\|0.98\|0.99\|0.99 | 0.99 | 0.991 | 0.0059 | 19 |
| Q12906 | 24.8 | 6.80 | 2.96 | 0.99 | ILF3 | 1.345\|1.283\|1.406\|1.456 | 5.49 | .\|.\|.\|. | 4 | 1 | 0.99\|0.99\|0.99\|0.99 | 0.99 | 0.9945 | 0.0033 | 52 |
| P05023 | 20.5 | 1.37 | 1.25 | 1 | ATP1A1 | 1.388\|1.478\|1.282\|1.253 | 5.40 | 0.06\|.\|0.06\|. | 4 | 0.99 | 1.00\|1.00\|1.00\|0.99 | 1.00 | 0.997 | 0.0018 | 36 |
| Q9NR30 | 24.7 | 6.77 | 2.96 | 1 | DDX21 | 1.464\|1.343\|1.336\|1.248 | 5.39 | .\|.\|.\|. | 4 | 1 | 1.00\|0.99\|1.00\|1.00 | 1.00 | 0.997 | 0.0023 | 54 |
| Q9UEY8 | 24.7 | 6.77 | 2.96 | 0.99 | ADD3 | 1.361\|1.205\|1.432\|1.385 | 5.38 | .\|.\|.\|. | 4 | 1 | 0.99\|0.99\|1.00\|0.99 | 0.99 | 0.995 | 0.0029 | 5 |
| Q14764 | 24.6 | 6.76 | 2.96 | 0.99 | MVP | 1.441\|1.114\|1.405\|1.379 | 5.34 | .\|.\|.\|. | 4 | 0.99 | 0.99\|0.99\|1.00\|0.99 | 0.99 | 0.996 | 0.0039 | 2 |
| P18077 | 24.7 | 6.76 | 2.96 | 0.99 | RPL35A | 1.423\|1.398\|1.332\|1.212 | 5.37 | .\|.\|.\|. | 4 | 0.99 | 1.00\|1.00\|0.99\|0.99 | 0.99 | 0.9965 | 0.0024 | 36 |
| Q13573 | 22.9 | 2.44 | 1.78 | 1 | SNW1 | 1.190\|1.133\|1.454\|1.488 | 5.27 | .\|.\|.\|0.04 | 4 | 1 | 1.00\|0.99\|1.00\|1.00 | 1.00 | 0.998 | 0.0018 | 16 |
| Q16891 | 24.3 | 6.67 | 2.94 | 0.99 | IMMT | 1.516\|1.516\|1.140\|0.911 | 5.08 | .\|.\|.\|. | 4 | 0.99 | 1.00\|1.00\|0.99\|0.99 | 0.99 | 0.998 | 0.0029 | 15 |
| P13796 | 17 | 0.82 | 0.87 | 0.99 | LCP1 | 1.419\|1.242\|1.317\|1.331 | 5.31 | 0.05\|0.05\|0.23\|. | 4 | 0.99 | 0.98\|0.99\|0.99\|0.99 | 0.99 | 0.989 | 0.0057 | 15 |
| Q2TAY7 | 24.5 | 6.73 | 2.95 | 0.99 | SMU1 | 1.210\|1.123\|1.458\|1.448 | 5.24 | .\|.\|.\|. | 4 | 0.99 | 0.99\|0.99\|1.00\|1.00 | 0.99 | 0.996 | 0.0028 | 17 |
| Q2VIR3 | 22.7 | 6.31 | 2.87 | 0.99 | EIF2S3L | 0.941\|0.881\|0.899\|0.954 | 3.68 | .\|.\|.\|. | 4 | 0.99 | 0.99\|0.98\|0.99\|0.99 | 0.99 | 0.9895 | 0.0054 | 33 |

| | | | | | | | | | | | | | | | |
|---|---|---|---|---|---|---|---|---|---|---|---|---|---|---|---|
| Q15025 | 24.5 | 6.72 | 2.95 | 1 | TNIP1 | 1.370\|1.494\|1.203\|1.144 | 5.21 | .\|.\|.\|. | 4 | 1 | 1.00\|1.00\|1.00\|0.99 | 1.00 | 0.9995 | 0.0015 | 0 |
| P08195 | 24.5 | 6.72 | 2.95 | 0.99 | SLC3A2 | 1.418\|1.443\|1.202\|1.141 | 5.21 | .\|.\|.\|. | 4 | 0.99 | 0.99\|1.00\|0.99\|0.99 | 0.99 | 0.996 | 0.0038 | 15 |
| Q13247 | 24.6 | 6.74 | 2.95 | 1 | SRSF6 | 1.345\|1.228\|1.285\|1.396 | 5.26 | .\|.\|.\|. | 4 | 1 | 1.00\|1.00\|1.00\|1.00 | 1.00 | 0.9985 | 0.0017 | 44 |
| Q15459 | 24.5 | 6.73 | 2.95 | 0.99 | SF3A1 | 1.261\|1.110\|1.384\|1.444 | 5.20 | .\|.\|.\|. | 4 | 0.99 | 0.99\|0.99\|0.99\|1.00 | 0.99 | 0.996 | 0.0026 | 40 |
| P52701 | 24.6 | 6.73 | 2.95 | 1 | MSH6 | 1.336\|1.400\|1.242\|1.252 | 5.23 | .\|.\|.\|. | 4 | 1 | 1.00\|1.00\|1.00\|1.00 | 1.00 | 0.9985 | 0.0012 | 11 |
| P41250 | 24.5 | 6.73 | 2.95 | 0.99 | GARS | 1.354\|1.355\|1.259\|1.244 | 5.21 | .\|.\|.\|. | 4 | 0.99 | 0.99\|0.99\|0.99\|0.99 | 0.99 | 0.994 | 0.0031 | 15 |
| O43175 | 24.5 | 6.73 | 2.95 | 0.99 | PHGDH | 1.279\|1.383\|1.227\|1.319 | 5.21 | .\|.\|.\|. | 4 | 0.99 | 0.99\|0.99\|0.99\|0.99 | 0.99 | 0.9945 | 0.0038 | 51 |
| Q13283 | 24.5 | 6.72 | 2.95 | 0.99 | G3BP1 | 1.371\|1.362\|1.234\|1.198 | 5.17 | .\|.\|.\|. | 4 | 0.99 | 0.99\|1.00\|0.99\|0.99 | 0.99 | 0.995 | 0.0029 | 49 |
| Q7Z417 | 24.5 | 6.72 | 2.95 | 0.99 | NUFIP2 | 1.357\|1.361\|1.265\|1.177 | 5.16 | .\|.\|.\|. | 4 | 0.99 | 0.99\|0.99\|0.99\|0.99 | 0.99 | 0.994 | 0.0038 | 30 |
| Q702N8 | 24.5 | 6.71 | 2.95 | 0.99 | XIRP1 | 1.366\|1.180\|1.311\|1.281 | 5.14 | .\|.\|.\|. | 4 | 0.99 | 1.00\|0.99\|1.00\|0.99 | 1.00 | 0.997 | 0.0023 | 1 |
| Q14444 | 24.5 | 6.71 | 2.95 | 0.99 | CAPRIN1 | 1.326\|1.339\|1.220\|1.246 | 5.13 | .\|.\|.\|. | 4 | 0.99 | 0.99\|1.00\|0.99\|0.99 | 0.99 | 0.9965 | 0.0027 | 43 |
| P24666 | 24.4 | 6.70 | 2.95 | 0.99 | ACP1 | 1.231\|1.301\|1.251\|1.302 | 5.09 | .\|.\|.\|. | 4 | 1 | 1.00\|1.00\|1.00\|0.99 | 1.00 | 0.996 | 0.0023 | 8 |
| P62879 | 24.4 | 6.69 | 2.94 | 0.99 | GNB2 | 1.372\|1.258\|1.191\|1.235 | 5.06 | .\|.\|.\|. | 4 | 1 | 1.00\|1.00\|0.99\|0.99 | 1.00 | 0.997 | 0.0023 | 17 |
| Q12874 | 24.4 | 6.69 | 2.94 | 1 | SF3A3 | 1.189\|1.160\|1.348\|1.342 | 5.04 | .\|.\|.\|. | 4 | 1 | 0.99\|1.00\|1.00\|1.00 | 1.00 | 0.998 | 0.0019 | 32 |
| P21796 | 24.2 | 6.65 | 2.93 | 0.99 | VDAC1 | 1.409\|1.422\|1.061\|1.035 | 4.93 | .\|.\|.\|. | 4 | 0.99 | 1.00\|1.00\|0.99\|0.99 | 0.99 | 0.9975 | 0.003 | 13 |
| P06493 | 24.3 | 6.68 | 2.94 | 0.99 | CDK1 | 1.326\|1.343\|1.195\|1.168 | 5.03 | .\|.\|.\|. | 4 | 1 | 1.00\|1.00\|0.99\|0.99 | 0.99 | 0.9965 | 0.0024 | 27 |
| Q96FV9 | 24.3 | 6.69 | 2.94 | 0.99 | THOC1 | 1.291\|1.208\|1.233\|1.295 | 5.03 | .\|.\|.\|. | 4 | 0.99 | 1.00\|0.99\|0.99\|0.99 | 0.99 | 0.995 | 0.0034 | 11 |
| Q9BUJ2 | 22.4 | 2.16 | 1.66 | 0.99 | HNRNPUL1 | 1.280\|1.230\|1.270\|1.245 | 5.03 | .\|.\|.\|0.05 | 4 | 0.99 | 0.99\|0.99\|0.99\|0.99 | 0.99 | 0.993 | 0.0039 | 41 |
| Q13151 | 24.3 | 6.69 | 2.94 | 1 | HNRNPA0 | 1.239\|1.206\|1.265\|1.310 | 5.02 | .\|.\|.\|. | 4 | 1 | 0.99\|0.99\|1.00\|1.00 | 1.00 | 0.9975 | 0.0021 | 50 |
| Q9P258 | 24.3 | 6.68 | 2.94 | 0.99 | RCC2 | 1.257\|1.268\|1.241\|1.240 | 5.01 | .\|.\|.\|. | 4 | 0.99 | 1.00\|1.00\|0.99\|1.00 | 0.99 | 0.997 | 0.0024 | 35 |
| Q92878 | 23.7 | 6.52 | 2.91 | 0.99 | RAD50 | 1.460\|1.444\|0.864\|0.821 | 4.59 | .\|.\|.\|. | 4 | 1 | 1.00\|1.00\|0.99\|0.99 | 0.99 | 0.999 | 0.0036 | 14 |
| P27348 | 22.4 | 2.23 | 1.69 | 1 | YWHAQ | 1.274\|1.223\|1.245\|1.204 | 4.95 | 0.05\|.\|.\|. | 4 | 1 | 1.00\|0.99\|1.00\|1.00 | 1.00 | 0.9965 | 0.0021 | 52 |
| Q9UKM9 | 24.2 | 6.66 | 2.94 | 0.99 | RALY | 1.202\|1.120\|1.327\|1.273 | 4.92 | .\|.\|.\|. | 4 | 0.99 | 0.99\|0.99\|0.99\|0.99 | 0.99 | 0.993 | 0.0041 | 16 |
| P21580 | 24.3 | 6.67 | 2.94 | 0.99 | TNFAIP3 | 1.248\|1.235\|1.225\|1.228 | 4.94 | .\|.\|.\|. | 4 | 0.99 | 0.99\|1.00\|0.99\|0.99 | 0.99 | 0.9955 | 0.0034 | 1 |
| O75533 | 24.2 | 6.65 | 2.94 | 0.99 | SF3B1 | 1.150\|1.190\|1.315\|1.234 | 4.89 | .\|.\|.\|. | 4 | 1 | 0.99\|0.99\|0.99\|1.00 | 0.99 | 0.997 | 0.0026 | 46 |
| Q9BY77 | 24.2 | 6.65 | 2.93 | 1 | POLDIP3 | 1.279\|1.170\|1.187\|1.219 | 4.86 | .\|.\|.\|. | 4 | 1 | 1.00\|1.00\|1.00\|1.00 | 1.00 | 0.9975 | 0.0017 | 23 |
| P45880 | 24 | 6.61 | 2.93 | 1 | VDAC2 | 1.327\|1.315\|1.075\|1.037 | 4.75 | .\|.\|.\|. | 4 | 1 | 1.00\|1.00\|0.99\|0.99 | 1.00 | 0.9985 | 0.0021 | 14 |
| O00139 | 24.1 | 6.63 | 2.93 | 0.99 | KIF2A | 1.164\|1.181\|1.214\|1.247 | 4.81 | .\|.\|.\|. | 4 | 1 | 0.99\|0.99\|1.00\|0.99 | 0.99 | 0.995 | 0.0032 | 10 |
| O00148 | 23.9 | 6.59 | 2.92 | 0.99 | DDX39A | 1.187\|1.112\|1.200\|1.132 | 4.63 | .\|.\|.\|. | 4 | 0.99 | 0.99\|0.99\|0.99\|0.99 | 0.99 | 0.9925 | 0.0036 | 45 |
| P17987 | 24.1 | 6.63 | 2.93 | 0.99 | TCP1 | 1.209\|1.152\|1.228\|1.202 | 4.79 | .\|.\|.\|. | 4 | 0.99 | 0.99\|0.99\|0.99\|0.99 | 0.99 | 0.9945 | 0.004 | 56 |
| Q13769 | 24.1 | 6.62 | 2.93 | 0.99 | THOC5 | 1.208\|1.133\|1.262\|1.172 | 4.77 | .\|.\|.\|. | 4 | 0.99 | 0.99\|0.99\|1.00\|0.99 | 0.99 | 0.9955 | 0.0043 | 7 |
| Q8NCA5 | 22.3 | 6.21 | 2.85 | 0.98 | FAM98A | 0.867\|0.845\|0.798\|0.880 | 3.39 | .\|.\|.\|. | 4 | 0.98 | 0.98\|0.98\|0.98\|0.98 | 0.98 | 0.981 | 0.0071 | 24 |
| Q07955 | 24 | 6.61 | 2.93 | 0.99 | SRSF1 | 1.187\|1.089\|1.169\|1.290 | 4.74 | .\|.\|.\|. | 4 | 0.99 | 0.99\|0.99\|0.99\|0.99 | 0.99 | 0.9935 | 0.0046 | 45 |

| | | | | | | | | | | | | | | | |
|---|---|---|---|---|---|---|---|---|---|---|---|---|---|---|---|
| P07814 | 24 | 6.60 | 2.93 | 0.99 | EPRS | 1.268\|1.217\|1.157\|1.043 | 4.69 | .\|.\|.\|. | 4 | 0.99 | 0.99\|0.99\|0.99\|0.99 | 0.99 | 0.9945 | 0.0035 | 46 |
| Q14974 | 24 | 6.60 | 2.93 | 0.99 | KPNB1 | 1.242\|1.188\|1.175\|1.064 | 4.67 | .\|.\|.\|. | 4 | 0.99 | 0.99\|0.99\|0.99\|0.99 | 0.99 | 0.9905 | 0.0046 | 53 |
| P12956 | 24 | 6.60 | 2.93 | 0.99 | XRCC6 | 1.201\|1.186\|1.126\|1.167 | 4.68 | .\|.\|.\|. | 4 | 0.99 | 0.99\|0.99\|0.99\|1.00 | 0.99 | 0.995 | 0.0028 | 56 |
| Q99459 | 23.9 | 6.59 | 2.92 | 0.99 | CDC5L | 1.099\|1.035\|1.253\|1.249 | 4.64 | .\|.\|.\|. | 4 | 0.99 | 0.99\|0.99\|0.99\|1.00 | 0.99 | 0.9955 | 0.0034 | 23 |
| Q13557 | 23.7 | 6.52 | 2.91 | 0.99 | CAMK2D | 1.227\|1.382\|0.951\|0.913 | 4.47 | .\|.\|.\|. | 4 | 0.99 | 0.99\|1.00\|0.98\|0.98 | 0.99 | 0.9955 | 0.0047 | 5 |
| P09661 | 23.9 | 6.57 | 2.92 | 0.99 | SNRPA1 | 1.081\|1.035\|1.181\|1.290 | 4.59 | .\|.\|.\|. | 4 | 0.99 | 0.99\|0.99\|1.00\|0.99 | 0.99 | 0.9955 | 0.0033 | 39 |
| P04843 | 23.9 | 6.57 | 2.92 | 0.99 | RPN1 | 1.248\|1.202\|1.103\|1.040 | 4.59 | .\|.\|.\|. | 4 | 0.99 | 0.99\|1.00\|0.99\|0.99 | 0.99 | 0.9955 | 0.0029 | 29 |
| Q13951 | 23.6 | 6.50 | 2.91 | 0.99 | CBFB | 1.288\|1.285\|0.970\|0.860 | 4.40 | .\|.\|.\|. | 4 | 0.99 | 0.99\|0.99\|0.99\|0.99 | 0.99 | 0.994 | 0.0042 | 1 |
| Q13045 | 23.8 | 6.57 | 2.92 | 0.99 | FLII | 1.126\|1.124\|1.167\|1.142 | 4.56 | .\|.\|.\|. | 4 | 0.98 | 0.98\|0.99\|0.99\|0.99 | 0.99 | 0.989 | 0.0056 | 12 |
| Q9Y3X0 | 23.8 | 6.57 | 2.92 | 0.99 | CCDC9 | 1.135\|1.085\|1.122\|1.206 | 4.55 | .\|.\|.\|. | 4 | 0.99 | 0.99\|0.99\|0.99\|0.99 | 0.99 | 0.993 | 0.005 | 1 |
| P31942 | 23.8 | 6.56 | 2.92 | 0.99 | HNRNPH3 | 1.139\|1.066\|1.196\|1.142 | 4.54 | .\|.\|.\|. | 4 | 0.99 | 0.99\|0.99\|1.00\|0.99 | 0.99 | 0.995 | 0.0031 | 39 |
| Q9UJV9 | 23.8 | 6.57 | 2.92 | 0.99 | DDX41 | 1.133\|1.122\|1.132\|1.160 | 4.55 | .\|.\|.\|. | 4 | 0.99 | 1.00\|0.99\|0.99\|1.00 | 0.99 | 0.9965 | 0.0026 | 29 |
| P51116 | 23.7 | 6.54 | 2.91 | 0.99 | FXR2 | 1.203\|1.263\|1.022\|0.985 | 4.47 | .\|.\|.\|. | 4 | 0.99 | 1.00\|0.99\|0.98\|0.98 | 0.99 | 0.995 | 0.0048 | 15 |
| P49368 | 23.8 | 6.56 | 2.92 | 1 | CCT3 | 1.162\|1.131\|1.123\|1.099 | 4.52 | .\|.\|.\|. | 4 | 1 | 1.00\|0.99\|1.00\|1.00 | 1.00 | 0.9965 | 0.002 | 53 |
| P43490 | 23.6 | 6.51 | 2.91 | 0.99 | NAMPT | 1.336\|1.028\|1.136\|0.892 | 4.39 | .\|.\|.\|. | 4 | 1 | 1.00\|1.00\|0.99\|0.99 | 0.99 | 0.9975 | 0.0028 | 7 |
| P78371 | 21.8 | 2.10 | 1.63 | 0.99 | CCT2 | 1.134\|1.052\|1.190\|1.107 | 4.48 | .\|0.05\|.\|. | 4 | 0.99 | 0.99\|0.99\|0.99\|0.99 | 0.99 | 0.9905 | 0.0052 | 60 |
| P53814 | 23.7 | 6.54 | 2.91 | 0.98 | SMTN | 1.122\|1.200\|1.035\|1.109 | 4.47 | .\|.\|.\|. | 4 | 0.98 | 0.98\|0.99\|0.98\|0.98 | 0.98 | 0.989 | 0.0065 | 3 |
| Q14739 | 19.8 | 1.43 | 1.28 | 0.99 | LBR | 1.204\|1.245\|1.055\|0.881 | 4.39 | 0.05\|.\|0.05\|. | 4 | 0.99 | 0.99\|0.99\|0.99\|0.99 | 0.99 | 0.9945 | 0.0033 | 12 |
| Q9Y3Z3 | 23.7 | 6.54 | 2.91 | 0.99 | SAMHD1 | 1.147\|1.134\|1.099\|1.078 | 4.46 | .\|.\|.\|. | 4 | 0.99 | 0.99\|0.99\|0.99\|0.99 | 0.99 | 0.9925 | 0.0045 | 6 |
| P50990 | 23.7 | 6.54 | 2.91 | 0.99 | CCT8 | 1.132\|1.103\|1.142\|1.081 | 4.46 | .\|.\|.\|. | 4 | 0.99 | 0.99\|0.99\|0.99\|0.99 | 0.99 | 0.994 | 0.003 | 58 |
| Q6I9Y2 | 23.7 | 6.54 | 2.91 | 0.99 | THOC7 | 1.147\|1.052\|1.126\|1.115 | 4.44 | .\|.\|.\|. | 4 | 0.98 | 0.99\|0.98\|0.99\|0.99 | 0.99 | 0.9905 | 0.0053 | 4 |
| Q6IS14 | 19.7 | 1.52 | 1.33 | 0.97 | EIF5AL1 | 1.137\|1.140\|1.138\|1.134 | 4.55 | .\|.\|0.29\|. | 4 | 0.98 | 0.98\|0.97\|0.98\|0.97 | 0.97 | 0.9755 | 0.0084 | 36 |
| Q13242 | 23.7 | 6.54 | 2.91 | 0.99 | SRSF9 | 1.122\|1.110\|1.085\|1.123 | 4.44 | .\|.\|.\|. | 4 | 0.99 | 0.99\|1.00\|0.99\|0.99 | 0.99 | 0.996 | 0.0027 | 26 |
| Q9ULV4 | 23.7 | 6.53 | 2.91 | 0.99 | CORO1C | 1.188\|1.024\|1.146\|1.057 | 4.42 | .\|.\|.\|. | 4 | 0.99 | 0.99\|0.99\|1.00\|0.99 | 0.99 | 0.9955 | 0.0039 | 33 |
| Q12905 | 23.7 | 6.53 | 2.91 | 0.99 | ILF2 | 1.115\|1.033\|1.200\|1.065 | 4.41 | .\|.\|.\|. | 4 | 0.99 | 0.99\|0.99\|0.99\|0.99 | 0.99 | 0.9945 | 0.0038 | 49 |
| O95816 | 23.6 | 6.52 | 2.91 | 0.99 | BAG2 | 1.185\|1.158\|1.024\|1.038 | 4.41 | .\|.\|.\|. | 4 | 0.99 | 0.99\|1.00\|1.00\|0.99 | 0.99 | 0.995 | 0.0026 | 28 |
| P42704 | 23.6 | 6.52 | 2.91 | 0.99 | LRPPRC | 1.189\|1.059\|1.154\|0.995 | 4.40 | .\|.\|.\|. | 4 | 0.99 | 1.00\|0.99\|0.99\|0.99 | 0.99 | 0.996 | 0.0037 | 36 |
| P08865 | 21.8 | 2.57 | 1.84 | 0.99 | RPSA | 1.149\|1.096\|1.076\|1.100 | 4.42 | .\|.\|0.05\|. | 4 | 0.99 | 0.99\|0.99\|0.99\|0.99 | 0.99 | 0.9945 | 0.0036 | 57 |
| O60884 | 23.7 | 6.53 | 2.91 | 0.97 | DNAJA2 | 1.126\|1.122\|1.121\|1.036 | 4.41 | .\|.\|.\|. | 4 | 0.98 | 0.98\|0.97\|0.97\|0.98 | 0.97 | 0.9765 | 0.0082 | 34 |
| O43684 | 23.4 | 6.47 | 2.9 | 0.99 | BUB3 | 0.919\|0.866\|1.222\|1.250 | 4.26 | .\|.\|.\|. | 4 | 0.98 | 0.99\|0.98\|1.00\|0.99 | 0.99 | 0.995 | 0.0057 | 34 |
| O43390 | 23.6 | 6.51 | 2.91 | 0.99 | HNRNPR | 1.049\|1.030\|1.160\|1.101 | 4.34 | .\|.\|.\|. | 4 | 0.99 | 0.99\|0.99\|1.00\|0.99 | 0.99 | 0.9965 | 0.0036 | 58 |
| P50402 | 23.5 | 6.50 | 2.91 | 0.99 | EMD | 1.147\|1.159\|1.028\|0.982 | 4.32 | .\|.\|.\|. | 4 | 0.99 | 0.99\|1.00\|0.99\|0.99 | 0.99 | 0.9955 | 0.0031 | 24 |
| Q92945 | 23.5 | 6.50 | 2.91 | 0.99 | KHSRP | 1.041\|1.004\|1.090\|1.185 | 4.32 | .\|.\|.\|. | 4 | 0.99 | 0.99\|0.99\|0.99\|0.99 | 0.99 | 0.991 | 0.0044 | 44 |

| | | | | | | | | | | | | | | | |
|---|---|---|---|---|---|---|---|---|---|---|---|---|---|---|---|
| Q9Y230 | 23.5 | 6.49 | 2.91 | 0.99 | RUVBL2 | 1.134\|1.186\|1.025\|0.957 | 4.30 | .\|.\|.\|. | 4 | 0.99 | 0.99\|0.99\|0.99\|0.99 | 0.99 | 0.9925 | 0.004 | 47 |
| P51149 | 23.6 | 6.50 | 2.91 | 0.99 | RAB7A | 1.126\|1.144\|1.034\|1.014 | 4.32 | .\|.\|.\|. | 4 | 0.99 | 0.99\|0.99\|0.99\|0.98 | 0.99 | 0.9905 | 0.0054 | 12 |
| P62304 | 23.5 | 6.50 | 2.91 | 0.99 | SNRPE | 1.022\|1.022\|1.093\|1.173 | 4.31 | .\|.\|.\|. | 4 | 0.99 | 0.99\|0.99\|0.99\|0.99 | 0.99 | 0.99 | 0.005 | 34 |
| Q02978 | 23.5 | 6.48 | 2.9 | 0.99 | SLC25A11 | 1.131\|1.186\|0.958\|0.989 | 4.27 | .\|.\|.\|. | 4 | 0.99 | 0.99\|0.99\|0.99\|0.99 | 0.99 | 0.9945 | 0.0039 | 31 |
| P36873 | 23.5 | 6.49 | 2.91 | 0.99 | PPP1CC | 1.065\|1.051\|1.016\|1.156 | 4.29 | .\|.\|.\|. | 4 | 0.99 | 0.99\|0.99\|0.99\|0.99 | 0.99 | 0.9925 | 0.0043 | 44 |
| P05161 | 23.4 | 6.47 | 2.9 | 0.99 | ISG15 | 0.988\|1.195\|0.897\|1.141 | 4.22 | .\|.\|.\|. | 4 | 0.99 | 0.99\|0.99\|0.99\|0.99 | 0.99 | 0.9945 | 0.0035 | 1 |
| P36542 | 23.5 | 6.49 | 2.9 | 0.99 | ATP5C1 | 1.094\|1.066\|1.084\|1.022 | 4.27 | .\|.\|.\|. | 4 | 0.99 | 0.99\|0.99\|0.99\|0.99 | 0.99 | 0.9925 | 0.0045 | 26 |
| O75369 | 23.4 | 6.47 | 2.9 | 0.99 | FLNB | 1.090\|0.902\|1.143\|1.093 | 4.23 | .\|.\|.\|. | 4 | 0.99 | 0.99\|0.99\|0.99\|0.99 | 0.99 | 0.9935 | 0.0037 | 47 |
| P49458 | 12.3 | 0.54 | 0.62 | 0.99 | SRP9 | 0.951\|1.122\|1.081\|1.085 | 4.24 | 0.05\|0.06\|0.19\|0.18 | 4 | 0.99 | 0.99\|0.98\|0.99\|0.99 | 0.99 | 0.991 | 0.0058 | 30 |
| P17980 | 23.4 | 6.48 | 2.9 | 0.98 | PSMC3 | 1.091\|1.094\|1.075\|0.975 | 4.23 | .\|.\|.\|. | 4 | 0.98 | 0.98\|0.99\|0.97\|0.98 | 0.98 | 0.985 | 0.0072 | 32 |
| P55769 | 23.4 | 6.47 | 2.9 | 0.99 | SNU13 | 1.123\|0.954\|1.122\|1.019 | 4.22 | .\|.\|.\|. | 4 | 0.99 | 0.99\|0.99\|0.99\|0.99 | 0.99 | 0.994 | 0.0034 | 21 |
| Q15007 | 23.4 | 6.47 | 2.9 | 0.99 | WTAP | 1.123\|1.018\|0.997\|1.071 | 4.21 | .\|.\|.\|. | 4 | 1 | 0.99\|0.99\|0.99\|1.00 | 0.99 | 0.995 | 0.003 | 13 |
| Q9UBM7 | 23.3 | 6.43 | 2.89 | 0.99 | DHCR7 | 1.128\|1.182\|0.925\|0.885 | 4.12 | .\|.\|.\|. | 4 | 0.99 | 0.99\|0.99\|0.99\|0.98 | 0.99 | 0.9915 | 0.0052 | 4 |
| P16615 | 22.7 | 6.31 | 2.87 | 0.99 | ATP2A2 | 0.939\|0.973\|0.866\|0.915 | 3.69 | .\|.\|.\|. | 4 | 0.99 | 0.99\|0.99\|0.99\|0.99 | 0.99 | 0.9925 | 0.0041 | 26 |
| P40227 | 23.3 | 6.45 | 2.9 | 0.99 | CCT6A | 1.033\|0.907\|1.119\|1.088 | 4.15 | .\|.\|.\|. | 4 | 0.99 | 0.99\|0.99\|0.99\|0.99 | 0.99 | 0.991 | 0.0045 | 55 |
| P78527 | 23.3 | 6.45 | 2.9 | 0.99 | PRKDC | 1.103\|1.104\|0.992\|0.942 | 4.14 | .\|.\|.\|. | 4 | 0.99 | 0.99\|1.00\|0.99\|0.99 | 0.99 | 0.995 | 0.0039 | 50 |
| Q96J01 | 23.3 | 6.45 | 2.9 | 0.99 | THOC3 | 1.044\|1.026\|1.020\|1.058 | 4.15 | .\|.\|.\|. | 4 | 0.99 | 0.99\|0.99\|0.99\|0.99 | 0.99 | 0.9935 | 0.0041 | 15 |
| P02786 | 23.3 | 6.43 | 2.89 | 0.99 | TFRC | 1.141\|1.088\|0.982\|0.891 | 4.10 | .\|.\|.\|. | 4 | 0.99 | 0.99\|0.99\|0.99\|0.98 | 0.99 | 0.9935 | 0.0049 | 12 |
| P20700 | 23.1 | 6.39 | 2.89 | 0.98 | LMNB1 | 1.176\|1.101\|0.912\|0.817 | 4.01 | .\|.\|.\|. | 4 | 0.98 | 1.00\|0.99\|0.99\|0.97 | 0.99 | 0.995 | 0.0059 | 37 |
| Q13761 | 23.3 | 6.43256 | 2.89 | 0.99 | RUNX3 | 1.085\|1.060\|0.983\|0.957 | 4.085 | .\|.\|.\|. | 4 | 0.99 | 0.99\|0.99\|0.99\|0.98 | 0.99 | 0.992 | 0.0051 | 0 |
| P50991 | 23.3 | 6.44 | 2.89 | 0.99 | CCT4 | 1.036\|0.998\|1.007\|1.055 | 4.10 | .\|.\|.\|. | 4 | 1 | 0.99\|1.00\|0.99\|1.00 | 0.99 | 0.996 | 0.0025 | 56 |
| Q69YQ0 | 23.3 | 6.44 | 2.89 | 0.99 | SPECC1L | 0.997\|1.006\|1.007\|1.082 | 4.09 | .\|.\|.\|. | 4 | 0.99 | 0.99\|0.99\|0.99\|0.99 | 0.99 | 0.9925 | 0.0042 | 4 |
| Q15637 | 23.3 | 6.44 | 2.89 | 0.99 | SF1 | 1.001\|1.024\|0.984\|1.077 | 4.09 | .\|.\|.\|. | 4 | 0.99 | 0.99\|0.99\|0.99\|0.99 | 0.99 | 0.992 | 0.0041 | 26 |
| Q6UN15 | 23.3 | 6.44 | 2.89 | 0.99 | FIP1L1 | 1.037\|1.004\|1.033\|1.012 | 4.09 | .\|.\|.\|. | 4 | 0.99 | 0.99\|0.99\|0.99\|0.99 | 0.99 | 0.994 | 0.004 | 21 |
| Q8IYB3 | 23.2 | 6.42 | 2.89 | 1 | SRRM1 | 1.017\|1.004\|0.946\|1.086 | 4.05 | .\|.\|.\|. | 4 | 0.99 | 1.00\|0.99\|0.99\|1.00 | 1.00 | 0.997 | 0.0022 | 24 |
| Q9Y3F4 | 23.2 | 6.43 | 2.89 | 0.99 | STRAP | 1.048\|1.037\|0.998\|0.975 | 4.06 | .\|.\|.\|. | 4 | 0.99 | 1.00\|0.99\|0.99\|0.99 | 0.99 | 0.995 | 0.0034 | 37 |
| P62310 | 23.1 | 6.41 | 2.89 | 0.99 | LSM3 | 0.892\|0.907\|1.096\|1.111 | 4.01 | .\|.\|.\|. | 4 | 0.99 | 0.99\|0.99\|0.99\|0.99 | 0.99 | 0.992 | 0.0044 | 16 |
| Q96SB3 | 23.2 | 6.43 | 2.89 | 0.99 | PPP1R9B | 1.022\|0.995\|0.980\|1.056 | 4.05 | .\|.\|.\|. | 4 | 0.99 | 0.99\|0.99\|0.99\|0.99 | 0.99 | 0.994 | 0.0031 | 3 |
| P61026 | 23.2 | 6.42 | 2.89 | 0.99 | RAB10 | 1.074\|1.004\|0.995\|0.955 | 4.03 | .\|.\|.\|. | 4 | 0.99 | 0.99\|0.99\|0.99\|0.99 | 0.99 | 0.9925 | 0.0049 | 14 |
| Q92888 | 23.2 | 6.41 | 2.89 | 0.98 | ARHGEF1 | 1.047\|1.037\|0.980\|0.939 | 4.00 | .\|.\|.\|. | 4 | 0.98 | 0.99\|0.99\|0.98\|0.98 | 0.98 | 0.9875 | 0.0061 | 0 |
| P48444 | 23.1 | 6.40 | 2.89 | 0.99 | ARCN1 | 1.055\|0.985\|0.985\|0.950 | 3.98 | .\|.\|.\|. | 4 | 0.98 | 0.99\|0.99\|0.99\|0.99 | 0.99 | 0.9895 | 0.0053 | 24 |
| Q9UPN4 | 22.6 | 6.29 | 2.87 | 0.98 | CEP131 | 0.719\|0.696\|1.171\|1.132 | 3.72 | .\|.\|.\|. | 4 | 0.98 | 0.97\|0.97\|0.99\|0.99 | 0.98 | 0.9945 | 0.0072 | 2 |
| Q9BWJ5 | 15.8 | 0.81 | 0.86 | 0.99 | SF3B5 | 0.946\|0.876\|1.032\|1.087 | 3.94 | 0.05\|.\|0.20\|0.05 | 4 | 0.99 | 0.99\|0.99\|0.99\|0.99 | 0.99 | 0.9925 | 0.0047 | 20 |

| | | | | | | | | | | | | | | | |
|---|---|---|---|---|---|---|---|---|---|---|---|---|---|---|---|
| Q9H3U1 | 23.1 | 6.39 | 2.89 | 0.99 | UNC45A | 1.049\|1.002\|0.999\|0.904 | 3.96 | .\|.\|.\|. | 4 | 0.99 | 1.00\|0.99\|0.99\|0.99 | 0.99 | 0.996 | 0.0037 | 8 |
| P61158 | 23.1 | 6.39 | 2.88 | 0.98 | ACTR3 | 1.017\|0.920\|1.041\|0.948 | 3.93 | .\|.\|.\|. | 4 | 0.98 | 0.98\|0.97\|0.98\|0.98 | 0.98 | 0.9835 | 0.0075 | 17 |
| Q9Y613 | 23.1 | 6.39 | 2.88 | 0.99 | FHOD1 | 0.947\|0.927\|1.037\|1.016 | 3.93 | .\|.\|.\|. | 4 | 0.99 | 0.99\|0.98\|0.99\|0.99 | 0.99 | 0.9905 | 0.0055 | 1 |
| P53007 | 23 | 6.37 | 2.88 | 0.99 | SLC25A1 | 1.058\|1.008\|0.987\|0.850 | 3.90 | .\|.\|.\|. | 4 | 0.99 | 0.99\|0.99\|0.99\|0.98 | 0.99 | 0.9915 | 0.0055 | 9 |
| O75569 | 23 | 6.38 | 2.88 | 0.99 | PRKRA | 0.907\|0.902\|1.036\|1.059 | 3.90 | .\|.\|.\|. | 4 | 0.99 | 0.99\|0.99\|0.99\|0.99 | 0.99 | 0.9935 | 0.0042 | 12 |
| Q9NX63 | 22.7 | 6.30 | 2.87 | 0.99 | CHCHD3 | 1.149\|1.087\|0.791\|0.720 | 3.75 | .\|.\|.\|. | 4 | 0.99 | 0.99\|0.99\|0.98\|0.98 | 0.99 | 0.994 | 0.0057 | 11 |
| P14625 | 22.8 | 6.33 | 2.87 | 0.99 | HSP90B1 | 0.784\|0.777\|1.119\|1.101 | 3.78 | .\|.\|.\|. | 4 | 0.99 | 0.98\|0.98\|0.99\|1.00 | 0.99 | 0.995 | 0.0056 | 59 |
| P56134 | 23 | 6.37 | 2.88 | 0.98 | ATP5J2 | 1.008\|1.055\|0.920\|0.908 | 3.89 | .\|.\|.\|. | 4 | 0.98 | 0.98\|0.98\|0.98\|0.98 | 0.98 | 0.9845 | 0.0072 | 6 |
| Q53EZ4 | 23 | 6.38 | 2.88 | 0.99 | CEP55 | 0.883\|0.986\|0.997\|1.028 | 3.89 | .\|.\|.\|. | 4 | 1 | 0.99\|1.00\|0.99\|1.00 | 1.00 | 0.997 | 0.0023 | 1 |
| Q9NW64 | 23 | 6.37 | 2.88 | 0.99 | RBM22 | 0.883\|0.875\|1.031\|1.077 | 3.87 | .\|.\|.\|. | 4 | 0.99 | 0.99\|0.99\|0.99\|0.99 | 0.99 | 0.9915 | 0.005 | 8 |
| P22087 | 22.9 | 6.36 | 2.88 | 0.96 | FBL | 1.063\|0.838\|1.047\|0.913 | 3.86 | .\|.\|.\|. | 4 | 0.96 | 0.96\|0.96\|0.97\|0.96 | 0.96 | 0.9675 | 0.009 | 43 |
| Q16531 | 23 | 6.38 | 2.88 | 0.99 | DDB1 | 0.935\|0.957\|0.992\|1.015 | 3.90 | .\|.\|.\|. | 4 | 0.99 | 0.99\|0.99\|0.99\|0.99 | 0.99 | 0.9905 | 0.0052 | 42 |
| P61163 | 24.2 | 6.66 | 2.94 | 0.99 | ACTR1A | 1.229\|1.209\|1.261\|1.202 | 4.90 | .\|.\|.\|. | 4 | 0.99 | 0.99\|0.99\|0.99\|1.00 | 0.99 | 0.996 | 0.0032 | 34 |
| P61160 | 23 | 6.37 | 2.88 | 0.99 | ACTR2 | 1.015\|0.943\|0.992\|0.932 | 3.88 | .\|.\|.\|. | 4 | 0.99 | 0.99\|0.99\|1.00\|0.99 | 0.99 | 0.997 | 0.0043 | 19 |
| Q5SW79 | 23 | 6.36 | 2.88 | 0.99 | CEP170 | 1.010\|1.051\|0.917\|0.884 | 3.86 | .\|.\|.\|. | 4 | 0.99 | 0.99\|0.99\|0.99\|0.98 | 0.99 | 0.9915 | 0.0048 | 10 |
| Q9P2I0 | 23 | 6.36 | 2.88 | 0.99 | CPSF2 | 1.010\|0.965\|0.974\|0.906 | 3.86 | .\|.\|.\|. | 4 | 0.99 | 0.99\|1.00\|0.99\|0.99 | 0.99 | 0.995 | 0.0037 | 16 |
| P39656 | 22.7 | 6.30 | 2.87 | 0.98 | DDOST | 1.092\|1.062\|0.804\|0.761 | 3.72 | .\|.\|.\|. | 4 | 0.99 | 0.99\|0.99\|0.98\|0.97 | 0.98 | 0.9885 | 0.0067 | 18 |
| O94925 | 22.9 | 6.35 | 2.88 | 0.98 | GLS | 1.006\|0.953\|0.956\|0.902 | 3.82 | .\|.\|.\|. | 4 | 0.98 | 0.99\|0.98\|0.99\|0.98 | 0.98 | 0.9865 | 0.0064 | 3 |
| Q9Y265 | 22.9 | 6.34 | 2.88 | 0.99 | RUVBL1 | 1.016\|1.032\|0.891\|0.850 | 3.79 | .\|.\|.\|. | 4 | 0.99 | 1.00\|0.99\|0.99\|0.99 | 0.99 | 0.9955 | 0.0037 | 47 |
| Q3MHD2 | 22.9 | 6.35 | 2.88 | 0.96 | LSM12 | 0.995\|0.971\|0.934\|0.915 | 3.82 | .\|.\|.\|. | 4 | 0.96 | 0.96\|0.97\|0.96\|0.96 | 0.96 | 0.9655 | 0.009 | 27 |
| P62333 | 22.9 | 6.34 | 2.88 | 0.99 | PSMC6 | 0.963\|0.938\|0.943\|0.930 | 3.77 | .\|.\|.\|. | 4 | 0.99 | 0.99\|0.99\|0.99\|0.99 | 0.99 | 0.994 | 0.0047 | 30 |
| P62306 | 22.8 | 6.33 | 2.87 | 0.98 | SNRPF | 0.929\|0.923\|0.999\|0.914 | 3.77 | .\|.\|.\|. | 4 | 0.98 | 0.98\|0.98\|0.99\|0.98 | 0.98 | 0.9905 | 0.0064 | 36 |
| Q8NAV1 | 22.7 | 6.31 | 2.87 | 0.99 | PRPF38A | 0.828\|0.792\|1.064\|1.010 | 3.69 | .\|.\|.\|. | 4 | 0.99 | 0.99\|0.98\|0.99\|0.99 | 0.99 | 0.993 | 0.0051 | 15 |
| P22695 | 22.8 | 6.33 | 2.87 | 0.99 | UQCRC2 | 0.933\|0.941\|0.977\|0.904 | 3.75 | .\|.\|.\|. | 4 | 0.99 | 0.99\|0.99\|0.99\|0.99 | 0.99 | 0.988 | 0.0053 | 13 |
| P48643 | 22.8 | 6.32 | 2.87 | 0.99 | CCT5 | 0.951\|0.889\|1.005\|0.888 | 3.73 | .\|.\|.\|. | 4 | 0.99 | 0.99\|0.99\|0.99\|0.98 | 0.99 | 0.989 | 0.0056 | 56 |
| P30504 | 22.7 | 6.31 | 2.87 | 0.98 | HLA-C | 0.965\|1.025\|0.819\|0.899 | 3.71 | .\|.\|.\|. | 4 | 0.98 | 0.99\|0.98\|0.98\|0.99 | 0.98 | 0.988 | 0.0062 | 7 |
| Q9Y2D5 | 22.7 | 6.31 | 2.87 | 0.99 | AKAP2 | 0.875\|0.846\|0.917\|1.060 | 3.70 | .\|.\|.\|. | 4 | 0.98 | 0.98\|0.98\|0.99\|0.99 | 0.99 | 0.9885 | 0.0058 | 3 |
| Q13561 | 22.8 | 6.32 | 2.87 | 0.99 | DCTN2 | 0.907\|0.908\|0.964\|0.951 | 3.73 | .\|.\|.\|. | 4 | 0.99 | 0.99\|0.98\|0.99\|0.99 | 0.99 | 0.992 | 0.0054 | 30 |
| P35249 | 22.8 | 6.32 | 2.87 | 0.99 | RFC4 | 0.956\|0.974\|0.919\|0.872 | 3.72 | .\|.\|.\|. | 4 | 0.99 | 0.99\|0.99\|0.99\|0.99 | 0.99 | 0.992 | 0.0044 | 13 |
| Q7RTV0 | 19 | 1.38 | 1.25 | 0.98 | PHF5A | 0.864\|0.863\|0.989\|0.983 | 3.70 | .\|0.05\|0.05\|. | 4 | 0.99 | 0.98\|0.99\|0.99\|0.99 | 0.98 | 0.9865 | 0.006 | 19 |
| P53618 | 22.7 | 6.31 | 2.87 | 0.99 | COPB1 | 0.998\|0.921\|0.903\|0.881 | 3.70 | .\|.\|.\|. | 4 | 0.98 | 0.99\|0.99\|0.99\|0.99 | 0.99 | 0.991 | 0.005 | 21 |
| Q6PKG0 | 22.7 | 6.31 | 2.87 | 0.99 | LARP1 | 0.961\|0.936\|0.893\|0.905 | 3.70 | .\|.\|.\|. | 4 | 0.99 | 0.99\|1.00\|0.99\|0.99 | 0.99 | 0.995 | 0.0036 | 26 |
| Q06787 | 22.6 | 6.27 | 2.86 | 0.99 | FMR1 | 1.039\|1.013\|0.823\|0.732 | 3.61 | .\|.\|.\|. | 4 | 0.99 | 1.00\|1.00\|0.98\|0.98 | 0.99 | 0.9955 | 0.0056 | 14 |

| | | | | | | | | | | | | | | | |
|---|---|---|---|---|---|---|---|---|---|---|---|---|---|---|---|
| Q13123 | 22.7 | 6.31 | 2.87 | 0.99 | IK | 0.912\|0.847\|0.945\|0.977 | 3.68 | .\|.\|.\|. | 4 | 0.99 | 0.99\|0.99\|0.99\|0.99 | 0.99 | 0.994 | 0.0042 | 8 |
| P51571 | 22.7 | 6.30 | 2.87 | 0.99 | SSR4 | 0.953\|0.918\|0.971\|0.827 | 3.67 | .\|.\|.\|. | 4 | 0.99 | 0.99\|0.99\|0.99\|0.99 | 0.99 | 0.9885 | 0.0054 | 18 |
| Q9UN86 | 22.7 | 6.30 | 2.87 | 0.99 | G3BP2 | 0.962\|0.912\|0.906\|0.888 | 3.67 | .\|.\|.\|. | 4 | 0.99 | 0.99\|0.99\|0.98\|0.99 | 0.99 | 0.9915 | 0.0052 | 29 |
| P14868 | 22.7 | 6.30 | 2.87 | 0.99 | DARS | 0.975\|0.884\|0.911\|0.884 | 3.66 | .\|.\|.\|. | 4 | 0.99 | 0.99\|0.99\|0.99\|0.99 | 0.99 | 0.9895 | 0.0051 | 41 |
| Q9H223 | 22.7 | 6.30 | 2.87 | 0.98 | EHD4 | 0.888\|0.959\|0.872\|0.930 | 3.65 | .\|.\|.\|. | 4 | 0.98 | 0.98\|0.98\|0.98\|0.98 | 0.98 | 0.984 | 0.0066 | 1 |
| Q14257 | 22.6 | 6.29 | 2.87 | 0.98 | RCN2 | 0.883\|0.898\|0.885\|0.948 | 3.62 | .\|.\|.\|. | 4 | 0.98 | 0.98\|0.98\|0.99\|0.98 | 0.98 | 0.9865 | 0.0063 | 40 |
| Q99661 | 22.6 | 6.28 | 2.86 | 0.99 | KIF2C | 0.891\|0.915\|0.888\|0.896 | 3.59 | .\|.\|.\|. | 4 | 0.99 | 0.99\|0.99\|0.99\|0.99 | 0.99 | 0.992 | 0.0048 | 5 |
| Q9NZ01 | 22.5 | 6.26 | 2.86 | 0.99 | TECR | 0.959\|0.954\|0.826\|0.809 | 3.55 | .\|.\|.\|. | 4 | 0.99 | 0.99\|0.99\|0.99\|0.99 | 0.99 | 0.992 | 0.0046 | 12 |
| Q9P0L0 | 22.5 | 6.26 | 2.86 | 0.98 | VAPA | 0.921\|0.913\|0.907\|0.807 | 3.55 | .\|.\|.\|. | 4 | 0.98 | 0.98\|0.98\|0.98\|0.98 | 0.98 | 0.984 | 0.0069 | 7 |
| Q10570 | 22.5 | 6.26 | 2.86 | 0.98 | CPSF1 | 0.944\|0.858\|0.896\|0.844 | 3.54 | .\|.\|.\|. | 4 | 0.98 | 0.99\|0.98\|0.98\|0.98 | 0.98 | 0.985 | 0.0071 | 21 |
| Q13148 | 22.5 | 6.26 | 2.86 | 0.99 | TARDBP | 0.931\|0.867\|0.891\|0.847 | 3.54 | .\|.\|.\|. | 4 | 0.99 | 0.99\|0.99\|0.99\|0.99 | 0.99 | 0.992 | 0.0043 | 30 |
| P20592 | 22.1 | 6.17 | 2.84 | 0.97 | MX2 | 0.688\|1.040\|0.641\|0.986 | 3.36 | .\|.\|.\|. | 4 | 0.97 | 0.96\|0.98\|0.95\|0.98 | 0.97 | 0.984 | 0.0086 | 0 |
| O14579 | 22.4 | 6.25 | 2.86 | 0.99 | COPE | 0.813\|0.812\|0.977\|0.899 | 3.50 | .\|.\|.\|. | 4 | 0.99 | 0.99\|0.98\|0.99\|0.99 | 0.99 | 0.9915 | 0.0056 | 11 |
| Q15428 | 22.4 | 6.24 | 2.86 | 0.98 | SF3A2 | 0.807\|0.811\|0.949\|0.926 | 3.49 | .\|.\|.\|. | 4 | 0.98 | 0.97\|0.98\|0.99\|0.98 | 0.98 | 0.986 | 0.0074 | 21 |
| Q01196 | 22.4 | 6.23 | 2.85 | 0.98 | RUNX1 | 0.964\|0.883\|0.871\|0.756 | 3.48 | .\|.\|.\|. | 4 | 0.98 | 0.98\|0.98\|0.98\|0.96 | 0.98 | 0.9845 | 0.0078 | 1 |
| Q99623 | 22.4 | 6.23 | 2.85 | 0.98 | PHB2 | 0.948\|0.897\|0.798\|0.818 | 3.46 | .\|.\|.\|. | 4 | 0.98 | 0.98\|0.98\|0.97\|0.98 | 0.98 | 0.982 | 0.0079 | 33 |
| Q7Z6E9 | 22.4 | 6.23 | 2.85 | 0.98 | RBBP6 | 0.858\|0.796\|0.906\|0.895 | 3.46 | .\|.\|.\|. | 4 | 0.98 | 0.98\|0.98\|0.99\|0.98 | 0.98 | 0.9875 | 0.0066 | 6 |
| Q02040 | 22.4 | 6.23 | 2.85 | 0.99 | AKAP17A | 0.835\|0.849\|0.891\|0.865 | 3.44 | .\|.\|.\|. | 4 | 0.99 | 1.00\|0.99\|0.99\|0.99 | 0.99 | 0.9955 | 0.0032 | 4 |
| P02794 | 20.5 | 2.51 | 1.81 | 0.98 | FTH1 | 0.868\|0.854\|0.811\|0.902 | 3.44 | .\|.\|0.05\|. | 4 | 0.98 | 0.99\|0.98\|0.98\|0.99 | 0.98 | 0.9875 | 0.0065 | 1 |
| Q96DI7 | 16.6 | 0.94 | 0.95 | 0.99 | SNRNP40 | 0.785\|0.736\|0.930\|0.939 | 3.39 | 0.05\|.\|0.05\|0.05 | 4 | 0.99 | 0.99\|0.99\|1.00\|1.00 | 0.99 | 0.9975 | 0.0027 | 24 |
| O15143 | 20.4 | 2.47 | 1.79 | 0.98 | ARPC1B | 0.915\|0.788\|0.847\|0.874 | 3.42 | .\|.\|0.05\|. | 4 | 0.98 | 0.99\|0.98\|0.98\|0.99 | 0.98 | 0.9895 | 0.0063 | 7 |
| P29692 | 22.3 | 6.21 | 2.85 | 0.98 | EEF1D | 0.824\|0.764\|0.901\|0.920 | 3.41 | .\|.\|.\|. | 4 | 0.98 | 0.98\|0.97\|0.98\|0.98 | 0.98 | 0.9795 | 0.008 | 53 |
| Q9UPN7 | 22.3 | 6.21 | 2.85 | 0.98 | PPP6R1 | 0.904\|0.920\|0.801\|0.778 | 3.40 | .\|.\|.\|. | 4 | 0.98 | 0.98\|0.98\|0.97\|0.97 | 0.98 | 0.9825 | 0.0079 | 3 |
| P09382 | 22.3 | 6.21 | 2.85 | 0.98 | LGALS1 | 0.855\|0.841\|0.830\|0.869 | 3.40 | .\|.\|.\|. | 4 | 0.98 | 0.98\|0.98\|0.98\|0.98 | 0.98 | 0.984 | 0.0074 | 8 |
| Q96FS4 | 22.3 | 6.20 | 2.85 | 0.98 | SIPA1 | 0.832\|0.879\|0.807\|0.861 | 3.38 | .\|.\|.\|. | 4 | 0.98 | 0.98\|0.98\|0.98\|0.98 | 0.98 | 0.9835 | 0.0068 | 1 |
| Q9Y608 | 22.3 | 6.20 | 2.85 | 0.98 | LRRFIP2 | 0.866\|0.850\|0.833\|0.832 | 3.38 | .\|.\|.\|. | 4 | 0.98 | 0.99\|0.98\|0.98\|0.98 | 0.98 | 0.986 | 0.0061 | 7 |
| O15042 | 22.2 | 6.19 | 2.85 | 0.98 | U2SURP | 0.820\|0.807\|0.833\|0.884 | 3.34 | .\|.\|.\|. | 4 | 0.98 | 0.98\|0.98\|0.98\|0.99 | 0.98 | 0.985 | 0.0068 | 27 |
| Q96C19 | 22.2 | 6.18 | 2.84 | 0.98 | EFHD2 | 0.768\|0.806\|0.805\|0.941 | 3.32 | .\|.\|.\|. | 4 | 0.98 | 0.98\|0.98\|0.98\|0.99 | 0.98 | 0.99 | 0.0061 | 3 |
| P28066 | 22.2 | 6.19 | 2.85 | 0.97 | PSMA5 | 0.865\|0.870\|0.803\|0.796 | 3.33 | .\|.\|.\|. | 4 | 0.98 | 0.98\|0.98\|0.98\|0.97 | 0.97 | 0.979 | 0.0082 | 38 |
| Q13077 | 22.2 | 6.18 | 2.84 | 1 | TRAF1 | 0.810\|0.847\|0.846\|0.820 | 3.32 | .\|.\|.\|. | 4 | 0.99 | 0.99\|1.00\|1.00\|1.00 | 1.00 | 0.9965 | 0.0021 | 1 |
| P08574 | 22.1 | 6.17 | 2.84 | 0.98 | CYC1 | 0.875\|0.863\|0.794\|0.772 | 3.30 | .\|.\|.\|. | 4 | 0.98 | 0.99\|0.98\|0.98\|0.97 | 0.98 | 0.9855 | 0.0074 | 3 |
| Q9NTJ3 | 22.1 | 6.17 | 2.84 | 0.98 | SMC4 | 0.879\|0.846\|0.810\|0.754 | 3.29 | .\|.\|.\|. | 4 | 0.98 | 0.99\|0.99\|0.98\|0.98 | 0.98 | 0.9885 | 0.006 | 27 |
| O00165 | 22.1 | 6.16 | 2.84 | 0.98 | HAX1 | 0.931\|0.789\|0.817\|0.729 | 3.27 | .\|.\|.\|. | 4 | 0.98 | 0.99\|0.98\|0.99\|0.97 | 0.98 | 0.989 | 0.0067 | 8 |

| | | | | | | | | | | | | | | | |
|---|---|---|---|---|---|---|---|---|---|---|---|---|---|---|---|
| Q9NZR1 | 22.1 | 6.17 | 2.84 | 0.99 | TMOD2 | 0.851\|0.797\|0.828\|0.816 | 3.29 | .\|.\|.\|. | 4 | 0.99 | 0.99\|0.99\|0.99\|0.99 | 0.99 | 0.99 | 0.0049 | 7 |
| O00743 | 22.1 | 6.16 | 2.84 | 0.97 | PPP6C | 0.886\|0.833\|0.779\|0.780 | 3.28 | .\|.\|.\|. | 4 | 0.98 | 0.98\|0.97\|0.97\|0.97 | 0.97 | 0.9775 | 0.0085 | 5 |
| P52732 | 21.8 | 6.09 | 2.83 | 0.98 | KIF11 | 0.799\|1.006\|0.650\|0.680 | 3.14 | .\|.\|.\|. | 4 | 0.98 | 0.99\|0.99\|0.97\|0.97 | 0.98 | 0.993 | 0.0069 | 41 |
| P24534 | 22.1 | 6.15 | 2.84 | 0.99 | EEF1B2 | 0.793\|0.792\|0.850\|0.805 | 3.24 | .\|.\|.\|. | 4 | 0.99 | 0.98\|0.98\|0.99\|0.98 | 0.99 | 0.99 | 0.0058 | 54 |
| Q9H7N4 | 22 | 6.14 | 2.84 | 0.99 | SCAF1 | 0.834\|0.752\|0.818\|0.813 | 3.22 | .\|.\|.\|. | 4 | 0.99 | 0.99\|0.98\|0.99\|0.99 | 0.99 | 0.993 | 0.0045 | 3 |
| Q13595 | 22 | 6.14 | 2.84 | 1 | TRA2A | 0.813\|0.796\|0.825\|0.779 | 3.21 | .\|.\|.\|. | 4 | 1 | 1.00\|0.99\|1.00\|1.00 | 1.00 | 0.9975 | 0.0019 | 15 |
| P78406 | 22 | 6.14 | 2.84 | 0.98 | RAE1 | 0.790\|0.773\|0.840\|0.795 | 3.20 | .\|.\|.\|. | 4 | 0.98 | 0.98\|0.98\|0.99\|0.98 | 0.98 | 0.9885 | 0.0065 | 16 |
| Q00653 | 21.9 | 6.12 | 2.83 | 0.98 | NFKB2 | 0.849\|0.772\|0.830\|0.712 | 3.16 | .\|.\|.\|. | 4 | 0.99 | 0.99\|0.98\|0.99\|0.98 | 0.98 | 0.992 | 0.006 | 0 |
| Q1KMD3 | 20.1 | 2.08 | 1.63 | 0.98 | HNRNPUL2 | 0.831\|0.755\|0.781\|0.806 | 3.17 | 0.05\|.\|.\|. | 4 | 0.98 | 0.99\|0.98\|0.99\|0.98 | 0.98 | 0.987 | 0.0062 | 20 |
| P41240 | 21.9 | 6.12 | 2.83 | 0.98 | CSK | 0.808\|0.785\|0.749\|0.809 | 3.15 | .\|.\|.\|. | 4 | 0.98 | 0.98\|0.98\|0.97\|0.98 | 0.98 | 0.9835 | 0.0075 | 2 |
| P48047 | 18.3 | 1.59 | 1.37 | 0.98 | ATP5O | 0.834\|0.830\|0.723\|0.744 | 3.13 | .\|.\|0.18\|. | 4 | 0.98 | 0.98\|0.99\|0.97\|0.97 | 0.98 | 0.9865 | 0.0078 | 25 |
| P61221 | 21.9 | 6.11 | 2.83 | 0.96 | ABCE1 | 0.826\|0.785\|0.780\|0.743 | 3.13 | .\|.\|.\|. | 4 | 0.97 | 0.97\|0.96\|0.96\|0.96 | 0.96 | 0.9665 | 0.0092 | 10 |
| Q9Y5M8 | 21.8 | 6.10 | 2.83 | 0.98 | SRPRB | 0.827\|0.849\|0.718\|0.719 | 3.11 | .\|.\|.\|. | 4 | 0.98 | 0.99\|0.99\|0.98\|0.98 | 0.98 | 0.987 | 0.0064 | 6 |
| Q99832 | 21.8 | 6.10 | 2.83 | 0.98 | CCT7 | 0.767\|0.771\|0.836\|0.736 | 3.11 | .\|.\|.\|. | 4 | 0.98 | 0.97\|0.97\|0.99\|0.97 | 0.98 | 0.988 | 0.0079 | 51 |
| P31939 | 21.8 | 6.09 | 2.83 | 0.97 | ATIC | 0.803\|0.777\|0.772\|0.739 | 3.09 | .\|.\|.\|. | 4 | 0.97 | 0.97\|0.98\|0.98\|0.97 | 0.97 | 0.978 | 0.0083 | 15 |
| Q53GQ0 | 21.7 | 6.08 | 2.82 | 0.98 | HSD17B12 | 0.831\|0.809\|0.743\|0.678 | 3.06 | .\|.\|.\|. | 4 | 0.98 | 0.98\|0.98\|0.98\|0.97 | 0.98 | 0.9845 | 0.0069 | 5 |
| O00214 | 21.7 | 6.07 | 2.82 | 0.98 | LGALS8 | 0.689\|0.764\|0.722\|0.854 | 3.03 | .\|.\|.\|. | 4 | 0.98 | 0.97\|0.98\|0.98\|0.99 | 0.98 | 0.9905 | 0.007 | 1 |
| P40926 | 21.7 | 6.06 | 2.82 | 0.99 | MDH2 | 0.716\|0.843\|0.756\|0.707 | 3.02 | .\|.\|.\|. | 4 | 0.99 | 0.99\|0.99\|0.99\|0.98 | 0.99 | 0.994 | 0.0045 | 42 |
| P08579 | 21.7 | 6.07 | 2.82 | 0.98 | SNRPB2 | 0.773\|0.736\|0.785\|0.740 | 3.04 | .\|.\|.\|. | 4 | 0.99 | 0.98\|0.98\|0.98\|0.98 | 0.98 | 0.9845 | 0.0068 | 31 |
| Q02818 | 21.7 | 6.06 | 2.82 | 0.98 | NUCB1 | 0.787\|0.805\|0.706\|0.709 | 3.01 | .\|.\|.\|. | 4 | 0.98 | 0.99\|0.99\|0.98\|0.98 | 0.99 | 0.99 | 0.0059 | 3 |
| Q13523 | 21.6 | 6.06 | 2.82 | 0.99 | PRPF4B | 0.756\|0.739\|0.759\|0.743 | 3.00 | .\|.\|.\|. | 4 | 0.99 | 0.99\|0.99\|0.99\|0.99 | 0.99 | 0.989 | 0.0051 | 8 |
| O95864 | 21.6 | 6.04 | 2.82 | 0.98 | FADS2 | 0.722\|0.821\|0.694\|0.734 | 2.97 | .\|.\|.\|. | 4 | 0.98 | 0.98\|0.99\|0.97\|0.98 | 0.98 | 0.9855 | 0.007 | 1 |
| O60306 | 21.6 | 6.04 | 2.82 | 0.98 | AQR | 0.706\|0.675\|0.749\|0.831 | 2.96 | .\|.\|.\|. | 4 | 0.98 | 0.98\|0.97\|0.99\|0.98 | 0.98 | 0.9865 | 0.0073 | 8 |
| P11387 | 21.6 | 6.04 | 2.82 | 0.98 | TOP1 | 0.759\|0.777\|0.709\|0.721 | 2.97 | .\|.\|.\|. | 4 | 0.98 | 0.98\|0.99\|0.98\|0.98 | 0.98 | 0.987 | 0.0062 | 40 |
| Q96T37 | 21.6 | 6.04 | 2.82 | 0.97 | RBM15 | 0.759\|0.741\|0.674\|0.785 | 2.96 | .\|.\|.\|. | 4 | 0.97 | 0.97\|0.98\|0.97\|0.98 | 0.97 | 0.9815 | 0.0082 | 7 |
| P08754 | 21.6 | 6.04 | 2.82 | 0.97 | GNAI3 | 0.784\|0.746\|0.730\|0.705 | 2.96 | .\|.\|.\|. | 4 | 0.97 | 0.98\|0.97\|0.97\|0.96 | 0.97 | 0.975 | 0.0085 | 16 |
| Q14699 | 21.6 | 6.04 | 2.82 | 0.97 | RFTN1 | 0.747\|0.699\|0.789\|0.723 | 2.96 | .\|.\|.\|. | 4 | 0.97 | 0.97\|0.97\|0.97\|0.98 | 0.97 | 0.9785 | 0.0084 | 2 |
| Q8TBC3 | 21.6 | 6.04 | 2.82 | 0.98 | SHKBP1 | 0.757\|0.714\|0.719\|0.772 | 2.96 | .\|.\|.\|. | 4 | 0.98 | 0.97\|0.98\|0.98\|0.98 | 0.98 | 0.98 | 0.008 | 2 |
| P24539 | 21.5 | 6.01 | 2.81 | 0.95 | ATP5F1 | 0.788\|0.824\|0.673\|0.624 | 2.91 | .\|.\|.\|. | 4 | 0.95 | 0.96\|0.96\|0.95\|0.94 | 0.95 | 0.9635 | 0.0094 | 15 |
| Q9Y305 | 21.2 | 5.95 | 2.8 | 0.96 | ACOT9 | 0.578\|0.564\|0.899\|0.740 | 2.78 | .\|.\|.\|. | 4 | 0.97 | 0.96\|0.94\|0.98\|0.98 | 0.96 | 0.983 | 0.0087 | 2 |
| Q8IWX8 | 21.4 | 6.00 | 2.81 | 0.98 | CHERP | 0.690\|0.690\|0.719\|0.771 | 2.87 | .\|.\|.\|. | 4 | 0.97 | 0.98\|0.98\|0.98\|0.98 | 0.98 | 0.9815 | 0.0076 | 23 |
| P46087 | 21.4 | 6.00 | 2.81 | 0.98 | NOP2 | 0.752\|0.715\|0.723\|0.676 | 2.87 | .\|.\|.\|. | 4 | 0.98 | 0.99\|0.98\|0.98\|0.97 | 0.98 | 0.986 | 0.0074 | 29 |
| P34897 | 21.3 | 5.99 | 2.8 | 0.98 | SHMT2 | 0.784\|0.698\|0.711\|0.642 | 2.84 | .\|.\|.\|. | 4 | 0.97 | 0.98\|0.98\|0.98\|0.97 | 0.98 | 0.9845 | 0.0078 | 39 |

| | | | | | | | | | | | | | | | |
|---|---|---|---|---|---|---|---|---|---|---|---|---|---|---|---|
| Q15427 | 21.4 | 5.99 | 2.81 | 0.98 | SF3B4 | 0.700\|0.635\|0.747\|0.756 | 2.84 | .\|.\|.\|. | 4 | 0.98 | 0.98\|0.98\|0.98\|0.98 | 0.98 | 0.984 | 0.0066 | 33 |
| Q15125 | 21.2 | 5.94 | 2.8 | 0.96 | EBP | 0.760\|0.741\|0.639\|0.604 | 2.74 | .\|.\|.\|. | 4 | 0.96 | 0.97\|0.97\|0.95\|0.96 | 0.96 | 0.9725 | 0.0089 | 3 |
| Q63HN8 | 21.2 | 5.94 | 2.8 | 0.98 | RNF213 | 0.724\|0.759\|0.623\|0.638 | 2.74 | .\|.\|.\|. | 4 | 0.98 | 0.98\|0.98\|0.98\|0.96 | 0.98 | 0.983 | 0.0081 | 0 |
| P63162 | 24.8 | 6.79 | 2.96 | 1 | SNRPN | 1.328\|1.235\|1.432\|1.478 | 5.47 | .\|.\|.\|. | 4 | 1 | 1.00\|1.00\|1.00\|1.00 | 1.00 | 0.999 | 0.0011 | 53 |
| P38159 | 26.2 | 7.13 | 3.02 | 1 | RBMX | 1.821\|1.754\|1.880\|1.773 | 7.23 | .\|.\|.\|. | 4 | 1 | 1.00\|1.00\|1.00\|1.00 | 1.00 | 0.9975 | 0.0017 | 60 |
| Q8N684 | 20.8 | 5.85 | 2.78 | 0.98 | CPSF7 | 0.637\|0.629\|0.619\|0.647 | 2.53 | .\|.\|.\|. | 4 | 0.97 | 0.98\|0.98\|0.97\|0.98 | 0.98 | 0.983 | 0.0072 | 18 |
| P00338 | 12.6 | 0.82 | 0.87 | 0.96 | LDHA | 0.190\|0.188\|0.798\|0.680 | 1.86 | 0.05\|0.05\|0.05\|. | 4 | 0.96 | 0.93\|0.94\|1.00\|0.99 | 0.96 | 0.995 | 0.0088 | 47 |
| Q13155 | 20.6 | 5.81 | 2.77 | 0.96 | AIMP2 | 0.652\|0.614\|0.611\|0.578 | 2.46 | .\|.\|.\|. | 4 | 0.97 | 0.97\|0.96\|0.95\|0.96 | 0.96 | 0.9695 | 0.0091 | 21 |
| **P17844** | 24.4 | 2.7598 | 1.91 | 1 | DDX5 | 1.947\|1.801\|1.831\|1.871 | 7.451 | .\|.\|0.05\|. | 4 | 1 | 1.00\|1.00\|1.00\|1.00 | 0.9985 | 0.9995 | 9.00E-04 | **76** |
| **P63151** | 19.6 | 5.5651 | 2.71 | 0.13 | PPP2R2A | 0.721\|0.712\|0.727\|0.178 | 2.338 | .\|.\|.\|. | 4 | 0.14 | 0.17\|0.17\|0.17\|0.02 | **0.1306** | 0.1695 | 0.0885 | 25 |
| **P30153** | 19.8 | 5.6017 | 2.72 | 0.79 | PPP2R1A | 0.750\|0.834\|0.696\|0.173 | 2.452 | .\|.\|.\|. | 4 | 0.78 | 0.97\|0.97\|0.96\|0.24 | **0.7861** | 0.974 | 0.0174 | 46 |
| **P67775** | 19.8 | 5.6241 | 2.73 | 0.76 | PPP2CA | 0.740\|0.198\|0.729\|0.703 | 2.37 | .\|.\|.\|. | 4 | 0.78 | 0.93\|0.26\|0.93\|0.93 | **0.7615** | 0.9305 | 0.0229 | 33 |

| | |
|---|---|
| P17844 | Present in CRAPome >60 % |
| P63151 | Poor SP due to low intensity in one replicate |
| P30153 | Poor SP due to low intensity in one replicate |
| P67775 | Poor SP due to low intensity in one replicate |
