## Supplementary Table 1B for "Protein Interactome of the Cancerous Inhibitor of Protein Phosphatase 2A (CIP2A) in Th17 Cells"

### **Supplementry Table 1B:** IPA Assigned GO Annotation for the CIP2A Interactome

Table lists cellular location and functional class for the proteins detected in the CIP2A interactome. Ingenuity Pathway Analysis (IPA) was used to provide this annotation

| ID | Symbol | Entrez Gene Name | Location | Type(s) |
|---|---|---|---|---|
| B0I1T2 | MYO1G | myosin IG | Cytoplasm | other |
| Q8TCG1 | CIP2A | cell proliferation regulating inhibitor of protei | Cytoplasm | other |
| P11940 | PABPC1 | poly(A) binding protein cytoplasmic 1 | Cytoplasm | translation regulator |
| Q9NVI7 | ATAD3A | ATPase family, AAA domain containing 3A | Cytoplasm | other |
| Q9NVJ2 | ARL8B | ADP ribosylation factor like GTPase 8B | Plasma Membrane | enzyme |
| P19474 | TRIM21 | tripartite motif containing 21 | Nucleus | enzyme |
| P35580 | MYH10 | myosin heavy chain 10 | Cytoplasm | enzyme |
| Q01082 | SPTBN1 | spectrin beta, non-erythrocytic 1 | Plasma Membrane | other |
| P07910 | HNRNPC | heterogeneous nuclear ribonucleoprotein C (( | Nucleus | other |
| Q8NF50 | DOCK8 | dedicator of cytokinesis 8 | Cytoplasm | other |
| Q15149 | PLEC | plectin | Cytoplasm | other |
| Q9UKV3 | ACIN1 | apoptotic chromatin condensation inducer 1 | Nucleus | enzyme |
| Q9NYL9 | TMOD3 | tropomodulin 3 | Cytoplasm | other |
| P0DN76 | U2AF1/U2AF1 | U2 small nuclear RNA auxiliary factor 1 | Nucleus | other |
| Q6NYC8 | PPP1R18 | protein phosphatase 1 regulatory subunit 18 | Other | other |
| Q9H307 | PNN | pinin, desmosome associated protein | Plasma Membrane | other |
| Q7Z2W4 | ZC3HAV1 | zinc finger CCCH-type containing, antiviral 1 | Plasma Membrane | other |
| Q13428 | TCOF1 | treacle ribosome biogenesis factor 1 | Nucleus | transporter |
| P52907 | CAPZA1 | capping actin protein of muscle Z-line alpha s | Cytoplasm | other |
| Q7Z406 | MYH14 | myosin heavy chain 14 | Extracellular Space | enzyme |
| O43795 | MYO1B | myosin IB | Cytoplasm | enzyme |
| P38919 | EIF4A3 | eukaryotic translation initiation factor 4A3 | Nucleus | enzyme |
| O00159 | MYO1C | myosin IC | Cytoplasm | enzyme |
| Q9Y2W1 | THRAP3 | thyroid hormone receptor associated protein | Nucleus | transcription regulator |
| Q92614 | MYO18A | myosin XVIIIA | Cytoplasm | other |
| P62136 | PPP1CA | protein phosphatase 1 catalytic subunit alpha | Cytoplasm | phosphatase |
| P14866 | HNRNPL | heterogeneous nuclear ribonucleoprotein L | Nucleus | other |
| P47756 | CAPZB | capping actin protein of muscle Z-line beta su | Cytoplasm | other |
| Q9NYF8 | BCLAF1 | BCL2 associated transcription factor 1 | Nucleus | transcription regulator |
| O14974 | PPP1R12A | protein phosphatase 1 regulatory subunit 12A | Cytoplasm | phosphatase |
| Q6P2E9 | EDC4 | enhancer of mRNA decapping 4 | Cytoplasm | other |
| P12268 | IMPDH2 | inosine monophosphate dehydrogenase 2 | Cytoplasm | enzyme |
| Q5T9A4 | ATAD3B | ATPase family, AAA domain containing 3B | Nucleus | other |
| Q96HS1 | PGAM5 | PGAM family member 5, mitochondrial serine | Cytoplasm | enzyme |
| P62906 | RPL10A | ribosomal protein L10a | Nucleus | other |
| O00422 | SAP18 | Sin3A associated protein 18 | Nucleus | transcription regulator |
| Q02878 | RPL6 | ribosomal protein L6 | Nucleus | other |
| Q9Y6N5 | SQOR | sulfide quinone oxidoreductase | Cytoplasm | enzyme |
| P43243 | MATR3 | matrin 3 | Nucleus | other |
| P04114 | APOB | apolipoprotein B | Extracellular Space | transporter |
| P84103 | SRSF3 | serine and arginine rich splicing factor 3 | Nucleus | other |
| Q92804 | TAF15 | TATA-box binding protein associated factor 1! | Nucleus | other |
| P50914 | RPL14 | ribosomal protein L14 | Cytoplasm | other |
| P62834 | RAP1A | RAP1A, member of RAS oncogene family | Cytoplasm | enzyme |
| Q15154 | PCM1 | pericentriolar material 1 | Cytoplasm | other |

| | | | | |
|---|---|---|---|---|
| P27694 | RPA1 | replication protein A1 | Nucleus | other |
| Q16666 | IFI16 | interferon gamma inducible protein 16 | Nucleus | transcription regulator |
| P06753 | TPM3 | tropomyosin 3 | Cytoplasm | other |
| O75643 | SNRNP200 | small nuclear ribonucleoprotein U5 subunit 2( | Nucleus | enzyme |
| P26599 | PTBP1 | polypyrimidine tract binding protein 1 | Nucleus | enzyme |
| P35244 | RPA3 | replication protein A3 | Nucleus | other |
| Q9UHB6 | LIMA1 | LIM domain and actin binding 1 | Cytoplasm | other |
| P51114 | FXR1 | FMR1 autosomal homolog 1 | Cytoplasm | other |
| Q00325 | SLC25A3 | solute carrier family 25 member 3 | Cytoplasm | transporter |
| Q9Y5S9 | RBM8A | RNA binding motif protein 8A | Nucleus | other |
| Q02543 | RPL18A | ribosomal protein L18a | Cytoplasm | other |
| Q09028 | RBBP4 | RB binding protein 4, chromatin remodeling fa | Nucleus | enzyme |
| P61353 | RPL27 | ribosomal protein L27 | Cytoplasm | other |
| P46940 | IQGAP1 | IQ motif containing GTPase activating protein | Cytoplasm | other |
| O43143 | DHX15 | DEAH-box helicase 15 | Nucleus | enzyme |
| Q6P2Q9 | PRPF8 | pre-mRNA processing factor 8 | Nucleus | other |
| P04439 | HLA-A | major histocompatibility complex, class I, A | Plasma Membrane | other |
| P06396 | GSN | gelsolin | Extracellular Space | other |
| Q9UMS4 | PRPF19 | pre-mRNA processing factor 19 | Nucleus | enzyme |
| P04899 | GNAI2 | G protein subunit alpha i2 | Plasma Membrane | enzyme |
| Q9UQ35 | SRRM2 | serine/arginine repetitive matrix 2 | Nucleus | other |
| Q86U42 | PABPN1 | poly(A) binding protein nuclear 1 | Nucleus | enzyme |
| P14649 | MYL6B | myosin light chain 6B | Cytoplasm | other |
| Q13310 | PABPC4 | poly(A) binding protein cytoplasmic 4 | Cytoplasm | translation regulator |
| Q15029 | EFTUD2 | elongation factor Tu GTP binding domain con | Nucleus | enzyme |
| O43707 | ACTN4 | actinin alpha 4 | Cytoplasm | transcription regulator |
| P42224 | STAT1 | signal transducer and activator of transcriptio | Nucleus | transcription regulator |
| P62995 | TRA2B | transformer 2 beta homolog | Nucleus | other |
| Q15366 | PCBP2 | poly(rC) binding protein 2 | Nucleus | other |
| P26196 | DDX6 | DEAD-box helicase 6 | Nucleus | enzyme |
| Q15393 | SF3B3 | splicing factor 3b subunit 3 | Nucleus | other |
| Q96II8 | LRCH3 | leucine rich repeats and calponin homology d | Cytoplasm | other |
| Q15717 | ELAVL1 | ELAV like RNA binding protein 1 | Cytoplasm | other |
| P35611 | ADD1 | adducin 1 | Cytoplasm | other |
| P42167 | TMPO | thymopoietin | Nucleus | other |
| P62873 | GNB1 | G protein subunit beta 1 | Plasma Membrane | enzyme |
| Q15306 | IRF4 | Interferon regulatory factor 4 | Nucleus | |
| P27708 | CAD | carbamoyl-phosphate synthetase 2, aspartate | Cytoplasm | enzyme |
| P15927 | RPA2 | replication protein A2 | Nucleus | other |
| O94832 | MYO1D | myosin ID | Cytoplasm | enzyme |
| P63244 | RACK1 | receptor for activated C kinase 1 | Cytoplasm | enzyme |
| P47755 | CAPZA2 | capping actin protein of muscle Z-line alpha su | Cytoplasm | other |
| O75494 | SRSF10 | serine and arginine rich splicing factor 10 | Nucleus | other |
| P62140 | PPP1CB | protein phosphatase 1 catalytic subunit beta | Cytoplasm | phosphatase |
| O75531 | BANF1 | barrier to autointegration factor 1 | Nucleus | other |
| Q9Y224 | RTRAF | RNA transcription, translation and transport f | Nucleus | other |
| Q8NI27 | THOC2 | THO complex 2 | Nucleus | other |
| O00299 | CLIC1 | chloride intracellular channel 1 | Nucleus | ion channel |
| P49756 | RBM25 | RNA binding motif protein 25 | Nucleus | other |
| P60891 | PRPS1 | phosphoribosyl pyrophosphate synthetase 1 | Cytoplasm | kinase |

| | | | | |
|---|---|---|---|---|
| Q9H9B4 | SFXN1 | sideroflexin 1 | Cytoplasm | transporter |
| O14979 | HNRNPDL | heterogeneous nuclear ribonucleoprotein D li | Nucleus | other |
| P15153 | RAC2 | Rac family small GTPase 2 | Cytoplasm | enzyme |
| Q86W42 | THOC6 | THO complex 6 | Nucleus | other |
| P49327 | FASN | fatty acid synthase | Cytoplasm | enzyme |
| Q13427 | PPIG | peptidylprolyl isomerase G | Nucleus | enzyme |
| O14639 | ABLIM1 | actin binding LIM protein 1 | Cytoplasm | other |
| Q9Y3I0 | RTCB | RNA 2',3'-cyclic phosphate and 5'-OH ligase | Cytoplasm | enzyme |
| O75400 | PRPF40A | pre-mRNA processing factor 40 homolog A | Nucleus | other |
| P29350 | PTPN6 | protein tyrosine phosphatase, non-receptor t | Cytoplasm | phosphatase |
| Q8WWM7 | ATXN2L | ataxin 2 like | Nucleus | other |
| P55209 | NAP1L1 | nucleosome assembly protein 1 like 1 | Nucleus | other |
| Q9Y3U8 | RPL36 | ribosomal protein L36 | Cytoplasm | other |
| Q6WKZ4 | RAB11FIP1 | RAB11 family interacting protein 1 | Cytoplasm | other |
| P06239 | LCK | LCK proto-oncogene, Src family tyrosine kinas | Cytoplasm | kinase |
| Q9Y4I1 | MYO5A | myosin VA | Cytoplasm | enzyme |
| Q86V48 | LUZP1 | leucine zipper protein 1 | Nucleus | other |
| Q92499 | DDX1 | DEAD-box helicase 1 | Nucleus | enzyme |
| P63167 | DYNLL1 | dynein light chain LC8-type 1 | Cytoplasm | other |
| Q12906 | ILF3 | interleukin enhancer binding factor 3 | Nucleus | transcription regulator |
| P05023 | ATP1A1 | ATPase Na+/K+ transporting subunit alpha 1 | Plasma Membrane | transporter |
| Q9NR30 | DDX21 | DExD-box helicase 21 | Nucleus | enzyme |
| Q9UEY8 | ADD3 | adducin 3 | Cytoplasm | other |
| Q14764 | MVP | major vault protein | Nucleus | other |
| P18077 | RPL35A | ribosomal protein L35a | Cytoplasm | other |
| Q13573 | SNW1 | SNW domain containing 1 | Nucleus | transcription regulator |
| Q16891 | IMMT | inner membrane mitochondrial protein | Cytoplasm | other |
| P13796 | LCP1 | lymphocyte cytosolic protein 1 | Cytoplasm | other |
| Q2TAY7 | SMU1 | SMU1, DNA replication regulator and spliceos | Nucleus | other |
| Q2VIR3 | EIF2S3B | eukaryotic translation initiation factor 2 subu | Other | other |
| Q15025 | TNIP1 | TNFAIP3 interacting protein 1 | Nucleus | other |
| P08195 | SLC3A2 | solute carrier family 3 member 2 | Plasma Membrane | transporter |
| Q13247 | SRSF6 | serine and arginine rich splicing factor 6 | Nucleus | other |
| Q15459 | SF3A1 | splicing factor 3a subunit 1 | Nucleus | other |
| P52701 | MSH6 | mutS homolog 6 | Nucleus | enzyme |
| P41250 | GARS | glycyl-tRNA synthetase | Cytoplasm | enzyme |
| O43175 | PHGDH | phosphoglycerate dehydrogenase | Cytoplasm | enzyme |
| Q13283 | G3BP1 | G3BP stress granule assembly factor 1 | Nucleus | enzyme |
| Q7Z417 | NUFIP2 | NUFIP2, FMR1 interacting protein 2 | Cytoplasm | other |
| Q702N8 | XIRP1 | xin actin binding repeat containing 1 | Plasma Membrane | other |
| Q14444 | CAPRIN1 | cell cycle associated protein 1 | Plasma Membrane | other |
| P24666 | ACP1 | acid phosphatase 1 | Cytoplasm | phosphatase |
| P62879 | GNB2 | G protein subunit beta 2 | Plasma Membrane | enzyme |
| Q12874 | SF3A3 | splicing factor 3a subunit 3 | Nucleus | other |
| P21796 | VDAC1 | voltage dependent anion channel 1 | Cytoplasm | ion channel |
| P06493 | CDK1 | cyclin dependent kinase 1 | Nucleus | kinase |
| Q96FV9 | THOC1 | THO complex 1 | Nucleus | transcription regulator |
| Q9BUJ2 | HNRNPUL1 | heterogeneous nuclear ribonucleoprotein U li | Nucleus | other |
| Q13151 | HNRNPA0 | heterogeneous nuclear ribonucleoprotein A0 | Nucleus | other |
| Q9P258 | RCC2 | regulator of chromosome condensation 2 | Nucleus | other |

| | | | | |
|---|---|---|---|---|
| Q92878 | RAD50 | RAD50 double strand break repair protein | Nucleus | enzyme |
| P27348 | YWHAQ | tyrosine 3-monooxygenase/tryptophan 5-mo | Cytoplasm | other |
| Q9UKM9 | RALY | RALY heterogeneous nuclear ribonucleoprote | Nucleus | transcription regulator |
| P21580 | TNFAIP3 | TNF alpha induced protein 3 | Nucleus | enzyme |
| O75533 | SF3B1 | splicing factor 3b subunit 1 | Nucleus | other |
| Q9BY77 | POLDIP3 | DNA polymerase delta interacting protein 3 | Nucleus | other |
| P45880 | VDAC2 | voltage dependent anion channel 2 | Cytoplasm | ion channel |
| O00139 | KIF2A | kinesin family member 2A | Cytoplasm | other |
| O00148 | DDX39A | DExD-box helicase 39A | Nucleus | enzyme |
| P17987 | TCP1 | t-complex 1 | Cytoplasm | other |
| Q13769 | THOC5 | THO complex 5 | Cytoplasm | other |
| Q8NCA5 | FAM98A | family with sequence similarity 98 member A | Other | other |
| Q07955 | SRSF1 | serine and arginine rich splicing factor 1 | Nucleus | other |
| P07814 | EPRS | glutamyl-prolyl-tRNA synthetase | Cytoplasm | enzyme |
| Q14974 | KPNB1 | karyopherin subunit beta 1 | Nucleus | transporter |
| P12956 | XRCC6 | X-ray repair cross complementing 6 | Nucleus | enzyme |
| Q99459 | CDC5L | cell division cycle 5 like | Nucleus | transcription regulator |
| Q13557 | CAMK2D | calcium/calmodulin dependent protein kinase | Cytoplasm | kinase |
| P09661 | SNRPA1 | small nuclear ribonucleoprotein polypeptide A | Nucleus | other |
| P04843 | RPN1 | ribophorin I | Cytoplasm | enzyme |
| Q13951 | CBFB | core-binding factor beta subunit | Nucleus | transcription regulator |
| Q13045 | FLII | FLII, actin remodeling protein | Nucleus | other |
| Q9Y3X0 | CCDC9 | coiled-coil domain containing 9 | Extracellular Space | other |
| P31942 | HNRNPH3 | heterogeneous nuclear ribonucleoprotein H3 | Nucleus | other |
| Q9UJV9 | DDX41 | DEAD-box helicase 41 | Nucleus | enzyme |
| P51116 | FXR2 | FMR1 autosomal homolog 2 | Cytoplasm | other |
| P49368 | CCT3 | chaperonin containing TCP1 subunit 3 | Cytoplasm | other |
| P43490 | NAMPT | nicotinamide phosphoribosyltransferase | Extracellular Space | cytokine |
| P78371 | CCT2 | chaperonin containing TCP1 subunit 2 | Cytoplasm | kinase |
| P53814 | SMTN | smoothelin | Extracellular Space | other |
| Q14739 | LBR | lamin B receptor | Nucleus | enzyme |
| Q9Y3Z3 | SAMHD1 | SAM and HD domain containing deoxynucleos | Nucleus | enzyme |
| P50990 | CCT8 | chaperonin containing TCP1 subunit 8 | Cytoplasm | enzyme |
| Q6I9Y2 | THOC7 | THO complex 7 | Nucleus | other |
| Q6IS14 | EIF5AL1 | eukaryotic translation initiation factor 5A-like | Other | other |
| Q13242 | SRSF9 | serine and arginine rich splicing factor 9 | Nucleus | enzyme |
| Q9ULV4 | CORO1C | coronin 1C | Cytoplasm | other |
| Q12905 | ILF2 | interleukin enhancer binding factor 2 | Nucleus | transcription regulator |
| O95816 | BAG2 | BCL2 associated athanogene 2 | Cytoplasm | other |
| P42704 | LRPPRC | leucine rich pentatricopeptide repeat contain | Cytoplasm | other |
| P08865 | RPSA | ribosomal protein SA | Cytoplasm | translation regulator |
| O60884 | DNAJA2 | DnaJ heat shock protein family (Hsp40) memb | Nucleus | enzyme |
| O43684 | BUB3 | BUB3, mitotic checkpoint protein | Nucleus | other |
| O43390 | HNRNPR | heterogeneous nuclear ribonucleoprotein R | Nucleus | other |
| P50402 | EMD | emerin | Nucleus | other |
| Q92945 | KHSRP | KH-type splicing regulatory protein | Nucleus | enzyme |
| Q9Y230 | RUVBL2 | RuvB like AAA ATPase 2 | Nucleus | transcription regulator |
| P51149 | RAB7A | RAB7A, member RAS oncogene family | Cytoplasm | enzyme |
| P62304 | SNRPE | small nuclear ribonucleoprotein polypeptide E | Nucleus | other |
| Q02978 | SLC25A11 | solute carrier family 25 member 11 | Cytoplasm | transporter |

| | | | | |
|---|---|---|---|---|
| P36873 | PPP1CC | protein phosphatase 1 catalytic subunit gamm | Nucleus | phosphatase |
| P05161 | ISG15 | ISG15 ubiquitin-like modifier | Extracellular Space | other |
| P36542 | ATP5F1C | ATP synthase F1 subunit gamma | Cytoplasm | transporter |
| O75369 | FLNB | filamin B | Cytoplasm | other |
| P49458 | SRP9 | signal recognition particle 9 | Cytoplasm | other |
| P17980 | PSMC3 | proteasome 26S subunit, ATPase 3 | Nucleus | enzyme |
| P55769 | SNU13 | small nuclear ribonucleoprotein 13 | Nucleus | other |
| Q15007 | WTAP | WT1 associated protein | Nucleus | other |
| Q9UBM7 | DHCR7 | 7-dehydrocholesterol reductase | Cytoplasm | enzyme |
| P16615 | ATP2A2 | ATPase sarcoplasmic/endoplasmic reticulum ( | Cytoplasm | transporter |
| P40227 | CCT6A | chaperonin containing TCP1 subunit 6A | Cytoplasm | other |
| P78527 | PRKDC | protein kinase, DNA-activated, catalytic polyp | Nucleus | kinase |
| Q96J01 | THOC3 | THO complex 3 | Nucleus | other |
| P02786 | TFRC | transferrin receptor | Plasma Membrane | transporter |
| P20700 | LMNB1 | lamin B1 | Nucleus | other |
| Q13761 | RUNX3 | Runt Related Transcription Factor 3 | Nucleus | |
| P50991 | CCT4 | chaperonin containing TCP1 subunit 4 | Cytoplasm | other |
| Q69YQ0 | SPECC1L | sperm antigen with calponin homology and co | Extracellular Space | other |
| Q15637 | SF1 | splicing factor 1 | Nucleus | transcription regulator |
| Q6UN15 | FIP1L1 | factor interacting with PAPOLA and CPSF1 | Nucleus | other |
| Q8IYB3 | SRRM1 | serine and arginine repetitive matrix 1 | Nucleus | other |
| Q9Y3F4 | STRAP | serine/threonine kinase receptor associated p | Plasma Membrane | other |
| P62310 | LSM3 | LSM3 homolog, U6 small nuclear RNA and mR | Nucleus | other |
| Q96SB3 | PPP1R9B | protein phosphatase 1 regulatory subunit 9B | Cytoplasm | enzyme |
| P61026 | RAB10 | RAB10, member RAS oncogene family | Cytoplasm | enzyme |
| Q92888 | ARHGEF1 | Rho guanine nucleotide exchange factor 1 | Cytoplasm | other |
| P48444 | ARCN1 | archain 1 | Cytoplasm | other |
| Q9UPN4 | CEP131 | centrosomal protein 131 | Cytoplasm | other |
| Q9BWJ5 | SF3B5 | splicing factor 3b subunit 5 | Nucleus | other |
| Q9H3U1 | UNC45A | unc-45 myosin chaperone A | Plasma Membrane | other |
| P61158 | ACTR3 | ARP3 actin related protein 3 homolog | Plasma Membrane | other |
| Q9Y613 | FHOD1 | formin homology 2 domain containing 1 | Cytoplasm | other |
| P53007 | SLC25A1 | solute carrier family 25 member 1 | Other | transporter |
| O75569 | PRKRA | protein activator of interferon induced protei | Cytoplasm | other |
| Q9NX63 | CHCHD3 | coiled-coil-helix-coiled-coil-helix domain cont | Cytoplasm | transcription regulator |
| P14625 | HSP90B1 | heat shock protein 90 beta family member 1 | Cytoplasm | other |
| P56134 | ATP5MF | ATP synthase membrane subunit f | Cytoplasm | transporter |
| Q53EZ4 | CEP55 | centrosomal protein 55 | Cytoplasm | other |
| Q9NW64 | RBM22 | RNA binding motif protein 22 | Nucleus | other |
| P22087 | FBL | fibrillarin | Nucleus | enzyme |
| Q16531 | DDB1 | damage specific DNA binding protein 1 | Nucleus | other |
| P61163 | ACTR1A | ARP1 actin related protein 1 homolog A | Cytoplasm | other |
| P61160 | ACTR2 | ARP2 actin related protein 2 homolog | Plasma Membrane | other |
| Q5SW79 | CEP170 | centrosomal protein 170 | Nucleus | other |
| Q9P2I0 | CPSF2 | cleavage and polyadenylation specific factor 2 | Nucleus | other |
| P39656 | DDOST | dolichyl-diphosphooligosaccharide--protein g | Cytoplasm | enzyme |
| O94925 | GLS | glutaminase | Cytoplasm | enzyme |
| Q9Y265 | RUVBL1 | RuvB like AAA ATPase 1 | Nucleus | transcription regulator |
| Q3MHD2 | LSM12 | LSM12 homolog | Other | other |
| P62333 | PSMC6 | proteasome 26S subunit, ATPase 6 | Nucleus | peptidase |

| | | | | |
|---|---|---|---|---|
| P62306 | SNRPF | small nuclear ribonucleoprotein polypeptide F | Nucleus | other |
| Q8NAV1 | PRPF38A | pre-mRNA processing factor 38A | Nucleus | other |
| P22695 | UQCRC2 | ubiquinol-cytochrome c reductase core prote | Cytoplasm | enzyme |
| P48643 | CCT5 | chaperonin containing TCP1 subunit 5 | Cytoplasm | other |
| P30504 | HLA-C | major histocompatibility complex, class I, C | Plasma Membrane | other |
| Q9Y2D5 | AKAP2 | A-kinase anchoring protein 2 | Plasma Membrane | other |
| Q13561 | DCTN2 | dynactin subunit 2 | Cytoplasm | other |
| P35249 | RFC4 | replication factor C subunit 4 | Nucleus | other |
| Q7RTV0 | PHF5A | PHD finger protein 5A | Nucleus | transcription regulator |
| P53618 | COPB1 | coatomer protein complex subunit beta 1 | Cytoplasm | transporter |
| Q6PKG0 | LARP1 | La ribonucleoprotein domain family member | Cytoplasm | translation regulator |
| Q06787 | FMR1 | fragile X mental retardation 1 | Cytoplasm | translation regulator |
| Q13123 | IK | IK cytokine | Extracellular Space | cytokine |
| P51571 | SSR4 | signal sequence receptor subunit 4 | Cytoplasm | other |
| Q9UN86 | G3BP2 | G3BP stress granule assembly factor 2 | Cytoplasm | enzyme |
| P14868 | DARS | aspartyl-tRNA synthetase | Cytoplasm | enzyme |
| Q9H223 | EHD4 | EH domain containing 4 | Plasma Membrane | enzyme |
| Q14257 | RCN2 | reticulocalbin 2 | Cytoplasm | other |
| Q99661 | KIF2C | kinesin family member 2C | Nucleus | other |
| Q9NZ01 | TECR | trans-2,3-enoyl-CoA reductase | Plasma Membrane | enzyme |
| Q9P0L0 | VAPA | VAMP associated protein A | Plasma Membrane | other |
| Q10570 | CPSF1 | cleavage and polyadenylation specific factor 1 | Nucleus | other |
| Q13148 | TARDBP | TAR DNA binding protein | Nucleus | transcription regulator |
| P20592 | MX2 | MX dynamin like GTPase 2 | Nucleus | enzyme |
| O14579 | COPE | coatomer protein complex subunit epsilon | Cytoplasm | transporter |
| Q15428 | SF3A2 | splicing factor 3a subunit 2 | Nucleus | other |
| Q01196 | RUNX1 | runt related transcription factor 1 | Nucleus | transcription regulator |
| Q99623 | PHB2 | prohibitin 2 | Cytoplasm | transcription regulator |
| Q7Z6E9 | RBBP6 | RB binding protein 6, ubiquitin ligase | Nucleus | enzyme |
| Q02040 | AKAP17A | A-kinase anchoring protein 17A | Nucleus | other |
| P02794 | FTH1 | ferritin heavy chain 1 | Cytoplasm | enzyme |
| Q96DI7 | SNRNP40 | small nuclear ribonucleoprotein U5 subunit 40 | Nucleus | other |
| O15143 | ARPC1B | actin related protein 2/3 complex subunit 1B | Cytoplasm | other |
| P29692 | EEF1D | eukaryotic translation elongation factor 1 delt | Cytoplasm | translation regulator |
| Q9UPN7 | PPP6R1 | protein phosphatase 6 regulatory subunit 1 | Cytoplasm | other |
| P09382 | LGALS1 | galectin 1 | Extracellular Space | other |
| Q96FS4 | SIPA1 | signal-induced proliferation-associated 1 | Cytoplasm | other |
| Q9Y608 | LRRFIP2 | LRR binding FLII interacting protein 2 | Extracellular Space | other |
| O15042 | U2SURP | U2 snRNP associated SURP domain containing | Nucleus | other |
| Q96C19 | EFHD2 | EF-hand domain family member D2 | Other | other |
| P28066 | PSMA5 | proteasome subunit alpha 5 | Cytoplasm | peptidase |
| Q13077 | TRAF1 | TNF receptor associated factor 1 | Cytoplasm | other |
| P08574 | CYC1 | cytochrome c1 | Cytoplasm | enzyme |
| Q9NTJ3 | SMC4 | structural maintenance of chromosomes 4 | Nucleus | transporter |
| O00165 | HAX1 | HCLS1 associated protein X-1 | Cytoplasm | other |
| Q9NZR1 | TMOD2 | tropomodulin 2 | Cytoplasm | other |
| O00743 | PPP6C | protein phosphatase 6 catalytic subunit | Nucleus | phosphatase |
| P52732 | KIF11 | kinesin family member 11 | Nucleus | other |
| P24534 | EEF1B2 | eukaryotic translation elongation factor 1 bet | Cytoplasm | translation regulator |
| Q9H7N4 | SCAF1 | SR-related CTD associated factor 1 | Extracellular Space | other |

| | | | | |
|---|---|---|---|---|
| Q13595 | TRA2A | transformer 2 alpha homolog | Nucleus | other |
| P78406 | RAE1 | ribonucleic acid export 1 | Nucleus | other |
| Q00653 | NFKB2 | nuclear factor kappa B subunit 2 | Nucleus | transcription regulator |
| Q1KMD3 | HNRNPUL2 | heterogeneous nuclear ribonucleoprotein U li | Nucleus | other |
| P41240 | CSK | C-terminal Src kinase | Cytoplasm | kinase |
| P48047 | ATP5PO | ATP synthase peripheral stalk subunit OSCP | Cytoplasm | transporter |
| P61221 | ABCE1 | ATP binding cassette subfamily E member 1 | Cytoplasm | transporter |
| Q9Y5M8 | SRPRB | SRP receptor beta subunit | Cytoplasm | other |
| Q99832 | CCT7 | chaperonin containing TCP1 subunit 7 | Cytoplasm | other |
| P31939 | ATIC | 5-aminoimidazole-4-carboxamide ribonucleot | Cytoplasm | enzyme |
| Q53GQ0 | HSD17B12 | hydroxysteroid 17-beta dehydrogenase 12 | Cytoplasm | enzyme |
| O00214 | LGALS8 | galectin 8 | Extracellular Space | other |
| P40926 | MDH2 | malate dehydrogenase 2 | Cytoplasm | enzyme |
| P08579 | SNRPB2 | small nuclear ribonucleoprotein polypeptide [illegible] | Nucleus | other |
| Q02818 | NUCB1 | nucleobindin 1 | Cytoplasm | other |
| Q13523 | PRPF4B | pre-mRNA processing factor 4B | Nucleus | kinase |
| O95864 | FADS2 | fatty acid desaturase 2 | Plasma Membrane | enzyme |
| O60306 | AQR | aquarius intron-binding spliceosomal factor | Nucleus | other |
| P11387 | TOP1 | DNA topoisomerase I | Nucleus | enzyme |
| Q96T37 | RBM15 | RNA binding motif protein 15 | Nucleus | other |
| P08754 | GNAI3 | G protein subunit alpha i3 | Cytoplasm | enzyme |
| Q14699 | RFTN1 | raftlin, lipid raft linker 1 | Plasma Membrane | other |
| Q8TBC3 | SHKBP1 | SH3KBP1 binding protein 1 | Other | other |
| P24539 | ATP5PB | ATP synthase peripheral stalk-membrane sub | Cytoplasm | transporter |
| Q9Y305 | ACOT9 | acyl-CoA thioesterase 9 | Cytoplasm | enzyme |
| Q8IWX8 | CHERP | calcium homeostasis endoplasmic reticulum p | Cytoplasm | other |
| P46087 | NOP2 | NOP2 nucleolar protein | Nucleus | other |
| P34897 | SHMT2 | serine hydroxymethyltransferase 2 | Cytoplasm | enzyme |
| Q15427 | SF3B4 | splicing factor 3b subunit 4 | Nucleus | other |
| Q15125 | EBP | emopamil binding protein (sterol isomerase) | Cytoplasm | enzyme |
| Q63HN8 | RNF213 | ring finger protein 213 | Cytoplasm | enzyme |
| P63162 | SNRPN | small nuclear ribonucleoprotein polypeptide [illegible] | Nucleus | other |
| P38159 | RBMX | RNA binding motif protein, X-linked | Nucleus | other |
| Q8N684 | CPSF7 | cleavage and polyadenylation specific factor 7 | Nucleus | other |
| P00338 | LDHA | lactate dehydrogenase A | Cytoplasm | enzyme |
| Q13155 | AIMP2 | aminoacyl tRNA synthetase complex interacti | Plasma Membrane | other |
