## Supplementary Table 1C for "Protein Interactome of the Cancerous Inhibitor of Protein Phosphatase 2A (CIP2A) in Th17 Cells"

**Supplementry Table 1C:** Protein Scores, peptides detected and sequence coverage from the MaxQuant/Andromeda analysis

The data is filtered to only include proteins with SAINT SP values of > 0.95 and less than 60% representation in the CRAPOME

| Majority protein IDs | Protein names | Peptides | Razor + unique peptides | Unique peptides | Sequence coverage [%] | Unique + razor sequence coverage [%] | Unique sequence coverage [%] | Mol. weight [kDa] | Q-value | Score | Raw Intensity | MS/MS count |
|---|---|---|---|---|---|---|---|---|---|---|---|---|
| B0I1T2 | Unconventional myosin-Ig;Minor histocompatibility antigen HA-2 | 78 | 78 | 75 | 81 | 81 | 81 | 116.44 | 0 | 323.31 | 97663000000 | 1828 |
| Q8TCG1 | Protein CIP2A | 82 | 82 | 82 | 73 | 73 | 73 | 102.18 | 0 | 323.31 | 60687000000 | 1280 |
| P11940 | Polyadenylate-binding protein 1 | 36 | 36 | 22 | 61.8 | 61.8 | 40.6 | 70.67 | 0 | 323.31 | 12393000000 | 432 |
| Q9NVI7 | ATPase family AAA domain-containing protein 3A | 47 | 47 | 14 | 56.3 | 56.3 | 24 | 71.368 | 0 | 323.31 | 10900000000 | 513 |
| Q9NVI7 | ATPase family AAA domain-containing protein 3A | 47 | 47 | 14 | 56.3 | 56.3 | 24 | 71.368 | 0 | 323.31 | 10900000000 | 513 |
| P19474 | E3 ubiquitin-protein ligase TRIM21 | 29 | 29 | 29 | 58.3 | 58.3 | 58.3 | 54.169 | 0 | 323.31 | 10423000000 | 293 |
| P35580 | Myosin-10 | 113 | 80 | 74 | 58.4 | 46.9 | 43.4 | 229 | 0 | 323.31 | 7833800000 | 596 |
| Q01082 | Spectrin beta chain, non-erythrocytic 1 | 90 | 90 | 83 | 44.5 | 44.5 | 41.5 | 274.61 | 0 | 323.31 | 7520600000 | 535 |
| P07910 | Heterogeneous nuclear ribonucleoproteins C1/C2 | 16 | 16 | 16 | 46.1 | 46.1 | 46.1 | 33.67 | 0 | 189.95 | 7158400000 | 263 |
| Q8NF50 | Dedicator of cytokinesis protein 8 | 70 | 70 | 68 | 39.1 | 39.1 | 37.9 | 238.53 | 0 | 323.31 | 7318000000 | 499 |
| Q15149 | Plectin | 152 | 152 | 148 | 37.2 | 37.2 | 36.1 | 531.78 | 0 | 323.31 | 6856800000 | 697 |
| Q9UKV3 | Apoptotic chromatin condensation inducer in the nucleus | 39 | 39 | 39 | 31.5 | 31.5 | 31.5 | 151.86 | 0 | 323.31 | 5773000000 | 375 |
| Q9NYL9 | Tropomodulin-3 | 16 | 16 | 15 | 65.6 | 65.6 | 63.1 | 39.594 | 0 | 310.46 | 5462200000 | 216 |
| P0DMV9;P0DMV8 | Heat shock 70 kDa protein 1B;Heat shock 70 kDa protein 1A | 35 | 35 | 20 | 55.4 | 55.4 | 38.4 | 70.051 | 0 | 323.31 | 4676800000 | 230 |
| Q6NYC8 | Phostensin | 25 | 25 | 25 | 49.3 | 49.3 | 49.3 | 67.942 | 0 | 323.31 | 4337700000 | 213 |
| Q9H307 | Pinin | 28 | 28 | 28 | 29.3 | 29.3 | 29.3 | 81.613 | 0 | 268.65 | 4345600000 | 270 |
| Q7Z2W4 | Zinc finger CCCH-type antiviral protein 1 | 39 | 39 | 39 | 58.6 | 58.6 | 58.6 | 101.43 | 0 | 323.31 | 4490100000 | 302 |
| Q13428 | Treacle protein | 42 | 42 | 42 | 31.2 | 31.2 | 31.2 | 152.1 | 0 | 323.31 | 4354500000 | 321 |
| P52907 | F-actin-capping protein subunit alpha-1 | 16 | 16 | 13 | 65 | 65 | 56.6 | 32.922 | 0 | 229.2 | 4010200000 | 153 |
| Q7Z406 | Myosin-14 | 18 | 2 | 2 | 8.1 | 0.8 | 0.8 | 227.87 | 0 | 55.18 | 3772100000 | 60 |
| O43795 | Unconventional myosin-Ib | 30 | 30 | 28 | 32 | 32 | 30.2 | 131.98 | 0 | 323.31 | 3542800000 | 215 |

| | | | | | | | | | | | | |
|---|---|---|---|---|---|---|---|---|---|---|---|---|
| P38919 | Eukaryotic initiation factor 4A-III;Eukaryotic initiation factor 4A-III, N-terminally processed | 25 | 25 | 23 | 57.9 | 57.9 | 52.1 | 46.871 | 0 | 314.35 | 3630800000 | 250 |
| O00159 | Unconventional myosin-Ic | 40 | 40 | 40 | 44.6 | 44.6 | 44.6 | 121.68 | 0 | 323.31 | 3348700000 | 220 |
| Q9Y2W1 | Thyroid hormone receptor-associated protein 3 | 27 | 27 | 26 | 27 | 27 | 27 | 108.66 | 0 | 297.82 | 3233300000 | 258 |
| Q92614 | Unconventional myosin-XVIIIa | 42 | 42 | 42 | 26.8 | 26.8 | 26.8 | 233.11 | 0 | 323.31 | 3213800000 | 270 |
| P62136 | Serine/threonine-protein phosphatase PP1-alpha catalytic subunit | 20 | 20 | 4 | 59.4 | 59.4 | 16.1 | 37.512 | 0 | 293.35 | 2842400000 | 177 |
| P14866 | Heterogeneous nuclear ribonucleoprotein L | 18 | 18 | 18 | 51.8 | 51.8 | 51.8 | 64.132 | 0 | 316.73 | 3016000000 | 169 |
| P47756 | F-actin-capping protein subunit beta | 16 | 16 | 16 | 60.3 | 60.3 | 60.3 | 31.35 | 0 | 242.7 | 2901700000 | 183 |
| Q9NYF8 | Bcl-2-associated transcription factor 1 | 25 | 24 | 24 | 28.5 | 28.5 | 28.5 | 106.12 | 0 | 214.76 | 2837300000 | 207 |
| O14974 | Protein phosphatase 1 regulatory subunit 12A | 27 | 27 | 27 | 29.1 | 29.1 | 29.1 | 115.28 | 0 | 323.31 | 2298300000 | 177 |
| Q6P2E9 | Enhancer of mRNA-decapping protein 4 | 33 | 33 | 33 | 34.3 | 34.3 | 34.3 | 151.66 | 0 | 323.31 | 2067700000 | 168 |
| P12268 | Inosine-5-monophosphate dehydrogenase 2 | 24 | 24 | 24 | 55.1 | 55.1 | 55.1 | 55.804 | 0 | 228.84 | 2406600000 | 129 |
| Q5T9A4 | ATPase family AAA domain-containing protein 3B | 42 | 15 | 12 | 57.7 | 29.9 | 20.7 | 72.572 | 0 | 184.97 | 2177800000 | 109 |
| Q96HS1 | Serine/threonine-protein phosphatase PGAM5, mitochondrial | 18 | 18 | 18 | 65.1 | 65.1 | 65.1 | 32.004 | 0 | 130.9 | 2058800000 | 138 |
| P62906 | 60S ribosomal protein L10a | 10 | 10 | 10 | 46.1 | 46.1 | 46.1 | 24.831 | 0 | 133.48 | 1997100000 | 119 |
| O00422 | Histone deacetylase complex subunit SAP18 | 14 | 14 | 14 | 84.3 | 84.3 | 84.3 | 17.561 | 0 | 167.19 | 2061600000 | 128 |
| Q02878 | 60S ribosomal protein L6 | 18 | 18 | 18 | 53.1 | 53.1 | 53.1 | 32.728 | 0 | 149.03 | 2161700000 | 122 |
| Q9Y6N5 | Sulfide:quinone oxidoreductase, mitochondrial | 14 | 14 | 14 | 44.7 | 44.7 | 44.7 | 49.96 | 0 | 223.4 | 1929000000 | 96 |
| P43243 | Matrin-3 | 21 | 21 | 21 | 31.8 | 31.8 | 31.8 | 94.622 | 0 | 323.31 | 2025300000 | 164 |
| P04114 | Apolipoprotein B-100;Apolipoprotein B-48 | 11 | 11 | 11 | 2.2 | 2.2 | 2.2 | 515.6 | 0 | 67.649 | 1911400000 | 29 |
| P84103 | Serine/arginine-rich splicing factor 3 | 7 | 7 | 7 | 44.5 | 44.5 | 44.5 | 19.329 | 0 | 71.926 | 1817700000 | 63 |
| Q92804 | TATA-binding protein-associated factor 2N | 6 | 6 | 4 | 14.7 | 14.7 | 12.3 | 61.829 | 0 | 94.242 | 1831100000 | 84 |
| P50914 | 60S ribosomal protein L14 | 8 | 8 | 8 | 36.3 | 36.3 | 36.3 | 23.432 | 0 | 72.275 | 1923000000 | 85 |
| P62829 | 60S ribosomal protein L23 | 7 | 7 | 7 | 52.9 | 52.9 | 52.9 | 14.865 | 0 | 83.177 | 2527500000 | 113 |
| Q15154 | Pericentriolar material 1 protein | 35 | 35 | 35 | 22.6 | 22.6 | 22.6 | 228.53 | 0 | 323.31 | 1592300000 | 194 |
| P27694 | Replication protein A 70 kDa DNA-binding subunit;Replication protein A 70 kDa DNA-binding subunit, N-terminally processed | 17 | 17 | 17 | 38.8 | 38.8 | 38.8 | 68.137 | 0 | 241.95 | 1667300000 | 126 |

| | | | | | | | | | | | | |
|---|---|---|---|---|---|---|---|---|---|---|---|---|
| Q16666 | Gamma-interferon-inducible protein 16 | 25 | 25 | 25 | 40.5 | 40.5 | 40.5 | 88.255 | 0 | 217.5 | 1640200000 | 171 |
| P06753 | Tropomyosin alpha-3 chain | 19 | 10 | 10 | 51.9 | 34.7 | 34.7 | 32.95 | 0 | 131.33 | 1634500000 | 114 |
| O75643 | U5 small nuclear ribonucleoprotein 200 kDa helicase | 36 | 36 | 36 | 20.4 | 20.4 | 20.4 | 244.5 | 0 | 323.31 | 1494600000 | 143 |
| P26599 | Polypyrimidine tract-binding protein 1 | 16 | 16 | 16 | 45 | 45 | 45 | 57.221 | 0 | 187.85 | 1699600000 | 104 |
| P35244 | Replication protein A 14 kDa subunit | 9 | 9 | 9 | 89.3 | 89.3 | 89.3 | 13.569 | 0 | 98.381 | 1556100000 | 36 |
| Q9UHB6 | LIM domain and actin-binding protein 1 | 19 | 19 | 19 | 33.3 | 33.3 | 33.3 | 85.225 | 0 | 175.01 | 1435600000 | 94 |
| P51114 | Fragile X mental retardation syndrome-related protein 1 | 18 | 18 | 16 | 36.1 | 36.1 | 32.7 | 69.72 | 0 | 209.51 | 1478900000 | 102 |
| Q00325 | Phosphate carrier protein, mitochondrial | 10 | 10 | 10 | 30.1 | 30.1 | 30.1 | 40.094 | 0 | 85.688 | 1471900000 | 99 |
| Q9Y5S9 | RNA-binding protein 8A | 8 | 8 | 8 | 77.6 | 77.6 | 77.6 | 19.889 | 0 | 209.18 | 1474800000 | 66 |
| Q02543 | 60S ribosomal protein L18a | 14 | 14 | 14 | 58.5 | 58.5 | 58.5 | 20.762 | 0 | 115.39 | 1445800000 | 140 |
| Q08211 | ATP-dependent RNA helicase A | 26 | 26 | 26 | 24.3 | 24.3 | 24.3 | 140.96 | 0 | 214.19 | 1330600000 | 103 |
| P61353 | 60S ribosomal protein L27 | 7 | 7 | 7 | 54.4 | 54.4 | 54.4 | 15.798 | 0 | 54.62 | 1379000000 | 92 |
| P46940 | Ras GTPase-activating-like protein IQGAP1 | 25 | 25 | 25 | 19.8 | 19.8 | 19.8 | 189.25 | 0 | 223.2 | 1252200000 | 139 |
| O43143 | Pre-mRNA-splicing factor ATP-dependent RNA helicase DHX15 | 20 | 20 | 20 | 28.1 | 28.1 | 28.1 | 90.932 | 0 | 157.41 | 1247100000 | 100 |
| Q6P2Q9 | Pre-mRNA-processing-splicing factor 8 | 30 | 30 | 30 | 15.4 | 15.4 | 15.4 | 273.6 | 0 | 253.35 | 1135000000 | 133 |
| P04406 | Glyceraldehyde-3-phosphate dehydrogenase | 9 | 9 | 9 | 38.5 | 38.5 | 38.5 | 36.053 | 0 | 172.21 | 1372200000 | 97 |
| P06396 | Gelsolin | 15 | 15 | 7 | 26.1 | 26.1 | 18.4 | 85.696 | 0 | 123.84 | 1215400000 | 67 |
| Q9UMS4 | Pre-mRNA-processing factor 19 | 10 | 10 | 10 | 29.6 | 29.6 | 29.6 | 55.18 | 0 | 114.91 | 994020000 | 57 |
| P04899 | Guanine nucleotide-binding protein G(i) subunit alpha-2 | 11 | 11 | 9 | 45.6 | 45.6 | 38.9 | 40.45 | 0 | 129.29 | 1060500000 | 81 |
| Q9UQ35 | Serine/arginine repetitive matrix protein 2 | 19 | 19 | 19 | 9.8 | 9.8 | 9.8 | 299.61 | 0 | 222.75 | 1025600000 | 123 |
| Q86U42 | Polyadenylate-binding protein 2 | 5 | 5 | 5 | 20.3 | 20.3 | 20.3 | 32.749 | 0 | 104.23 | 972010000 | 62 |
| P14649 | Myosin light chain 6B | 10 | 8 | 8 | 55.8 | 44.2 | 44.2 | 22.764 | 0 | 130.61 | 1055500000 | 66 |
| Q13310 | Polyadenylate-binding protein 4 | 23 | 9 | 9 | 38.7 | 17.7 | 17.7 | 70.782 | 0 | 85.756 | 935250000 | 82 |
| Q15029 | 116 kDa U5 small nuclear ribonucleoprotein component | 19 | 19 | 19 | 23.4 | 23.4 | 23.4 | 109.43 | 0 | 221.09 | 893230000 | 62 |
| O43707 | Alpha-actinin-4 | 19 | 19 | 9 | 26.7 | 26.7 | 14.7 | 104.85 | 0 | 196.34 | 899650000 | 97 |
| P42224 | Signal transducer and activator of transcription 1-alpha/beta | 16 | 16 | 16 | 26.7 | 26.7 | 26.7 | 87.334 | 0 | 150.28 | 918860000 | 65 |
| P62995 | Transformer-2 protein homolog beta | 10 | 10 | 9 | 35.4 | 35.4 | 31.9 | 33.665 | 0 | 78.421 | 852790000 | 86 |
| Q15366 | Poly(rC)-binding protein 2 | 7 | 5 | 5 | 31.8 | 23.3 | 23.3 | 38.58 | 0 | 63.079 | 834510000 | 55 |
| P26196 | Probable ATP-dependent RNA helicase DDX6 | 15 | 15 | 15 | 48.4 | 48.4 | 48.4 | 54.416 | 0 | 169.77 | 767600000 | 59 |

| | | | | | | | | | | | | |
|---|---|---|---|---|---|---|---|---|---|---|---|---|
| Q15393 | Splicing factor 3B subunit 3 | 14 | 14 | 14 | 14.9 | 14.9 | 14.9 | 135.58 | 0 | 122.05 | 781370000 | 73 |
| Q96II8 | Leucine-rich repeat and calponin homology domain-containing protein 3 | 12 | 12 | 12 | 23.8 | 23.8 | 23.8 | 86.082 | 0 | 98.352 | 794360000 | 54 |
| Q15717 | ELAV-like protein 1 | 13 | 13 | 13 | 33.4 | 33.4 | 33.4 | 36.091 | 0 | 108.33 | 756520000 | 73 |
| P35611 | Alpha-adducin | 14 | 14 | 14 | 34.5 | 34.5 | 34.5 | 80.954 | 0 | 229.85 | 781850000 | 75 |
| P42167 | Lamina-associated polypeptide 2, isoforms beta/gamma;Thymopoietin;Thymopentin | 11 | 11 | 4 | 31.5 | 31.5 | 10.6 | 50.67 | 0 | 128.36 | 784320000 | 54 |
| P62873 | Guanine nucleotide-binding protein G(I)/G(S)/G(T) subunit beta-1 | 10 | 10 | 4 | 42.1 | 42.1 | 20.9 | 37.377 | 0 | 121.74 | 794750000 | 46 |
| Q15306 | Interferon regulatory factor 4 | 10 | 10 | 10 | 32.2 | 32.2 | 32.2 | 51.772 | 0 | 186.7 | 752150000 | 57 |
| P27708 | CAD protein;Glutamine-dependent carbamoyl-phosphate synthase;Aspartate carbamoyltransferase;Dihydroorotase | 18 | 18 | 18 | 10.5 | 10.5 | 10.5 | 242.98 | 0 | 185.74 | 768880000 | 80 |
| P15927 | Replication protein A 32 kDa subunit | 7 | 7 | 7 | 43.3 | 43.3 | 43.3 | 29.247 | 0 | 116.95 | 866580000 | 59 |
| O94832 | Unconventional myosin-Id | 22 | 19 | 19 | 24.9 | 23 | 23 | 116.2 | 0 | 162.42 | 781060000 | 72 |
| P63244 | Guanine nucleotide-binding protein subunit beta-2-like 1;Guanine nucleotide-binding protein subunit beta-2-like 1, N-terminally processed | 12 | 12 | 12 | 42.3 | 42.3 | 42.3 | 35.076 | 0 | 90.603 | 736670000 | 58 |
| P47755 | F-actin-capping protein subunit alpha-2 | 7 | 4 | 4 | 28.7 | 20.3 | 20.3 | 32.949 | 0 | 56.94 | 719340000 | 28 |
| O75494 | Serine/arginine-rich splicing factor 10 | 4 | 4 | 4 | 23.7 | 23.7 | 23.7 | 31.3 | 0 | 44.975 | 707120000 | 30 |
| P62140 | Serine/threonine-protein phosphatase PP1-beta catalytic subunit | 17 | 4 | 4 | 51.4 | 15.6 | 15.6 | 37.186 | 0 | 60.879 | 709760000 | 47 |
| O75531 | Barrier-to-autointegration factor;Barrier-to-autointegration factor, N-terminally processed | 6 | 6 | 6 | 67.4 | 67.4 | 67.4 | 10.058 | 0 | 47.847 | 814010000 | 34 |
| Q9Y224 | UPF0568 protein C14orf166 | 13 | 13 | 13 | 66.8 | 66.8 | 66.8 | 28.068 | 0 | 142.92 | 688470000 | 46 |
| Q8NI27 | THO complex subunit 2 | 14 | 14 | 14 | 12.1 | 12.1 | 12.1 | 182.77 | 0 | 106.91 | 682480000 | 44 |
| O00299 | Chloride intracellular channel protein 1 | 10 | 10 | 10 | 52.3 | 52.3 | 52.3 | 26.922 | 0 | 106.9 | 663420000 | 48 |
| P49756 | RNA-binding protein 25 | 14 | 14 | 14 | 17.6 | 17.6 | 17.6 | 100.18 | 0 | 101.75 | 680700000 | 81 |
| P60866 | 40S ribosomal protein S20 | 5 | 5 | 5 | 37.8 | 37.8 | 37.8 | 13.373 | 0 | 36.97 | 732290000 | 50 |
| Q9H9B4 | Sideroflexin-1 | 8 | 8 | 8 | 23.6 | 23.6 | 23.6 | 35.619 | 0 | 75.472 | 647270000 | 46 |
| O14979 | Heterogeneous nuclear ribonucleoprotein D-like | 7 | 5 | 5 | 15 | 12.6 | 12.6 | 46.437 | 0 | 37.705 | 625390000 | 36 |
| P15153 | Ras-related C3 botulinum toxin substrate 2 | 9 | 9 | 8 | 52.6 | 52.6 | 46.9 | 21.429 | 0 | 69.427 | 633370000 | 38 |
| Q86W42 | THO complex subunit 6 homolog | 10 | 10 | 10 | 38.7 | 38.7 | 38.7 | 37.535 | 0 | 102.72 | 605870000 | 39 |

| | | | | | | | | | | | | |
|---|---|---|---|---|---|---|---|---|---|---|---|---|
| P49327 | Fatty acid synthase;[Acyl-carrier-protein] S-acetyltransferase;[Acyl-carrier-protein] S-malonyltransferase;3-oxoacyl-[acyl-carrier-protein] synthase;3-oxoacyl-[acyl-carrier-protein] reductase;3-hydroxyacyl-[acyl-carrier-protein] dehydratase;Enoyl-[acyl-carrier-protein] reductase;Oleoyl-[acyl-carrier-protein] hydrolase | 18 | 18 | 18 | 9.2 | 9.2 | 9.2 | 273.42 | 0 | 166.18 | 622170000 | 69 |
| Q13427 | Peptidyl-prolyl cis-trans isomerase G | 8 | 8 | 8 | 12.3 | 12.3 | 12.3 | 88.616 | 0 | 95.76 | 572800000 | 51 |
| O14639 | Actin-binding LIM protein 1 | 10 | 10 | 10 | 17.7 | 17.7 | 17.7 | 87.687 | 0 | 103.64 | 589540000 | 46 |
| Q9Y3I0 | tRNA-splicing ligase RtcB homolog | 9 | 9 | 9 | 23.4 | 23.4 | 23.4 | 55.21 | 0 | 94.028 | 529040000 | 43 |
| O75400 | Pre-mRNA-processing factor 40 homolog A | 10 | 10 | 10 | 11.5 | 11.5 | 11.5 | 108.8 | 0 | 70.615 | 533270000 | 60 |
| P29350 | Tyrosine-protein phosphatase non-receptor type 6 | 12 | 12 | 12 | 27.2 | 27.2 | 27.2 | 67.56 | 0 | 119.66 | 507730000 | 47 |
| Q8WWM7 | Ataxin-2-like protein | 12 | 12 | 12 | 13.2 | 13.2 | 13.2 | 113.37 | 0 | 88.802 | 483710000 | 59 |
| P55209 | Nucleosome assembly protein 1-like 1 | 4 | 4 | 4 | 14.8 | 14.8 | 14.8 | 45.374 | 0 | 53.675 | 483090000 | 19 |
| Q9Y3U8 | 60S ribosomal protein L36 | 5 | 5 | 5 | 31.4 | 31.4 | 31.4 | 12.254 | 0 | 53.56 | 510590000 | 46 |
| Q6WKZ4 | Rab11 family-interacting protein 1 | 14 | 14 | 14 | 15.7 | 15.7 | 15.7 | 137.17 | 0 | 134.39 | 444210000 | 45 |
| P06239 | Tyrosine-protein kinase Lck | 10 | 10 | 9 | 25.5 | 25.5 | 23.6 | 58 | 0 | 90.464 | 453360000 | 46 |
| Q9Y4I1 | Unconventional myosin-Va | 16 | 16 | 15 | 11 | 11 | 10.2 | 215.4 | 0 | 122.25 | 442230000 | 44 |
| Q86V48 | Leucine zipper protein 1 | 20 | 20 | 20 | 22.1 | 22.1 | 22.1 | 120.27 | 0 | 141.51 | 519640000 | 73 |
| Q92499 | ATP-dependent RNA helicase DDX1 | 13 | 13 | 13 | 21.5 | 21.5 | 21.5 | 82.431 | 0 | 145.51 | 465420000 | 43 |
| P63162;P14678 | Small nuclear ribonucleoprotein-associated protein N;Small nuclear ribonucleoprotein-associated proteins B and B | 4 | 4 | 4 | 14.2 | 14.2 | 14.2 | 24.614 | 0 | 32.486 | 415740000 | 32 |
| Q12906 | Interleukin enhancer-binding factor 3 | 11 | 11 | 11 | 14.2 | 14.2 | 14.2 | 95.337 | 0 | 84.101 | 426360000 | 56 |
| P05023 | Sodium/potassium-transporting ATPase subunit alpha-1 | 13 | 13 | 8 | 15.5 | 15.5 | 10.2 | 112.89 | 0 | 96.673 | 484240000 | 51 |
| Q9NR30 | Nucleolar RNA helicase 2 | 12 | 12 | 12 | 20.9 | 20.9 | 20.9 | 87.343 | 0 | 88.743 | 417120000 | 24 |
| Q9UEY8 | Gamma-adducin | 7 | 7 | 7 | 16.1 | 16.1 | 16.1 | 79.154 | 0 | 69.615 | 431530000 | 22 |
| Q14764 | Major vault protein | 13 | 13 | 13 | 19.1 | 19.1 | 19.1 | 99.326 | 0 | 126.65 | 403950000 | 51 |
| P18077 | 60S ribosomal protein L35a | 7 | 7 | 7 | 54.5 | 54.5 | 54.5 | 12.538 | 0 | 43.41 | 412040000 | 52 |
| Q13573 | SNW domain-containing protein 1 | 10 | 10 | 10 | 20.9 | 20.9 | 20.9 | 61.494 | 0 | 74.943 | 382930000 | 45 |
| Q16891 | MICOS complex subunit MIC60 | 13 | 13 | 13 | 21.9 | 21.9 | 21.9 | 83.677 | 0 | 100.41 | 426180000 | 46 |
| P13796 | Plastin-2 | 11 | 11 | 11 | 20.6 | 20.6 | 20.6 | 70.288 | 0 | 144.12 | 441060000 | 38 |

| | | | | | | | | | | | | |
|---|---|---|---|---|---|---|---|---|---|---|---|---|
| Q2TAY7 | WD40 repeat-containing protein SMU1;WD40 repeat-containing protein SMU1, N-terminally processed | 8 | 8 | 8 | 17 | 17 | 17 | 57.543 | 0 | 68.157 | 361940000 | 17 |
| Q2TAY7 | WD40 repeat-containing protein SMU1;WD40 repeat-containing protein SMU1, N-terminally processed | 8 | 8 | 8 | 17 | 17 | 17 | 57.543 | 0 | 68.157 | 361940000 | 17 |
| Q15025 | TNFAIP3-interacting protein 1 | 11 | 11 | 11 | 18.6 | 18.6 | 18.6 | 71.863 | 0 | 103.02 | 390130000 | 38 |
| P08195 | 4F2 cell-surface antigen heavy chain | 10 | 10 | 10 | 19.8 | 19.8 | 19.8 | 67.993 | 0 | 78.39 | 403500000 | 23 |
| Q13247 | Serine/arginine-rich splicing factor 6 | 6 | 6 | 6 | 18.6 | 18.6 | 18.6 | 39.586 | 0 | 42.582 | 364840000 | 35 |
| Q15459 | Splicing factor 3A subunit 1 | 11 | 11 | 11 | 19 | 19 | 19 | 88.885 | 0 | 91.478 | 350260000 | 36 |
| P52701 | DNA mismatch repair protein Msh6 | 10 | 10 | 10 | 8.2 | 8.2 | 8.2 | 152.78 | 0 | 83.915 | 380160000 | 47 |
| P41250 | Glycine--tRNA ligase | 8 | 8 | 8 | 15.3 | 15.3 | 15.3 | 83.165 | 0 | 84.347 | 383480000 | 33 |
| O43175 | D-3-phosphoglycerate dehydrogenase | 9 | 9 | 9 | 25.1 | 25.1 | 25.1 | 56.65 | 0 | 82.735 | 353500000 | 32 |
| Q13283 | Ras GTPase-activating protein-binding protein 1 | 8 | 8 | 7 | 22.1 | 22.1 | 19.5 | 52.164 | 0 | 66.938 | 365270000 | 33 |
| Q7Z417 | Nuclear fragile X mental retardation-interacting protein 2 | 12 | 12 | 12 | 21.2 | 21.2 | 21.2 | 76.12 | 0 | 114.02 | 361560000 | 42 |
| Q702N8 | Xin actin-binding repeat-containing protein 1 | 14 | 14 | 14 | 10.5 | 10.5 | 10.5 | 198.56 | 0 | 107.04 | 350290000 | 45 |
| Q14444 | Caprin-1 | 8 | 8 | 8 | 14.2 | 14.2 | 14.2 | 78.365 | 0 | 76.374 | 375710000 | 44 |
| P24666 | Low molecular weight phosphotyrosine protein phosphatase | 6 | 6 | 6 | 34.8 | 34.8 | 34.8 | 18.042 | 0 | 69.874 | 325990000 | 29 |
| P62879 | Guanine nucleotide-binding protein G(I)/G(S)/G(T) subunit beta-2 | 9 | 3 | 3 | 35 | 13.8 | 13.8 | 37.331 | 0 | 27.793 | 343450000 | 21 |
| Q12874 | Splicing factor 3A subunit 3 | 9 | 9 | 9 | 24.4 | 24.4 | 24.4 | 58.848 | 0 | 100.71 | 315890000 | 39 |
| P21796 | Voltage-dependent anion-selective channel protein 1 | 7 | 7 | 7 | 30 | 30 | 30 | 30.772 | 0 | 45.743 | 343700000 | 20 |
| P06493 | Cyclin-dependent kinase 1 | 7 | 7 | 6 | 26.3 | 26.3 | 23.6 | 34.095 | 0 | 70.108 | 344020000 | 36 |
| Q96FV9 | THO complex subunit 1 | 8 | 8 | 8 | 15.1 | 15.1 | 15.1 | 75.665 | 0 | 56.18 | 342110000 | 23 |
| Q9BUJ2 | Heterogeneous nuclear ribonucleoprotein U-like protein 1 | 7 | 7 | 7 | 12.1 | 12.1 | 12.1 | 95.737 | 0 | 103.6 | 318610000 | 30 |
| Q13151 | Heterogeneous nuclear ribonucleoprotein A0 | 6 | 6 | 6 | 26.6 | 26.6 | 26.6 | 30.84 | 0 | 52.857 | 293200000 | 24 |
| Q9P258 | Protein RCC2 | 7 | 7 | 7 | 17 | 17 | 17 | 56.084 | 0 | 59.98 | 306920000 | 15 |
| Q92878 | DNA repair protein RAD50 | 14 | 14 | 14 | 11.7 | 11.7 | 11.7 | 153.89 | 0 | 104.96 | 326590000 | 39 |
| P27348 | 14-3-3 protein theta | 6 | 6 | 5 | 29.8 | 29.8 | 26.9 | 27.764 | 0 | 254.85 | 308570000 | 30 |
| Q9UKM9 | RNA-binding protein Raly | 6 | 6 | 6 | 26.5 | 26.5 | 26.5 | 32.463 | 0 | 40.935 | 287500000 | 25 |

| | | | | | | | | | | | | |
|---|---|---|---|---|---|---|---|---|---|---|---|---|
| P21580 | Tumor necrosis factor alpha-induced protein 3;A20p50;A20p37 | 10 | 10 | 10 | 14.8 | 14.8 | 14.8 | 89.613 | 0 | 79.08 | 296940000 | 40 |
| O75533 | Splicing factor 3B subunit 1 | 11 | 11 | 11 | 10.9 | 10.9 | 10.9 | 145.83 | 0 | 99.789 | 281980000 | 35 |
| Q9BY77 | Polymerase delta-interacting protein 3 | 8 | 8 | 8 | 29.2 | 29.2 | 29.2 | 46.089 | 0 | 85.224 | 276530000 | 24 |
| P45880 | Voltage-dependent anion-selective channel protein 2 | 5 | 5 | 5 | 24.8 | 24.8 | 24.8 | 31.566 | 0 | 39.035 | 282050000 | 25 |
| O00139 | Kinesin-like protein KIF2A | 9 | 9 | 7 | 12.3 | 12.3 | 9.6 | 79.954 | 0 | 97.377 | 254960000 | 32 |
| O00139 | Kinesin-like protein KIF2A | 9 | 9 | 7 | 12.3 | 12.3 | 9.6 | 79.954 | 0 | 97.377 | 254960000 | 32 |
| P17987 | T-complex protein 1 subunit alpha | 8 | 8 | 8 | 20 | 20 | 20 | 60.343 | 0 | 74.411 | 262720000 | 19 |
| Q13769 | THO complex subunit 5 homolog | 7 | 7 | 7 | 14.9 | 14.9 | 14.9 | 78.507 | 0 | 57.217 | 263810000 | 26 |
| Q8NC51 | Plasminogen activator inhibitor 1 RNA-binding protein | 8 | 8 | 8 | 26.2 | 26.2 | 26.2 | 44.965 | 0 | 130.05 | 285870000 | 34 |
| Q07955 | Serine/arginine-rich splicing factor 1 | 8 | 8 | 7 | 33.5 | 33.5 | 30.6 | 27.744 | 0 | 63.213 | 269280000 | 39 |
| P07814 | Bifunctional glutamate/proline--tRNA ligase;Glutamate--tRNA ligase;Proline--tRNA ligase | 10 | 10 | 10 | 7.9 | 7.9 | 7.9 | 170.59 | 0 | 58.312 | 254220000 | 19 |
| Q14974 | Importin subunit beta-1 | 7 | 7 | 7 | 9.7 | 9.7 | 9.7 | 97.169 | 0 | 70.834 | 248080000 | 38 |
| P12956 | X-ray repair cross-complementing protein 6 | 7 | 7 | 7 | 15.4 | 15.4 | 15.4 | 69.842 | 0 | 51.89 | 261570000 | 20 |
| Q99459 | Cell division cycle 5-like protein | 8 | 8 | 8 | 11.6 | 11.6 | 11.6 | 92.25 | 0 | 59.752 | 229620000 | 32 |
| Q13557 | Calcium/calmodulin-dependent protein kinase type II subunit delta | 7 | 7 | 7 | 19.8 | 19.8 | 19.8 | 56.369 | 0 | 62.995 | 244990000 | 26 |
| P09661 | U2 small nuclear ribonucleoprotein A | 5 | 5 | 5 | 23.1 | 23.1 | 23.1 | 28.415 | 0 | 57.703 | 217180000 | 28 |
| P04843 | Dolichyl-diphosphooligosaccharide--protein glycosyltransferase subunit 1 | 9 | 9 | 9 | 17.5 | 17.5 | 17.5 | 68.569 | 0 | 55.83 | 243460000 | 18 |
| Q13951 | Core-binding factor subunit beta | 7 | 7 | 7 | 38.5 | 38.5 | 38.5 | 21.508 | 0 | 52.499 | 236260000 | 24 |
| Q13045 | Protein flightless-1 homolog | 10 | 10 | 10 | 9.6 | 9.6 | 9.6 | 144.75 | 0 | 77.434 | 216330000 | 35 |
| Q9Y3X0 | Coiled-coil domain-containing protein 9 | 4 | 4 | 4 | 9.6 | 9.6 | 9.6 | 59.702 | 0 | 35.681 | 204720000 | 24 |
| P31942 | Heterogeneous nuclear ribonucleoprotein H3 | 4 | 4 | 4 | 15.3 | 15.3 | 15.3 | 36.926 | 0 | 45.813 | 219630000 | 23 |
| Q9UJV9 | Probable ATP-dependent RNA helicase DDX41 | 6 | 6 | 6 | 13 | 13 | 13 | 69.837 | 0 | 57.012 | 209350000 | 23 |
| P51116 | Fragile X mental retardation syndrome-related protein 2 | 8 | 6 | 6 | 14.1 | 11 | 11 | 74.222 | 0 | 58.597 | 224470000 | 34 |
| P49368 | T-complex protein 1 subunit gamma | 7 | 7 | 7 | 13.2 | 13.2 | 13.2 | 60.533 | 0 | 58 | 217790000 | 34 |
| P43490 | Nicotinamide phosphoribosyltransferase | 5 | 5 | 5 | 16.7 | 16.7 | 16.7 | 55.52 | 0 | 65.967 | 226680000 | 21 |
| P78371 | T-complex protein 1 subunit beta | 5 | 5 | 5 | 13.3 | 13.3 | 13.3 | 57.488 | 0 | 50.316 | 211650000 | 29 |
| P53814 | Smoothelin | 8 | 8 | 8 | 11 | 11 | 11 | 99.058 | 0 | 72.647 | 208460000 | 38 |

| | | | | | | | | | | | | |
|---|---|---|---|---|---|---|---|---|---|---|---|---|
| Q14739 | Lamin-B receptor | 7 | 7 | 7 | 16.4 | 16.4 | 16.4 | 70.702 | 0 | 50.164 | 229080000 | 27 |
| Q9Y3Z3 | Deoxynucleoside triphosphate triphosphohydrolase SAMHD1 | 7 | 7 | 7 | 15.5 | 15.5 | 15.5 | 72.2 | 0 | 71.443 | 206880000 | 36 |
| P50990 | T-complex protein 1 subunit theta | 7 | 7 | 7 | 14.1 | 14.1 | 14.1 | 59.62 | 0 | 52.862 | 198150000 | 26 |
| Q6I9Y2 | THO complex subunit 7 homolog | 6 | 6 | 6 | 35.3 | 35.3 | 35.3 | 23.743 | 0 | 46.321 | 224770000 | 23 |
| Q6I9Y2 | THO complex subunit 7 homolog | 6 | 6 | 6 | 35.3 | 35.3 | 35.3 | 23.743 | 0 | 46.321 | 224770000 | 23 |
| Q13242 | Serine/arginine-rich splicing factor 9 | 6 | 5 | 5 | 28.1 | 24.9 | 24.9 | 25.542 | 0 | 37.64 | 197660000 | 18 |
| Q9ULV4 | Coronin-1C | 8 | 8 | 8 | 15.2 | 15.2 | 15.2 | 53.248 | 0 | 69.506 | 201530000 | 42 |
| Q12905 | Interleukin enhancer-binding factor 2 | 5 | 5 | 5 | 18.5 | 18.5 | 18.5 | 43.062 | 0 | 38.634 | 200840000 | 25 |
| O95816 | BAG family molecular chaperone regulator 2 | 5 | 5 | 5 | 30.3 | 30.3 | 30.3 | 23.772 | 0 | 56.324 | 202370000 | 20 |
| P42704 | Leucine-rich PPR motif-containing protein, mitochondrial | 11 | 11 | 11 | 9.7 | 9.7 | 9.7 | 157.9 | 0 | 82.592 | 204930000 | 21 |
| P08865 | 40S ribosomal protein SA | 3 | 3 | 3 | 12.9 | 12.9 | 12.9 | 32.854 | 0 | 57.307 | 203580000 | 29 |
| O60884 | DnaJ homolog subfamily A member 2 | 4 | 4 | 4 | 11.7 | 11.7 | 11.7 | 45.745 | 0 | 41.261 | 193160000 | 34 |
| O43684 | Mitotic checkpoint protein BUB3 | 5 | 5 | 5 | 15.9 | 15.9 | 15.9 | 37.154 | 0 | 70.426 | 175630000 | 33 |
| O43390 | Heterogeneous nuclear ribonucleoprotein R | 9 | 4 | 4 | 13.9 | 7.4 | 7.4 | 70.942 | 0 | 41.497 | 196340000 | 8 |
| P50402 | Emerin | 5 | 5 | 5 | 24.4 | 24.4 | 24.4 | 28.994 | 0 | 72.251 | 185860000 | 42 |
| Q92945 | Far upstream element-binding protein 2 | 6 | 6 | 6 | 9.8 | 9.8 | 9.8 | 73.114 | 0 | 61.399 | 174370000 | 29 |
| Q9Y230 | RuvB-like 2 | 7 | 7 | 7 | 17.5 | 17.5 | 17.5 | 51.156 | 0 | 47.893 | 189350000 | 12 |
| P51149 | Ras-related protein Rab-7a | 4 | 4 | 4 | 22.2 | 22.2 | 22.2 | 23.489 | 0 | 37.452 | 186040000 | 16 |
| P62304 | Small nuclear ribonucleoprotein E | 2 | 2 | 2 | 29.3 | 29.3 | 29.3 | 10.803 | 0 | 27.207 | 250890000 | 22 |
| Q02978 | Mitochondrial 2-oxoglutarate/malate carrier protein | 4 | 4 | 4 | 22.6 | 22.6 | 22.6 | 34.061 | 0 | 71.404 | 182850000 | 21 |
| P36873 | Serine/threonine-protein phosphatase PP1-gamma catalytic subunit | 19 | 3 | 3 | 55.7 | 11.5 | 11.5 | 36.983 | 0 | 33.05 | 164290000 | 11 |
| P05161 | Ubiquitin-like protein ISG15 | 3 | 3 | 3 | 18.8 | 18.8 | 18.8 | 17.887 | 0 | 18.169 | 170250000 | 16 |
| P36542 | ATP synthase subunit gamma, mitochondrial | 3 | 3 | 3 | 11.1 | 11.1 | 11.1 | 32.996 | 0 | 27.573 | 170490000 | 13 |
| O75369 | Filamin-B | 13 | 6 | 6 | 5.3 | 2.8 | 2.8 | 278.16 | 0 | 58.234 | 166500000 | 18 |
| P49458 | Signal recognition particle 9 kDa protein | 4 | 4 | 4 | 44.2 | 44.2 | 44.2 | 10.112 | 0 | 41.46 | 177500000 | 32 |
| P17980 | 26S protease regulatory subunit 6A | 4 | 4 | 4 | 13.2 | 13.2 | 13.2 | 49.203 | 0 | 29.043 | 173880000 | 15 |
| P55769 | NHP2-like protein 1;NHP2-like protein 1, N-terminally processed | 3 | 3 | 3 | 31.2 | 31.2 | 31.2 | 14.173 | 0 | 38.703 | 170860000 | 22 |
| Q15007 | Pre-mRNA-splicing regulator WTAP | 6 | 6 | 6 | 20.5 | 20.5 | 20.5 | 44.243 | 0 | 67.83 | 169520000 | 34 |
| Q9UBM7 | 7-dehydrocholesterol reductase | 4 | 4 | 4 | 8.6 | 8.6 | 8.6 | 54.489 | 0 | 39.133 | 169120000 | 17 |

| | | | | | | | | | | | | |
|---|---|---|---|---|---|---|---|---|---|---|---|---|
| P16403 | Histone H1.2 | 9 | 2 | 2 | 33.8 | 9.9 | 9.9 | 21.364 | 0 | 14.48 | 160890000 | 24 |
| P40227 | T-complex protein 1 subunit zeta | 5 | 5 | 5 | 11.7 | 11.7 | 11.7 | 58.024 | 0 | 36.813 | 156640000 | 12 |
| P78527 | DNA-dependent protein kinase catalytic subunit | 12 | 12 | 12 | 2.9 | 2.9 | 2.9 | 469.08 | 0 | 95.614 | 164630000 | 29 |
| Q96J01 | THO complex subunit 3 | 6 | 6 | 6 | 18.8 | 18.8 | 18.8 | 38.771 | 0 | 38.935 | 154030000 | 13 |
| P02786 | Transferrin receptor protein 1;Transferrin receptor protein 1, serum form | 6 | 6 | 6 | 10.3 | 10.3 | 10.3 | 84.87 | 0 | 59.635 | 163510000 | 21 |
| P20700 | Lamin-B1 | 9 | 8 | 8 | 17.1 | 15.7 | 15.7 | 66.408 | 0 | 68.818 | 159250000 | 30 |
| Q13761 | Runt-related transcription factor 3 | 4 | 4 | 4 | 10.4 | 10.4 | 10.4 | 44.355 | 0 | 27.103 | 178800000 | 19 |
| P50991 | T-complex protein 1 subunit delta | 4 | 4 | 4 | 9.6 | 9.6 | 9.6 | 57.924 | 0 | 36.985 | 152590000 | 16 |
| Q69YQ0 | Cytospin-A | 9 | 9 | 9 | 9.1 | 9.1 | 9.1 | 124.6 | 0 | 58.871 | 155280000 | 31 |
| Q15637 | Splicing factor 1 | 4 | 4 | 4 | 8 | 8 | 8 | 68.329 | 0 | 34.911 | 147530000 | 11 |
| Q6UN15 | Pre-mRNA 3-end-processing factor FIP1 | 2 | 2 | 2 | 6.1 | 6.1 | 6.1 | 66.526 | 0 | 31.96 | 147210000 | 13 |
| Q8IYB3 | Serine/arginine repetitive matrix protein 1 | 3 | 3 | 3 | 5.6 | 5.6 | 5.6 | 102.33 | 0 | 32.966 | 149650000 | 27 |
| Q9Y3F4 | Serine-threonine kinase receptor-associated protein | 5 | 5 | 5 | 19.4 | 19.4 | 19.4 | 38.438 | 0 | 40.805 | 144840000 | 11 |
| P62310 | U6 snRNA-associated Sm-like protein LSm3 | 3 | 3 | 3 | 21.6 | 21.6 | 21.6 | 11.845 | 0 | 19.372 | 134280000 | 8 |
| Q96SB3 | Neurabin-2 | 9 | 9 | 9 | 11.7 | 11.7 | 11.7 | 89.191 | 0 | 53.266 | 141590000 | 24 |
| P61026 | Ras-related protein Rab-10 | 3 | 3 | 3 | 17 | 17 | 17 | 22.541 | 0 | 29.5 | 143580000 | 20 |
| Q92888 | Rho guanine nucleotide exchange factor 1 | 6 | 6 | 6 | 9.2 | 9.2 | 9.2 | 102.43 | 0 | 40.581 | 140680000 | 19 |
| P48444 | Coatomer subunit delta | 5 | 5 | 5 | 12.5 | 12.5 | 12.5 | 57.21 | 0 | 45.341 | 136660000 | 17 |
| Q9UPN4 | Centrosomal protein of 131 kDa | 12 | 12 | 12 | 13.2 | 13.2 | 13.2 | 122.15 | 0 | 76.471 | 118200000 | 29 |
| Q9BWJ5 | Splicing factor 3B subunit 5 | 3 | 3 | 3 | 59.3 | 59.3 | 59.3 | 10.135 | 0 | 30.721 | 138340000 | 16 |
| Q9H3U1 | Protein unc-45 homolog A | 5 | 5 | 5 | 7.1 | 7.1 | 7.1 | 103.08 | 0 | 42.593 | 133120000 | 12 |
| P61158 | Actin-related protein 3 | 5 | 5 | 5 | 14.6 | 14.6 | 14.6 | 47.371 | 0 | 92.645 | 133540000 | 14 |
| Q9Y613 | FH1/FH2 domain-containing protein 1 | 5 | 5 | 5 | 5.7 | 5.7 | 5.7 | 126.55 | 0 | 43.777 | 127920000 | 19 |
| P53007 | Tricarboxylate transport protein, mitochondrial | 4 | 4 | 4 | 13.8 | 13.8 | 13.8 | 34.012 | 0 | 24.876 | 133210000 | 8 |
| O75569 | Interferon-inducible double-stranded RNA-dependent protein kinase activator A | 4 | 4 | 4 | 16.3 | 16.3 | 16.3 | 34.404 | 0 | 28.7 | 119140000 | 13 |
| Q9NX63 | MICOS complex subunit MIC19 | 5 | 5 | 5 | 23.3 | 23.3 | 23.3 | 26.152 | 0 | 41.986 | 139810000 | 15 |
| P14625 | Endoplasmin | 7 | 6 | 6 | 8.2 | 6.5 | 6.5 | 92.468 | 0 | 34.788 | 114570000 | 12 |
| P56134 | ATP synthase subunit f, mitochondrial | 2 | 2 | 2 | 25.5 | 25.5 | 25.5 | 10.918 | 0 | 16.99 | 193510000 | 21 |
| Q53EZ4 | Centrosomal protein of 55 kDa | 4 | 4 | 4 | 14.9 | 14.9 | 14.9 | 54.178 | 0 | 79.825 | 116900000 | 17 |
| Q9NW64 | Pre-mRNA-splicing factor RBM22 | 5 | 5 | 5 | 13.6 | 13.6 | 13.6 | 46.895 | 0 | 34.868 | 117750000 | 18 |

| | | | | | | | | | | | | |
|---|---|---|---|---|---|---|---|---|---|---|---|---|
| P22087 | rRNA 2-O-methyltransferase fibrillarin | 5 | 5 | 5 | 19.3 | 19.3 | 19.3 | 33.784 | 0 | 38.598 | 128450000 | 22 |
| Q16531 | DNA damage-binding protein 1 | 6 | 6 | 6 | 7.5 | 7.5 | 7.5 | 126.97 | 0 | 45.594 | 120680000 | 9 |
| P61160 | Actin-related protein 2 | 5 | 5 | 5 | 16.5 | 16.5 | 16.5 | 44.76 | 0 | 35.772 | 120400000 | 13 |
| P61160 | Actin-related protein 2 | 5 | 5 | 5 | 16.5 | 16.5 | 16.5 | 44.76 | 0 | 35.772 | 120400000 | 13 |
| Q5SW79 | Centrosomal protein of 170 kDa | 6 | 6 | 6 | 4.2 | 4.2 | 4.2 | 175.29 | 0 | 36.424 | 126340000 | 17 |
| Q9P2I0 | Cleavage and polyadenylation specificity factor subunit 2 | 6 | 6 | 6 | 9.3 | 9.3 | 9.3 | 88.486 | 0 | 78.321 | 119810000 | 14 |
| P39656 | Dolichyl-diphosphooligosaccharide--protein glycosyltransferase 48 kDa subunit | 5 | 5 | 5 | 11 | 11 | 11 | 50.8 | 0 | 32.181 | 135810000 | 11 |
| O94925 | Glutaminase kidney isoform, mitochondrial | 4 | 4 | 4 | 8.8 | 8.8 | 8.8 | 73.46 | 0 | 39.805 | 117860000 | 15 |
| Q9Y265 | RuvB-like 1 | 5 | 5 | 5 | 12.5 | 12.5 | 12.5 | 50.227 | 0 | 33.397 | 118850000 | 16 |
| Q3MHD2 | Protein LSM12 homolog | 3 | 3 | 3 | 15.9 | 15.9 | 15.9 | 21.701 | 0 | 20.532 | 116030000 | 22 |
| P62333 | 26S protease regulatory subunit 10B | 5 | 5 | 5 | 18 | 18 | 18 | 44.172 | 0 | 70.109 | 115040000 | 5 |
| P62306 | Small nuclear ribonucleoprotein F | 3 | 3 | 3 | 33.7 | 33.7 | 33.7 | 9.7251 | 0 | 35.333 | 123650000 | 20 |
| Q8NAV1 | Pre-mRNA-splicing factor 38A | 6 | 6 | 6 | 25 | 25 | 25 | 37.476 | 0 | 43.544 | 102220000 | 10 |
| P22695 | Cytochrome b-c1 complex subunit 2, mitochondrial | 6 | 6 | 6 | 19.6 | 19.6 | 19.6 | 48.442 | 0 | 52.802 | 106110000 | 6 |
| P48643 | T-complex protein 1 subunit epsilon | 4 | 4 | 4 | 10.2 | 10.2 | 10.2 | 59.67 | 0 | 29.857 | 109440000 | 7 |
| P30504 | HLA class I histocompatibility antigen, Cw-4 alpha chain | 12 | 3 | 2 | 32 | 7.1 | 4.1 | 40.994 | 0 | 23.007 | 108810000 | 17 |
| Q9Y2D5 | A-kinase anchor protein 2 | 3 | 3 | 3 | 5 | 5 | 5 | 94.659 | 0 | 21.132 | 105310000 | 11 |
| Q13561 | Dynactin subunit 2 | 3 | 3 | 3 | 9.5 | 9.5 | 9.5 | 44.23 | 0 | 20.956 | 103240000 | 13 |
| P35249 | Replication factor C subunit 4 | 4 | 4 | 4 | 13.8 | 13.8 | 13.8 | 39.681 | 0 | 22.44 | 109470000 | 3 |
| Q7RTV0 | PHD finger-like domain-containing protein 5A | 4 | 4 | 4 | 37.3 | 37.3 | 37.3 | 12.405 | 0 | 28.213 | 112980000 | 18 |
| P53618 | Coatomer subunit beta | 6 | 6 | 6 | 8.1 | 8.1 | 8.1 | 107.14 | 0 | 36.069 | 108510000 | 7 |
| Q6PKG0 | La-related protein 1 | 6 | 6 | 6 | 7.1 | 7.1 | 7.1 | 123.51 | 0 | 46.598 | 105610000 | 29 |
| Q06787 | Fragile X mental retardation protein 1 | 6 | 5 | 5 | 9.2 | 7.9 | 7.9 | 71.174 | 0 | 48.142 | 111670000 | 21 |
| Q13123 | Protein Red | 2 | 2 | 2 | 4.3 | 4.3 | 4.3 | 65.601 | 0 | 21.746 | 96912000 | 5 |
| P51571 | Translocon-associated protein subunit delta | 3 | 3 | 3 | 17.9 | 17.9 | 17.9 | 18.998 | 0 | 23.325 | 103050000 | 10 |
| Q9UN86 | Ras GTPase-activating protein-binding protein 2 | 3 | 2 | 2 | 7.3 | 4.8 | 4.8 | 54.12 | 0 | 29.522 | 102660000 | 14 |
| P14868 | Aspartate--tRNA ligase, cytoplasmic | 5 | 5 | 5 | 10 | 10 | 10 | 57.136 | 0 | 31.104 | 99648000 | 23 |
| Q9H223 | EH domain-containing protein 4 | 3 | 3 | 3 | 7.6 | 7.6 | 7.6 | 61.174 | 0 | 34.226 | 99894000 | 9 |
| Q14257 | Reticulocalbin-2 | 3 | 3 | 3 | 10.1 | 10.1 | 10.1 | 36.876 | 0 | 19.187 | 93094000 | 4 |

| | | | | | | | | | | | | |
|---|---|---|---|---|---|---|---|---|---|---|---|---|
| Q99661 | Kinesin-like protein KIF2C | 6 | 4 | 4 | 8.6 | 5.9 | 5.9 | 81.312 | 0 | 32.06 | 89367000 | 12 |
| Q9NZ01 | Very-long-chain enoyl-CoA reductase | 5 | 5 | 5 | 14.6 | 14.6 | 14.6 | 36.034 | 0 | 31.417 | 93833000 | 25 |
| Q9P0L0 | Vesicle-associated membrane protein-associated protein A | 5 | 5 | 5 | 17.7 | 17.7 | 17.7 | 27.893 | 0 | 29.014 | 92644000 | 15 |
| Q10570 | Cleavage and polyadenylation specificity factor subunit 1 | 5 | 5 | 5 | 5.1 | 5.1 | 5.1 | 160.88 | 0 | 43.329 | 94507000 | 20 |
| Q13148 | TAR DNA-binding protein 43 | 2 | 2 | 2 | 5.1 | 5.1 | 5.1 | 44.739 | 0 | 25.381 | 88781000 | 13 |
| P20592 | Interferon-induced GTP-binding protein Mx2 | 3 | 3 | 3 | 4.6 | 4.6 | 4.6 | 82.088 | 0 | 25.751 | 85788000 | 10 |
| O14579 | Coatomer subunit epsilon | 2 | 2 | 2 | 19.8 | 19.8 | 19.8 | 34.482 | 0 | 14.563 | 87924000 | 4 |
| Q15428 | Splicing factor 3A subunit 2 | 4 | 4 | 4 | 12.1 | 12.1 | 12.1 | 49.255 | 0 | 24.324 | 87008000 | 6 |
| Q01196 | Runt-related transcription factor 1 | 4 | 4 | 4 | 10.8 | 10.8 | 10.8 | 48.736 | 0 | 26.127 | 89851000 | 18 |
| Q99623 | Prohibitin-2 | 4 | 4 | 4 | 18.4 | 18.4 | 18.4 | 33.296 | 0 | 31.932 | 80305000 | 10 |
| Q7Z6E9 | E3 ubiquitin-protein ligase RBBP6 | 4 | 4 | 4 | 2.7 | 2.7 | 2.7 | 201.56 | 0 | 23.383 | 80851000 | 12 |
| Q02040 | A-kinase anchor protein 17A | 5 | 5 | 5 | 6.8 | 6.8 | 6.8 | 80.735 | 0 | 29.71 | 74321000 | 18 |
| P02794 | Ferritin heavy chain;Ferritin heavy chain, N-terminally processed | 3 | 3 | 3 | 16.9 | 16.9 | 16.9 | 21.225 | 0 | 18.376 | 85411000 | 8 |
| Q96DI7 | U5 small nuclear ribonucleoprotein 40 kDa protein | 2 | 2 | 2 | 13.4 | 13.4 | 13.4 | 39.31 | 0 | 38.742 | 82059000 | 6 |
| O15143 | Actin-related protein 2/3 complex subunit 1B | 2 | 2 | 2 | 8.6 | 8.6 | 8.6 | 40.949 | 0 | 20.352 | 81500000 | 9 |
| P29692 | Elongation factor 1-delta | 4 | 4 | 3 | 19.2 | 19.2 | 16 | 31.121 | 0 | 58.424 | 72171000 | 11 |
| Q9UPN7 | Serine/threonine-protein phosphatase 6 regulatory subunit 1 | 4 | 4 | 4 | 5.9 | 5.9 | 5.9 | 96.723 | 0 | 42.284 | 80663000 | 15 |
| P09382 | Galectin-1 | 3 | 3 | 3 | 26.7 | 26.7 | 26.7 | 14.716 | 0 | 19.036 | 70978000 | 4 |
| Q96FS4 | Signal-induced proliferation-associated protein 1 | 5 | 5 | 5 | 5.4 | 5.4 | 5.4 | 112.15 | 0 | 41.301 | 76013000 | 11 |
| Q9Y608 | Leucine-rich repeat flightless-interacting protein 2 | 4 | 4 | 4 | 5.3 | 5.3 | 5.3 | 82.17 | 0 | 39.833 | 69295000 | 12 |
| O15042 | U2 snRNP-associated SURP motif-containing protein | 2 | 2 | 2 | 2.5 | 2.5 | 2.5 | 118.29 | 0 | 12.828 | 70293000 | 4 |
| Q96C19 | EF-hand domain-containing protein D2 | 3 | 3 | 3 | 15 | 15 | 15 | 26.697 | 0 | 21.099 | 66097000 | 7 |
| P28066 | Proteasome subunit alpha type-5 | 3 | 3 | 3 | 16.2 | 16.2 | 16.2 | 26.411 | 0 | 19.348 | 68935000 | 5 |
| Q13077 | TNF receptor-associated factor 1 | 2 | 2 | 2 | 5.3 | 5.3 | 5.3 | 46.163 | 0 | 22.247 | 71993000 | 8 |
| P08574 | Cytochrome c1, heme protein, mitochondrial | 2 | 2 | 2 | 7.1 | 7.1 | 7.1 | 35.422 | 0 | 11.538 | 72755000 | 4 |
| Q9NTJ3 | Structural maintenance of chromosomes protein 4 | 6 | 6 | 6 | 5.1 | 5.1 | 5.1 | 147.18 | 0 | 46.637 | 70060000 | 13 |

| | | | | | | | | | | | | |
|---|---|---|---|---|---|---|---|---|---|---|---|---|
| O00165 | HCLS1-associated protein X-1 | 4 | 4 | 4 | 13.6 | 13.6 | 13.6 | 31.62 | 0 | 26.35 | 71720000 | 10 |
| Q9NZR1 | Tropomodulin-2 | 5 | 4 | 4 | 13.1 | 13.1 | 13.1 | 39.595 | 0 | 42.561 | 67811000 | 20 |
| O00743 | Serine/threonine-protein phosphatase 6 catalytic subunit;Serine/threonine-protein phosphatase 6 catalytic subunit, N-terminally processed | 2 | 2 | 2 | 7.2 | 7.2 | 7.2 | 35.144 | 0 | 20.751 | 68748000 | 11 |
| P52732 | Kinesin-like protein KIF11 | 4 | 4 | 4 | 3.7 | 3.7 | 3.7 | 119.16 | 0 | 23.905 | 71149000 | 4 |
| P24534 | Elongation factor 1-beta | 3 | 2 | 2 | 13.8 | 9.8 | 9.8 | 24.763 | 0 | 16.314 | 63293000 | 3 |
| Q9H7N4 | Splicing factor, arginine/serine-rich 19 | 4 | 4 | 4 | 2.9 | 2.9 | 2.9 | 139.27 | 0 | 24.436 | 61782000 | 7 |
| Q13595 | Transformer-2 protein homolog alpha | 5 | 4 | 4 | 18.8 | 15.2 | 15.2 | 32.688 | 0 | 25.46 | 55915000 | 10 |
| P78406 | mRNA export factor | 2 | 2 | 2 | 9.8 | 9.8 | 9.8 | 40.968 | 0 | 14.315 | 60725000 | 3 |
| Q00653 | Nuclear factor NF-kappa-B p100 subunit;Nuclear factor NF-kappa-B p52 subunit | 3 | 3 | 3 | 4.4 | 4.4 | 4.4 | 96.748 | 0 | 17.229 | 65005000 | 4 |
| Q1KMD3 | Heterogeneous nuclear ribonucleoprotein U-like protein 2 | 3 | 3 | 3 | 5.6 | 5.6 | 5.6 | 85.104 | 0 | 27.059 | 60864000 | 7 |
| P41240 | Tyrosine-protein kinase CSK | 2 | 2 | 2 | 4.9 | 4.9 | 4.9 | 50.704 | 0 | 11.839 | 61092000 | 6 |
| P48047 | ATP synthase subunit O, mitochondrial | 2 | 2 | 2 | 11.7 | 11.7 | 11.7 | 23.277 | 0 | 18.091 | 59828000 | 9 |
| P61221 | ATP-binding cassette sub-family E member 1 | 3 | 3 | 3 | 5.5 | 5.5 | 5.5 | 67.314 | 0 | 22.039 | 57450000 | 13 |
| Q9Y5M8 | Signal recognition particle receptor subunit beta | 2 | 2 | 2 | 12.5 | 12.5 | 12.5 | 29.702 | 0 | 17.037 | 58148000 | 5 |
| Q99832 | T-complex protein 1 subunit eta | 3 | 3 | 3 | 7.9 | 7.9 | 7.9 | 59.366 | 0 | 23.139 | 56418000 | 5 |
| P31939 | Bifunctional purine biosynthesis protein PURH;Phosphoribosylaminoimidazolecarboxamide formyltransferase;IMP cyclohydrolase | 2 | 2 | 2 | 6.2 | 6.2 | 6.2 | 64.615 | 0 | 16.33 | 56664000 | 7 |
| Q53GQ0 | Very-long-chain 3-oxoacyl-CoA reductase | 2 | 2 | 2 | 9.9 | 9.9 | 9.9 | 34.324 | 0 | 12.762 | 56161000 | 7 |
| O00214 | Galectin-8 | 2 | 2 | 2 | 6 | 6 | 6 | 35.808 | 0 | 11.995 | 48456000 | 12 |
| P40926 | Malate dehydrogenase, mitochondrial | 3 | 3 | 3 | 9.8 | 9.8 | 9.8 | 35.503 | 0 | 19.233 | 50812000 | 8 |
| P08579 | U2 small nuclear ribonucleoprotein B | 2 | 2 | 2 | 12 | 12 | 12 | 25.486 | 0 | 23.632 | 56642000 | 11 |
| Q02818 | Nucleobindin-1 | 2 | 2 | 2 | 2.8 | 2.8 | 2.8 | 53.879 | 0 | 11.458 | 49786000 | 6 |
| Q13523 | Serine/threonine-protein kinase PRP4 homolog | 2 | 2 | 2 | 2.7 | 2.7 | 2.7 | 116.99 | 0 | 14.801 | 49546000 | 5 |
| O95864 | Fatty acid desaturase 2 | 2 | 2 | 2 | 5.9 | 5.9 | 5.9 | 52.259 | 0 | 12.576 | 46484000 | 6 |
| O60306 | Intron-binding protein aquarius | 2 | 2 | 2 | 1.8 | 1.8 | 1.8 | 171.29 | 0 | 13.52 | 45085000 | 6 |
| P11387 | DNA topoisomerase 1 | 4 | 4 | 4 | 5.4 | 5.4 | 5.4 | 90.725 | 0 | 24.732 | 48065000 | 11 |

| | | | | | | | | | | | | |
|---|---|---|---|---|---|---|---|---|---|---|---|---|
| Q96T37 | Putative RNA-binding protein 15 | 2 | 2 | 2 | 2.5 | 2.5 | 2.5 | 107.19 | 0 | 22.321 | 47267000 | 9 |
| P08754 | Guanine nucleotide-binding protein G(k) subunit alpha | 4 | 2 | 2 | 13.8 | 7.1 | 7.1 | 40.532 | 0 | 15.626 | 48179000 | 8 |
| Q14699 | Raftlin | 2 | 2 | 2 | 5.2 | 5.2 | 5.2 | 63.145 | 0 | 12.293 | 45907000 | 5 |
| Q8TBC3 | SH3KBP1-binding protein 1 | 2 | 2 | 2 | 3.4 | 3.4 | 3.4 | 76.343 | 0 | 13.958 | 45258000 | 2 |
| P24539 | ATP synthase F(0) complex subunit B1, mitochondrial | 2 | 2 | 2 | 9.8 | 9.8 | 9.8 | 28.908 | 0 | 14.061 | 47049000 | 9 |
| Q9Y305 | Acyl-coenzyme A thioesterase 9, mitochondrial | 3 | 3 | 3 | 7.5 | 7.5 | 7.5 | 49.901 | 0 | 19.573 | 39821000 | 4 |
| Q8IWX8 | Calcium homeostasis endoplasmic reticulum protein | 2 | 2 | 2 | 2.9 | 2.9 | 2.9 | 103.7 | 0 | 15.327 | 39540000 | 8 |
| P46087 | Probable 28S rRNA (cytosine(4447)-C(5))-methyltransferase | 5 | 5 | 5 | 8.5 | 8.5 | 8.5 | 89.301 | 0 | 29.883 | 54687000 | 7 |
| P34897 | Serine hydroxymethyltransferase, mitochondrial | 3 | 3 | 3 | 6.9 | 6.9 | 6.9 | 55.992 | 0 | 19.949 | 42648000 | 6 |
| Q15427 | Splicing factor 3B subunit 4 | 2 | 2 | 2 | 8.7 | 8.7 | 8.7 | 44.385 | 0 | 19.124 | 39511000 | 10 |
| Q15125 | 3-beta-hydroxysteroid-Delta(8),Delta(7)-isomerase | 2 | 2 | 2 | 12.2 | 12.2 | 12.2 | 26.352 | 0 | 15.828 | 38378000 | 7 |
| Q63HN8 | E3 ubiquitin-protein ligase RNF213 | 4 | 4 | 4 | 0.7 | 0.7 | 0.7 | 591.4 | 0 | 24.762 | 36676000 | 7 |
| P63151;Q66LE6;Q00005 | Serine/threonine-protein phosphatase 2A 55 kDa regulatory subunit B alpha isoform;Serine/threonine-protein phosphatase 2A 55 kDa regulatory subunit B delta isoform;Serine/threonine-protein phosphatase 2A 55 kDa regulatory subunit B beta isoform | 2 | 2 | 2 | 5.6 | 5.6 | 5.6 | 51.691 | 0 | 23.661 | 38204000 | 4 |
| P37268 | Squalene synthase | 2 | 2 | 2 | 5.5 | 5.5 | 5.5 | 48.115 | 0 | 16.322 | 33264000 | 2 |
| Q8N684 | Cleavage and polyadenylation specificity factor subunit 7 | 2 | 2 | 2 | 4.2 | 4.2 | 4.2 | 52.049 | 0 | 11.576 | 25960000 | 5 |
| P00338 | L-lactate dehydrogenase A chain | 4 | 4 | 3 | 13.6 | 13.6 | 9.9 | 36.688 | 0 | 28.377 | 43127000 | 5 |
| Q13155 | Aminoacyl tRNA synthase complex-interacting multifunctional protein 2 | 2 | 2 | 2 | 7.8 | 7.8 | 7.8 | 35.348 | 0 | 11.35 | 24446000 | 3 |
| **P17844** | Probable ATP-dependent RNA helicase DDX5 | 22 | 14 | 14 | 38.3 | 23.9 | 23.9 | 69.147 | 0 | 137.92 | 1370100000 | 84 |
| **P63104** | 14-3-3 protein zeta/delta | 5 | 4 | 4 | 24.5 | 21.6 | 21.6 | 27.745 | 0 | 76.976 | 205910000 | 36 |
| **P30153** | Serine/threonine-protein phosphatase 2A 65 kDa regulatory subunit A alpha isoform | 4 | 4 | 4 | 7.3 | 7.3 | 7.3 | 65.308 | 0 | 26.505 | 45664000 | 10 |
| **P63244** | Guanine nucleotide-binding protein subunit beta-2-like 1;Guanine nucleotide-binding protein subunit beta-2-like 1, N-terminally processed | 12 | 12 | 12 | 42.3 | 42.3 | 42.3 | 35.076 | 0 | 90.603 | 736670000 | 58 |
