## Supplementary Table 2 for "Protein Interactome of the Cancerous Inhibitor of Protein Phosphatase 2A (CIP2A) in Th17 Cells"

**Supplementry Table 2: Gene Ontology terms associated with CIP2A interactome**

The table lists Gene Ontology terms associated with the CIP2A interactome by the DAVID Bioinformatics application (Huang et al., 2009). For biological process, the FAT terms were used and provide an overview of the association of the proteins in **Figure 5**. These terms are listed together with an indication of their occurrence cluster by cluster in terms of number of proteins and proportion of associated nodes.

| GO.FAT.Term | Count | % of Nodes in cluster included | Cluster | Annotation for cluster |
|---|---|---|---|---|
| GO:0016070~RNA metabolic process | 68 | 0.96 | 1 | RNA metabolic process |
| GO:0010467~gene expression | 67 | 0.94 | 1 | RNA metabolic process |
| GO:0016071~mRNA metabolic process | 65 | 0.92 | 1 | RNA metabolic process |
| GO:0006396~RNA processing | 65 | 0.92 | 1 | RNA metabolic process |
| GO:0006397~mRNA processing | 63 | 0.89 | 1 | RNA metabolic process |
| GO:0008380~RNA splicing | 62 | 0.87 | 1 | RNA metabolic process |
| GO:0000375~RNA splicing, via transesterification reactions | 55 | 0.77 | 1 | RNA metabolic process |
| GO:0000398~mRNA splicing, via spliceosome | 55 | 0.77 | 1 | RNA metabolic process |
| GO:0000377~RNA splicing, via transesterification reactions with bulged adenosine as nucleophile | 55 | 0.77 | 1 | RNA metabolic process |
| GO:0034645~cellular macromolecule biosynthetic process | 39 | 0.55 | 1 | RNA metabolic process |
| GO:0034654~nucleobase-containing compound biosynthetic process | 33 | 0.46 | 1 | RNA metabolic process |
| GO:0019438~aromatic compound biosynthetic process | 33 | 0.46 | 1 | RNA metabolic process |
| GO:0018130~heterocycle biosynthetic process | 33 | 0.46 | 1 | RNA metabolic process |
| GO:0051641~cellular localization | 25 | 0.35 | 1 | RNA metabolic process |
| GO:0051169~nuclear transport | 24 | 0.34 | 1 | RNA metabolic process |
| GO:0006913~nucleocytoplasmic transport | 24 | 0.34 | 1 | RNA metabolic process |
| GO:0051649~establishment of localization in cell | 24 | 0.34 | 1 | RNA metabolic process |
| GO:0046907~intracellular transport | 24 | 0.34 | 1 | RNA metabolic process |
| GO:0006369~termination of RNA polymerase II transcription | 22 | 0.31 | 1 | RNA metabolic process |
| GO:0006403~RNA localization | 20 | 0.28 | 1 | RNA metabolic process |
| GO:0051641~cellular localization | 17 | 0.49 | 2 | Cellular localisation |
| GO:0051649~establishment of localization in cell | 16 | 0.46 | 2 | Cellular localisation |
| GO:0010467~gene expression | 13 | 0.37 | 2 | Cellular localisation |

| | | | | |
|---|---|---|---|---|
| GO:0046907~intracellular transport | 13 | 0.37 | 2 | Cellular localisation |
| GO:0007010~cytoskeleton organization | 12 | 0.34 | 2 | Cellular localisation |
| GO:0034645~cellular macromolecule biosynthetic process | 11 | 0.31 | 2 | Cellular localisation |
| GO:0022607~cellular component assembly | 11 | 0.31 | 2 | Cellular localisation |
| GO:0016070~RNA metabolic process | 10 | 0.29 | 2 | Cellular localisation |
| GO:0034654~nucleobase-containing compound biosynthetic process | 9 | 0.26 | 2 | Cellular localisation |
| GO:0019438~aromatic compound biosynthetic process | 9 | 0.26 | 2 | Cellular localisation |
| GO:0018130~heterocycle biosynthetic process | 9 | 0.26 | 2 | Cellular localisation |
| GO:0060341~regulation of cellular localization | 8 | 0.23 | 2 | Cellular localisation |
| GO:1903829~positive regulation of cellular protein localization | 6 | 0.17 | 2 | Cellular localisation |
| GO:1903827~regulation of cellular protein localization | 6 | 0.17 | 2 | Cellular localisation |
| GO:0030029~actin filament-based process | 5 | 0.14 | 2 | Cellular localisation |
| GO:0065003~macromolecular complex assembly | 5 | 0.14 | 2 | Cellular localisation |
| GO:0045935~positive regulation of nucleobase-containing compound metabolic process | 5 | 0.14 | 2 | Cellular localisation |
| GO:0051173~positive regulation of nitrogen compound metabolic process | 5 | 0.14 | 2 | Cellular localisation |
| GO:0016032~viral process | 4 | 0.11 | 2 | Cellular localisation |
| GO:0044419~interspecies interaction between organisms | 4 | 0.11 | 2 | Cellular localisation |
| GO:0034654~nucleobase-containing compound biosynthetic process | 16 | 0.57 | 3 | Biosynthetic process |
| GO:0019438~aromatic compound biosynthetic process | 16 | 0.57 | 3 | Biosynthetic process |
| GO:0018130~heterocycle biosynthetic process | 16 | 0.57 | 3 | Biosynthetic process |
| GO:0034645~cellular macromolecule biosynthetic process | 13 | 0.46 | 3 | Biosynthetic process |
| GO:0022607~cellular component assembly | 11 | 0.39 | 3 | Biosynthetic process |
| GO:0051641~cellular localization | 10 | 0.36 | 3 | Biosynthetic process |
| GO:0065003~macromolecular complex assembly | 10 | 0.36 | 3 | Biosynthetic process |
| GO:0006403~RNA localization | 9 | 0.32 | 3 | Biosynthetic process |
| GO:1904874~positive regulation of telomerase RNA localization to Cajal body | 9 | 0.32 | 3 | Biosynthetic process |
| GO:1904872~regulation of telomerase RNA localization to Cajal body | 9 | 0.32 | 3 | Biosynthetic process |
| GO:0090672~telomerase RNA localization | 9 | 0.32 | 3 | Biosynthetic process |
| GO:0090671~telomerase RNA localization to Cajal body | 9 | 0.32 | 3 | Biosynthetic process |
| GO:0090670~RNA localization to Cajal body | 9 | 0.32 | 3 | Biosynthetic process |
| GO:0045935~positive regulation of nucleobase-containing compound metabolic process | 9 | 0.32 | 3 | Biosynthetic process |
| GO:0051173~positive regulation of nitrogen compound metabolic process | 9 | 0.32 | 3 | Biosynthetic process |
| GO:1904869~regulation of protein localization to Cajal body | 8 | 0.29 | 3 | Biosynthetic process |

| | | | | |
|---|---|---|---|---|
| GO:1904871~positive regulation of protein localization to Cajal body | 8 | 0.29 | 3 | Biosynthetic process |
| GO:1904867~protein localization to Cajal body | 8 | 0.29 | 3 | Biosynthetic process |
| GO:1903405~protein localization to nuclear body | 8 | 0.29 | 3 | Biosynthetic process |
| GO:1990173~protein localization to nucleoplasm | 8 | 0.29 | 3 | Biosynthetic process |
| GO:0030029~actin filament-based process | 13 | 0.62 | 4 | cytoskeleton organization |
| GO:0007010~cytoskeleton organization | 13 | 0.62 | 4 | cytoskeleton organization |
| GO:0030036~actin cytoskeleton organization | 12 | 0.57 | 4 | cytoskeleton organization |
| GO:0022607~cellular component assembly | 11 | 0.52 | 4 | cytoskeleton organization |
| GO:0007015~actin filament organization | 10 | 0.48 | 4 | cytoskeleton organization |
| GO:0007155~cell adhesion | 10 | 0.48 | 4 | cytoskeleton organization |
| GO:0022610~biological adhesion | 10 | 0.48 | 4 | cytoskeleton organization |
| GO:0065003~macromolecular complex assembly | 10 | 0.48 | 4 | cytoskeleton organization |
| GO:0043254~regulation of protein complex assembly | 10 | 0.48 | 4 | cytoskeleton organization |
| GO:0098609~cell-cell adhesion | 9 | 0.43 | 4 | cytoskeleton organization |
| GO:0008154~actin polymerization or depolymerization | 9 | 0.43 | 4 | cytoskeleton organization |
| GO:0008064~regulation of actin polymerization or depolymerization | 9 | 0.43 | 4 | cytoskeleton organization |
| GO:0030832~regulation of actin filament length | 9 | 0.43 | 4 | cytoskeleton organization |
| GO:0030833~regulation of actin filament polymerization | 9 | 0.43 | 4 | cytoskeleton organization |
| GO:0051641~cellular localization | 8 | 0.38 | 4 | cytoskeleton organization |
| GO:0030835~negative regulation of actin filament depolymerization | 5 | 0.24 | 4 | cytoskeleton organization |
| GO:0030042~actin filament depolymerization | 5 | 0.24 | 4 | cytoskeleton organization |
| GO:0051693~actin filament capping | 5 | 0.24 | 4 | cytoskeleton organization |
| GO:0051649~establishment of localization in cell | 5 | 0.24 | 4 | cytoskeleton organization |
| GO:0030834~regulation of actin filament depolymerization | 5 | 0.24 | 4 | cytoskeleton organization |
| GO:0034645~cellular macromolecule biosynthetic process | 16 | 0.80 | 5 | biosynthetic process |
| GO:0010467~gene expression | 13 | 0.65 | 5 | biosynthetic process |
| GO:0051641~cellular localization | 12 | 0.60 | 5 | biosynthetic process |
| GO:0016070~RNA metabolic process | 10 | 0.50 | 5 | biosynthetic process |
| GO:0016032~viral process | 10 | 0.50 | 5 | biosynthetic process |
| GO:0044419~interspecies interaction between organisms | 10 | 0.50 | 5 | biosynthetic process |
| GO:0044403~symbiosis, encompassing mutualism through parasitism | 10 | 0.50 | 5 | biosynthetic process |
| GO:0044764~multi-organism cellular process | 10 | 0.50 | 5 | biosynthetic process |
| GO:0051649~establishment of localization in cell | 10 | 0.50 | 5 | biosynthetic process |

| | | | | |
|---|---|---|---|---|
| GO:0046907~intracellular transport | 10 | 0.50 | 5 | biosynthetic process |
| GO:0016071~mRNA metabolic process | 9 | 0.45 | 5 | biosynthetic process |
| GO:0006396~RNA processing | 9 | 0.45 | 5 | biosynthetic process |
| GO:0019058~viral life cycle | 9 | 0.45 | 5 | biosynthetic process |
| GO:0034654~nucleobase-containing compound biosynthetic process | 9 | 0.45 | 5 | biosynthetic process |
| GO:0019438~aromatic compound biosynthetic process | 9 | 0.45 | 5 | biosynthetic process |
| GO:0018130~heterocycle biosynthetic process | 9 | 0.45 | 5 | biosynthetic process |
| GO:0007155~cell adhesion | 7 | 0.35 | 5 | biosynthetic process |
| GO:0022610~biological adhesion | 7 | 0.35 | 5 | biosynthetic process |
| GO:0098609~cell-cell adhesion | 7 | 0.35 | 5 | biosynthetic process |
| GO:0022607~cellular component assembly | 5 | 0.25 | 5 | biosynthetic process |
| GO:0034654~nucleobase-containing compound biosynthetic process | 10 | 0.67 | 6 | biosynthetic process |
| GO:0034645~cellular macromolecule biosynthetic process | 10 | 0.67 | 6 | biosynthetic process |
| GO:0019438~aromatic compound biosynthetic process | 10 | 0.67 | 6 | biosynthetic process |
| GO:0018130~heterocycle biosynthetic process | 10 | 0.67 | 6 | biosynthetic process |
| GO:0045935~positive regulation of nucleobase-containing compound metabolic process | 8 | 0.53 | 6 | biosynthetic process |
| GO:0051173~positive regulation of nitrogen compound metabolic process | 8 | 0.53 | 6 | biosynthetic process |
| GO:0060249~anatomical structure homeostasis | 7 | 0.47 | 6 | biosynthetic process |
| GO:0000723~telomere maintenance | 7 | 0.47 | 6 | biosynthetic process |
| GO:0032200~telomere organization | 7 | 0.47 | 6 | biosynthetic process |
| GO:0022607~cellular component assembly | 7 | 0.47 | 6 | biosynthetic process |
| GO:0065003~macromolecular complex assembly | 7 | 0.47 | 6 | biosynthetic process |
| GO:0016070~RNA metabolic process | 5 | 0.33 | 6 | biosynthetic process |
| GO:0016032~viral process | 5 | 0.33 | 6 | biosynthetic process |
| GO:0044419~interspecies interaction between organisms | 5 | 0.33 | 6 | biosynthetic process |
| GO:0044403~symbiosis, encompassing mutualism through parasitism | 5 | 0.33 | 6 | biosynthetic process |
| GO:0044764~multi-organism cellular process | 5 | 0.33 | 6 | biosynthetic process |
| GO:0010467~gene expression | 5 | 0.33 | 6 | biosynthetic process |
| GO:0043254~regulation of protein complex assembly | 4 | 0.27 | 6 | biosynthetic process |
| GO:0051641~cellular localization | 2 | 0.13 | 6 | biosynthetic process |
| GO:0007155~cell adhesion | 2 | 0.13 | 6 | biosynthetic process |
| GO:0051641~cellular localization | 5 | 0.42 | 7 | cellular localization |
| GO:0051649~establishment of localization in cell | 5 | 0.42 | 7 | cellular localization |

| | | | | |
|---|---|---|---|---|
| GO:0046907~intracellular transport | 5 | 0.42 | 7 | cellular localization |
| GO:0034654~nucleobase-containing compound biosynthetic process | 5 | 0.42 | 7 | cellular localization |
| GO:0019438~aromatic compound biosynthetic process | 5 | 0.42 | 7 | cellular localization |
| GO:0018130~heterocycle biosynthetic process | 5 | 0.42 | 7 | cellular localization |
| GO:0010467~gene expression | 2 | 0.17 | 7 | cellular localization |
| GO:0016032~viral process | 1 | 0.08 | 7 | cellular localization |
| GO:0044419~interspecies interaction between organisms | 1 | 0.08 | 7 | cellular localization |
| GO:0044403~symbiosis, encompassing mutualism through parasitism | 1 | 0.08 | 7 | cellular localization |
| GO:0044764~multi-organism cellular process | 1 | 0.08 | 7 | cellular localization |
| GO:0007155~cell adhesion | 1 | 0.08 | 7 | cellular localization |
| GO:0022610~biological adhesion | 1 | 0.08 | 7 | cellular localization |
| GO:0098609~cell-cell adhesion | 1 | 0.08 | 7 | cellular localization |
| GO:0034645~cellular macromolecule biosynthetic process | 1 | 0.08 | 7 | cellular localization |
| GO:0022607~cellular component assembly | 1 | 0.08 | 7 | cellular localization |
| GO:0032272~negative regulation of protein polymerization | 1 | 0.08 | 7 | cellular localization |
| GO:0065003~macromolecular complex assembly | 1 | 0.08 | 7 | cellular localization |
| GO:0043254~regulation of protein complex assembly | 1 | 0.08 | 7 | cellular localization |
| GO:0008380~RNA splicing | 0 | 0.00 | 7 | cellular localization |
| GO:0010467~gene expression | 9 | 0.75 | 8 | gene expression |
| GO:0016070~RNA metabolic process | 8 | 0.67 | 8 | gene expression |
| GO:0034645~cellular macromolecule biosynthetic process | 7 | 0.58 | 8 | gene expression |
| GO:0008380~RNA splicing | 6 | 0.50 | 8 | gene expression |
| GO:0000375~RNA splicing, via transesterification reactions | 6 | 0.50 | 8 | gene expression |
| GO:0000398~mRNA splicing, via spliceosome | 6 | 0.50 | 8 | gene expression |
| GO:0000377~RNA splicing, via transesterification reactions with bulged adenosine as nucleophile | 6 | 0.50 | 8 | gene expression |
| GO:0006397~mRNA processing | 6 | 0.50 | 8 | gene expression |
| GO:0016071~mRNA metabolic process | 6 | 0.50 | 8 | gene expression |
| GO:0006396~RNA processing | 6 | 0.50 | 8 | gene expression |
| GO:0034654~nucleobase-containing compound biosynthetic process | 6 | 0.50 | 8 | gene expression |
| GO:0019438~aromatic compound biosynthetic process | 6 | 0.50 | 8 | gene expression |
| GO:0018130~heterocycle biosynthetic process | 6 | 0.50 | 8 | gene expression |
| GO:0048024~regulation of mRNA splicing, via spliceosome | 4 | 0.33 | 8 | gene expression |
| GO:0043484~regulation of RNA splicing | 4 | 0.33 | 8 | gene expression |

| | | | | |
|---|---|---|---|---|
| GO:1903311~regulation of mRNA metabolic process | 4 | 0.33 | 8 | gene expression |
| GO:0050684~regulation of mRNA processing | 4 | 0.33 | 8 | gene expression |
| GO:0048025~negative regulation of mRNA splicing, via spliceosome | 3 | 0.25 | 8 | gene expression |
| GO:0033119~negative regulation of RNA splicing | 3 | 0.25 | 8 | gene expression |
| GO:1903312~negative regulation of mRNA metabolic process | 3 | 0.25 | 8 | gene expression |
| GO:0007155~cell adhesion | 7 | 0.70 | 9 | Cell adhesion |
| GO:0022610~biological adhesion | 7 | 0.70 | 9 | Cell adhesion |
| GO:0051641~cellular localization | 6 | 0.60 | 9 | Cell adhesion |
| GO:0098609~cell-cell adhesion | 5 | 0.50 | 9 | Cell adhesion |
| GO:0022607~cellular component assembly | 5 | 0.50 | 9 | Cell adhesion |
| GO:0007010~cytoskeleton organization | 5 | 0.50 | 9 | Cell adhesion |
| GO:0030029~actin filament-based process | 4 | 0.40 | 9 | Cell adhesion |
| GO:0030036~actin cytoskeleton organization | 4 | 0.40 | 9 | Cell adhesion |
| GO:0007015~actin filament organization | 4 | 0.40 | 9 | Cell adhesion |
| GO:0051649~establishment of localization in cell | 4 | 0.40 | 9 | Cell adhesion |
| GO:0060341~regulation of cellular localization | 4 | 0.40 | 9 | Cell adhesion |
| GO:0030835~negative regulation of actin filament depolymerization | 2 | 0.20 | 9 | Cell adhesion |
| GO:0030042~actin filament depolymerization | 2 | 0.20 | 9 | Cell adhesion |
| GO:0051693~actin filament capping | 2 | 0.20 | 9 | Cell adhesion |
| GO:0030834~regulation of actin filament depolymerization | 2 | 0.20 | 9 | Cell adhesion |
| GO:1901880~negative regulation of protein depolymerization | 2 | 0.20 | 9 | Cell adhesion |
| GO:0043242~negative regulation of protein complex disassembly | 2 | 0.20 | 9 | Cell adhesion |
| GO:0051261~protein depolymerization | 2 | 0.20 | 9 | Cell adhesion |
| GO:0032272~negative regulation of protein polymerization | 2 | 0.20 | 9 | Cell adhesion |
| GO:0065003~macromolecular complex assembly | 2 | 0.20 | 9 | Cell adhesion |
| GO:0016032~viral process | 5 | 0.83 | 10 | viral process |
| GO:0044419~interspecies interaction between organisms | 5 | 0.83 | 10 | viral process |
| GO:0044403~symbiosis, encompassing mutualism through parasitism | 5 | 0.83 | 10 | viral process |
| GO:0044764~multi-organism cellular process | 5 | 0.83 | 10 | viral process |
| GO:0007155~cell adhesion | 3 | 0.50 | 10 | viral process |
| GO:0022610~biological adhesion | 3 | 0.50 | 10 | viral process |
| GO:0098609~cell-cell adhesion | 3 | 0.50 | 10 | viral process |
| GO:0034654~nucleobase-containing compound biosynthetic process | 3 | 0.50 | 10 | viral process |

| | | | | |
|---|---|---|---|---|
| GO:0034645~cellular macromolecule biosynthetic process | 3 | 0.50 | 10 | viral process |
| GO:0019438~aromatic compound biosynthetic process | 3 | 0.50 | 10 | viral process |
| GO:0018130~heterocycle biosynthetic process | 3 | 0.50 | 10 | viral process |
| GO:0016070~RNA metabolic process | 2 | 0.33 | 10 | viral process |
| GO:0010467~gene expression | 2 | 0.33 | 10 | viral process |
| GO:0045935~positive regulation of nucleobase-containing compound metabolic process | 2 | 0.33 | 10 | viral process |
| GO:0051173~positive regulation of nitrogen compound metabolic process | 2 | 0.33 | 10 | viral process |
| GO:0006403~RNA localization | 1 | 0.17 | 10 | viral process |
| GO:0050657~nucleic acid transport | 1 | 0.17 | 10 | viral process |
| GO:0050658~RNA transport | 1 | 0.17 | 10 | viral process |
| GO:0051236~establishment of RNA localization | 1 | 0.17 | 10 | viral process |
| GO:0051028~mRNA transport | 1 | 0.17 | 10 | viral process |
| GO:0010467~gene expression | 5 | 1.00 | 11 | gene expression |
| GO:0016070~RNA metabolic process | 4 | 0.80 | 11 | gene expression |
| GO:0006396~RNA processing | 3 | 0.60 | 11 | gene expression |
| GO:0016032~viral process | 3 | 0.60 | 11 | gene expression |
| GO:0044419~interspecies interaction between organisms | 3 | 0.60 | 11 | gene expression |
| GO:0044403~symbiosis, encompassing mutualism through parasitism | 3 | 0.60 | 11 | gene expression |
| GO:0044764~multi-organism cellular process | 3 | 0.60 | 11 | gene expression |
| GO:0034645~cellular macromolecule biosynthetic process | 3 | 0.60 | 11 | gene expression |
| GO:0006403~RNA localization | 1 | 0.20 | 11 | gene expression |
| GO:0031124~mRNA 3-end processing | 1 | 0.20 | 11 | gene expression |
| GO:0071426~ribonucleoprotein complex export from nucleus | 1 | 0.20 | 11 | gene expression |
| GO:0071166~ribonucleoprotein complex localization | 1 | 0.20 | 11 | gene expression |
| GO:0051168~nuclear export | 1 | 0.20 | 11 | gene expression |
| GO:0051641~cellular localization | 1 | 0.20 | 11 | gene expression |
| GO:0051169~nuclear transport | 1 | 0.20 | 11 | gene expression |
| GO:0006913~nucleocytoplasmic transport | 1 | 0.20 | 11 | gene expression |
| GO:0007155~cell adhesion | 1 | 0.20 | 11 | gene expression |
| GO:0022610~biological adhesion | 1 | 0.20 | 11 | gene expression |
| GO:0051649~establishment of localization in cell | 1 | 0.20 | 11 | gene expression |
| GO:0019058~viral life cycle | 1 | 0.20 | 11 | gene expression |
| GO:0010467~gene expression | 3 | 0.75 | 12 | gene expression |

| | | | | |
|---|---|---|---|---|
| GO:0034645~cellular macromolecule biosynthetic process | 3 | 0.75 | 12 | gene expression |
| GO:0008380~RNA splicing | 1 | 0.25 | 12 | gene expression |
| GO:0000375~RNA splicing, via transesterification reactions | 1 | 0.25 | 12 | gene expression |
| GO:0000398~mRNA splicing, via spliceosome | 1 | 0.25 | 12 | gene expression |
| GO:0000377~RNA splicing, via transesterification reactions with bulged adenosine as nucleophile | 1 | 0.25 | 12 | gene expression |
| GO:0006397~mRNA processing | 1 | 0.25 | 12 | gene expression |
| GO:0016071~mRNA metabolic process | 1 | 0.25 | 12 | gene expression |
| GO:0006396~RNA processing | 1 | 0.25 | 12 | gene expression |
| GO:0006403~RNA localization | 1 | 0.25 | 12 | gene expression |
| GO:0048024~regulation of mRNA splicing, via spliceosome | 1 | 0.25 | 12 | gene expression |
| GO:0050657~nucleic acid transport | 1 | 0.25 | 12 | gene expression |
| GO:0050658~RNA transport | 1 | 0.25 | 12 | gene expression |
| GO:0051236~establishment of RNA localization | 1 | 0.25 | 12 | gene expression |
| GO:0051028~mRNA transport | 1 | 0.25 | 12 | gene expression |
| GO:0015931~nucleobase-containing compound transport | 1 | 0.25 | 12 | gene expression |
| GO:0043484~regulation of RNA splicing | 1 | 0.25 | 12 | gene expression |
| GO:1903311~regulation of mRNA metabolic process | 1 | 0.25 | 12 | gene expression |
| GO:0050684~regulation of mRNA processing | 1 | 0.25 | 12 | gene expression |
| GO:0000381~regulation of alternative mRNA splicing, via spliceosome | 1 | 0.25 | 12 | gene expression |
| GO:0030029~actin filament-based process | 2 | 0.50 | 13 | actin filament-based process |
| GO:0007155~cell adhesion | 2 | 0.50 | 13 | actin filament-based process |
| GO:0022610~biological adhesion | 2 | 0.50 | 13 | actin filament-based process |
| GO:0030048~actin filament-based movement | 2 | 0.50 | 13 | actin filament-based process |
| GO:0051641~cellular localization | 1 | 0.25 | 13 | actin filament-based process |
| GO:0051649~establishment of localization in cell | 1 | 0.25 | 13 | actin filament-based process |
| GO:0046907~intracellular transport | 1 | 0.25 | 13 | actin filament-based process |
| GO:0060341~regulation of cellular localization | 1 | 0.25 | 13 | actin filament-based process |
| GO:0008380~RNA splicing | 0 | 0.00 | 13 | actin filament-based process |
| GO:0000375~RNA splicing, via transesterification reactions | 0 | 0.00 | 13 | actin filament-based process |
| GO:0000398~mRNA splicing, via spliceosome | 0 | 0.00 | 13 | actin filament-based process |
| GO:0000377~RNA splicing, via transesterification reactions with bulged adenosine as nucleophile | 0 | 0.00 | 13 | actin filament-based process |
| GO:0006397~mRNA processing | 0 | 0.00 | 13 | actin filament-based process |
| GO:0016071~mRNA metabolic process | 0 | 0.00 | 13 | actin filament-based process |

| | | | | |
|---|---|---|---|---|
| GO:0006396~RNA processing | 0 | 0.00 | 13 | actin filament-based process |
| GO:0006403~RNA localization | 0 | 0.00 | 13 | actin filament-based process |
| GO:0006369~termination of RNA polymerase II transcription | 0 | 0.00 | 13 | actin filament-based process |
| GO:0048024~regulation of mRNA splicing, via spliceosome | 0 | 0.00 | 13 | actin filament-based process |
| GO:0050657~nucleic acid transport | 0 | 0.00 | 13 | actin filament-based process |
| GO:0050658~RNA transport | 0 | 0.00 | 13 | actin filament-based process |
| GO:0051641~cellular localization | 2 | 0.50 | 14 | cellular localisation |
| GO:0051169~nuclear transport | 2 | 0.50 | 14 | cellular localisation |
| GO:0051649~establishment of localization in cell | 2 | 0.50 | 14 | cellular localisation |
| GO:0046907~intracellular transport | 2 | 0.50 | 14 | cellular localisation |
| GO:0007084~mitotic nuclear envelope reassembly | 2 | 0.50 | 14 | cellular localisation |
| GO:0051168~nuclear export | 1 | 0.25 | 14 | cellular localisation |
| GO:0016032~viral process | 1 | 0.25 | 14 | cellular localisation |
| GO:0044419~interspecies interaction between organisms | 1 | 0.25 | 14 | cellular localisation |
| GO:0044403~symbiosis, encompassing mutualism through parasitism | 1 | 0.25 | 14 | cellular localisation |
| GO:0044764~multi-organism cellular process | 1 | 0.25 | 14 | cellular localisation |
| GO:0006913~nucleocytoplasmic transport | 1 | 0.25 | 14 | cellular localisation |
| GO:0007155~cell adhesion | 1 | 0.25 | 14 | cellular localisation |
| GO:0022610~biological adhesion | 1 | 0.25 | 14 | cellular localisation |
| GO:0060341~regulation of cellular localization | 1 | 0.25 | 14 | cellular localisation |
| GO:0098609~cell-cell adhesion | 1 | 0.25 | 14 | cellular localisation |
| GO:1903829~positive regulation of cellular protein localization | 1 | 0.25 | 14 | cellular localisation |
| GO:1903827~regulation of cellular protein localization | 1 | 0.25 | 14 | cellular localisation |
| GO:0008380~RNA splicing | 0 | 0.00 | 14 | cellular localisation |
| GO:0000375~RNA splicing, via transesterification reactions | 0 | 0.00 | 14 | cellular localisation |
| GO:0000398~mRNA splicing, via spliceosome | 0 | 0.00 | 14 | cellular localisation |
| GO:0008380~RNA splicing | 3 | 0.75 | 15 | RNA splicing |
| GO:0006396~RNA processing | 3 | 0.75 | 15 | RNA splicing |
| GO:0016070~RNA metabolic process | 3 | 0.75 | 15 | RNA splicing |
| GO:0010467~gene expression | 3 | 0.75 | 15 | RNA splicing |
| GO:0016032~viral process | 2 | 0.50 | 15 | RNA splicing |
| GO:0044419~interspecies interaction between organisms | 2 | 0.50 | 15 | RNA splicing |
| GO:0044403~symbiosis, encompassing mutualism through parasitism | 2 | 0.50 | 15 | RNA splicing |

| | | | | |
|---|---|---|---|---|
| GO:0044764~multi-organism cellular process | 2 | 0.50 | 15 | RNA splicing |
| GO:0034654~nucleobase-containing compound biosynthetic process | 2 | 0.50 | 15 | RNA splicing |
| GO:0034645~cellular macromolecule biosynthetic process | 2 | 0.50 | 15 | RNA splicing |
| GO:0019438~aromatic compound biosynthetic process | 2 | 0.50 | 15 | RNA splicing |
| GO:0018130~heterocycle biosynthetic process | 2 | 0.50 | 15 | RNA splicing |
| GO:0000375~RNA splicing, via transesterification reactions | 1 | 0.25 | 15 | RNA splicing |
| GO:0000398~mRNA splicing, via spliceosome | 1 | 0.25 | 15 | RNA splicing |
| GO:0000377~RNA splicing, via transesterification reactions with bulged adenosine as nucleophile | 1 | 0.25 | 15 | RNA splicing |
| GO:0006397~mRNA processing | 1 | 0.25 | 15 | RNA splicing |
| GO:0016071~mRNA metabolic process | 1 | 0.25 | 15 | RNA splicing |
| GO:0006403~RNA localization | 1 | 0.25 | 15 | RNA splicing |
| GO:0050657~nucleic acid transport | 1 | 0.25 | 15 | RNA splicing |
| GO:0050658~RNA transport | 1 | 0.25 | 15 | RNA splicing |
