## Supplementary Table 3 for "Protein Interactome of the Cancerous Inhibitor of Protein Phosphatase 2A (CIP2A) in Th17 Cells"

**Supplementry Table 3:** GO SLIM Biological Process

Enriched biological process for the proteins associated with CIP2A interactome. The biological processes enrichment was calculated relative to Th17 cells proteins background in earlier report from a proteomic characterisation (Tripathi et al., 2019)

| PANTHER GO-Slim Biological Process | Fold Enrichment | Raw P-value | FDR |
|---|---|---|---|
| sensory perception of sound (GO:0007605) | 10.26 | 1.22E-04 | 4.02E-03 |
| muscle contraction (GO:0006936) | 5.88 | 3.25E-05 | 1.25E-03 |
| RNA splicing, via transesterification reactions (GO:0000375) | 5.21 | 3.94E-13 | 3.03E-11 |
| mRNA splicing, via spliceosome (GO:0000398) | 4.98 | 1.03E-13 | 1.19E-11 |
| mesoderm development (GO:0007498) | 4.7 | 7.21E-04 | 1.67E-02 |
| mRNA processing (GO:0006397) | 4.43 | 2.57E-15 | 5.94E-13 |
| sensory perception (GO:0007600) | 3.96 | 1.88E-03 | 2.89E-02 |
| mitochondrion organization (GO:0007005) | 3.37 | 5.41E-04 | 1.39E-02 |
| protein folding (GO:0006457) | 3.23 | 1.23E-03 | 2.37E-02 |
| cellular component morphogenesis (GO:0032989) | 3.19 | 1.68E-05 | 9.72E-04 |
| cellular component movement (GO:0006928) | 2.45 | 3.30E-03 | 4.23E-02 |
| system process (GO:0003008) | 2.27 | 1.96E-03 | 2.83E-02 |
| multicellular organismal process (GO:0032501) | 2.09 | 1.29E-03 | 2.29E-02 |
| single-multicellular organism process (GO:0044707) | 2.09 | 1.29E-03 | 2.13E-02 |
| RNA metabolic process (GO:0016070) | 1.66 | 3.01E-05 | 1.39E-03 |
| cellular component organization (GO:0016043) | 1.57 | 1.70E-04 | 4.92E-03 |
| cellular component organization or biogenesis (GO:0071840) | 1.46 | 8.59E-04 | 1.80E-02 |
| nucleobase-containing compound metabolic process (GO:0006139) | 1.33 | 2.46E-03 | 3.34E-02 |
| Unclassified (UNCLASSIFIED) | 0.8 | 3.78E-03 | 4.59E-02 |
