## Supplementary Table 4 for "Protein Interactome of the Cancerous Inhibitor of Protein Phosphatase 2A (CIP2A) in Th17 Cells"

**Supplementary Table 4:** SRM MS CIP2A interactome validations

The table lists the peptides, associated transitions, collision energies and retention time windows used for the SRM analysis of the CIP2A vs. control IgG pull downs for proteins validated for the CIP2A interactome. The analysis was performed using an Easy-nLC with a 20 x 0.1 mm i.d. pre-column and a 150 mm x 75 µm i.d. analytical column, both packed with 5 µm Reprosil C18-bonded silica (Dr Maisch GmbH). A gradient of 8% to 43% B in 27 min, then to 100% B in 3 min, was used at a flow rate of 300 nl/min .

| Protein Name | Uniprot | Amino Acid Sequence | Ion | Natural Peptide Q1 (m/z) | Natural Peptide Q3 (m/z) | SIS Peptide Q1 (M/z) | SIS Peptide Q3 (m/z) | Collision Energy (V) | Retention Time window start (min) | Retention Time Window end (min) |
|---|---|---|---|---|---|---|---|---|---|---|
| AGK_HUMAN | Q53H12 | ATVFLNPAAC[+57.0215]K | y8 | 596.3132 | 920.4658 | 600.3203 | 928.48 | 20.8 | 16.06 | 18.06 |
| AGK_HUMAN | Q53H12 | ATVFLNPAAC[+57.0215]K | y7 | 596.3132 | 773.3974 | 600.3203 | 781.4116 | 20.8 | 16.06 | 18.06 |
| AGK_HUMAN | Q53H12 | ATVFLNPAAC[+57.0215]K | y6 | 596.3132 | 660.3134 | 600.3203 | 668.3276 | 20.8 | 16.06 | 18.06 |
| AGK_HUMAN | Q53H12 | ATVFLNPAAC[+57.0215]K | y5 | 596.3132 | 546.2704 | 600.3203 | 554.2846 | 20.8 | 16.06 | 18.06 |
| AGK_HUMAN | Q53H12 | ATVFLNPAAC[+57.0215]K | y2 | 596.3132 | 307.1435 | 600.3203 | 315.1577 | 20.8 | 16.06 | 18.06 |
| AGK_HUMAN | Q53H12 | VQHITDATLAIVK | y5 | 470.2768 | 543.3865 | 472.9482 | 551.4007 | 20.2 | 16.48 | 18.48 |
| AGK_HUMAN | Q53H12 | VQHITDATLAIVK | y4 | 470.2768 | 430.3024 | 472.9482 | 438.3166 | 20.2 | 16.48 | 18.48 |
| AGK_HUMAN | Q53H12 | VQHITDATLAIVK | y2 | 470.2768 | 246.1812 | 472.9482 | 254.1954 | 20.2 | 16.48 | 18.48 |
| AGK_HUMAN | Q53H12 | VQHITDATLAIVK | b3 | 470.2768 | 365.1932 | 472.9482 | 365.1932 | 20.2 | 16.48 | 18.48 |
| AGK_HUMAN | Q53H12 | YWYLGPLK | y6 | 520.2842 | 690.4185 | 524.2913 | 698.4327 | 18.5 | 22.4 | 24.4 |
| AGK_HUMAN | Q53H12 | YWYLGPLK | y5 | 520.2842 | 527.3552 | 524.2913 | 535.3694 | 18.5 | 22.4 | 24.4 |
| AGK_HUMAN | Q53H12 | YWYLGPLK | y4 | 520.2842 | 414.2711 | 524.2913 | 422.2853 | 18.5 | 22.4 | 24.4 |
| AGK_HUMAN | Q53H12 | YWYLGPLK | y3 | 520.2842 | 357.2496 | 524.2913 | 365.2638 | 18.5 | 22.4 | 24.4 |
| AGK_HUMAN | Q53H12 | YWYLGPLK | y6 | 520.2842 | 345.7129 | 524.2913 | 349.72 | 18.5 | 22.4 | 24.4 |
| CSK_HUMAN | Q53H12 | LLYPPETGLFLVR | y10 | 759.44 | 1128.641 | 764.4441 | 1138.649 | 25.7 | 26.11 | 28.11 |
| CSK_HUMAN | P41240 | LLYPPETGLFLVR | y9 | 759.44 | 1031.588 | 764.4441 | 1041.597 | 25.7 | 26.11 | 28.11 |
| CSK_HUMAN | P41240 | LLYPPETGLFLVR | y10 | 759.44 | 564.8242 | 764.4441 | 569.8284 | 25.7 | 26.11 | 28.11 |
| CSK_HUMAN | P41240 | LLYPPETGLFLVR | y3 | 759.44 | 194.1394 | 764.4441 | 199.1435 | 25.7 | 26.11 | 28.11 |
| CSK_HUMAN | P41240 | LLYPPETGLFLVR | b2 | 759.44 | 227.1754 | 764.4441 | 227.1754 | 25.7 | 26.11 | 28.11 |
| CSK_HUMAN | P41240 | LLQTIGK | y6 | 386.75 | 659.4087 | 390.7571 | 667.4229 | 14.5 | 11.34 | 13.34 |
| CSK_HUMAN | P41240 | LLQTIGK | y5 | 386.75 | 546.3246 | 390.7571 | 554.3388 | 14.5 | 11.34 | 13.34 |

| | | | | | | | | | | |
|---|---|---|---|---|---|---|---|---|---|---|
| CSK_HUMAN | P41240 | LLQTIGK | y4 | 386.75 | 418.266 | 390.7571 | 426.2802 | 14.5 | 11.34 | 13.34 |
| CSK_HUMAN | P41240 | LLQTIGK | y3 | 386.75 | 317.2183 | 390.7571 | 325.2325 | 14.5 | 11.34 | 13.34 |
| CSK_HUMAN | P41240 | LLQTIGK | y2 | 386.75 | 204.1343 | 390.7571 | 212.1485 | 14.5 | 11.34 | 13.34 |
| CSK_HUMAN | P41240 | WTAPEALR | y7 | 472.2534 | 757.4203 | 477.2576 | 767.4285 | 17.1 | 15.56 | 17.56 |
| CSK_HUMAN | P41240 | WTAPEALR | y6 | 472.2534 | 656.3726 | 477.2576 | 666.3809 | 17.1 | 15.56 | 17.56 |
| CSK_HUMAN | P41240 | WTAPEALR | y5 | 472.2534 | 585.3355 | 477.2576 | 595.3438 | 17.1 | 15.56 | 17.56 |
| CSK_HUMAN | P41240 | WTAPEALR | y3 | 472.2534 | 359.2401 | 477.2576 | 369.2484 | 17.1 | 15.56 | 17.56 |
| CSK_HUMAN | P41240 | WTAPEALR | y2 | 472.2534 | 288.203 | 477.2576 | 298.2113 | 17.1 | 15.56 | 17.56 |
| IF16_HUMAN | P41240 | EVDATSPAPSTSSTVK | y11 | 788.8887 | 1061.547 | 792.8958 | 1069.562 | 26.6 | 9.06 | 11.06 |
| IF16_HUMAN | Q16666 | EVDATSPAPSTSSTVK | y10 | 788.8887 | 974.5153 | 792.8958 | 982.5295 | 26.6 | 9.06 | 11.06 |
| IF16_HUMAN | Q16666 | EVDATSPAPSTSSTVK | y8 | 788.8887 | 806.4254 | 792.8958 | 814.4396 | 26.6 | 9.06 | 11.06 |
| IF16_HUMAN | Q16666 | EVDATSPAPSTSSTVK | b3 | 788.8887 | 344.1452 | 792.8958 | 344.1452 | 26.6 | 9.06 | 11.06 |
| IF16_HUMAN | Q16666 | EVDATSPAPSTSSTVK | b4 | 788.8887 | 415.1823 | 792.8958 | 415.1823 | 26.6 | 9.06 | 11.06 |
| IF16_HUMAN | Q16666 | TEGAEATPGAQK | y10 | 580.2831 | 929.4687 | 584.2902 | 937.4829 | 20.3 | 5.21 | 7.21 |
| IF16_HUMAN | Q16666 | TEGAEATPGAQK | y8 | 580.2831 | 801.4101 | 584.2902 | 809.4243 | 20.3 | 5.21 | 7.21 |
| IF16_HUMAN | Q16666 | TEGAEATPGAQK | y5 | 580.2831 | 500.2827 | 584.2902 | 508.2969 | 20.3 | 5.21 | 7.21 |
| IF16_HUMAN | Q16666 | TEGAEATPGAQK | b2 | 580.2831 | 231.0975 | 584.2902 | 231.0975 | 20.3 | 5.21 | 7.21 |
| IF16_HUMAN | Q16666 | TEGAEATPGAQK | b3 | 580.2831 | 288.119 | 584.2902 | 288.119 | 20.3 | 5.21 | 7.21 |
| IF16_HUMAN | Q16666 | LTC[+57.0215]FELAPK | y7 | 539.7837 | 864.4284 | 543.7908 | 872.4426 | 19.1 | 16.95 | 18.95 |
| IF16_HUMAN | Q16666 | LTC[+57.0215]FELAPK | y6 | 539.7837 | 704.3978 | 543.7908 | 712.412 | 19.1 | 16.95 | 18.95 |
| IF16_HUMAN | Q16666 | LTC[+57.0215]FELAPK | y3 | 539.7837 | 315.2027 | 543.7908 | 323.2169 | 19.1 | 16.95 | 18.95 |
| IF16_HUMAN | Q16666 | LTC[+57.0215]FELAPK | y2 | 539.7837 | 244.1656 | 543.7908 | 252.1798 | 19.1 | 16.95 | 18.95 |
| IF16_HUMAN | Q16666 | LTC[+57.0215]FELAPK | y7 | 539.7837 | 432.7178 | 543.7908 | 436.7249 | 19.1 | 16.95 | 18.95 |
| IRF4_HUMAN | Q16666 | ELTTSSPEGC[+57.0215]R | y9 | 618.7799 | 994.4258 | 623.784 | 1004.434 | 21.5 | 8.19 | 10.19 |
| IRF4_HUMAN | Q15306 | ELTTSSPEGC[+57.0215]R | y7 | 618.7799 | 792.3305 | 623.784 | 802.3387 | 21.5 | 8.19 | 10.19 |
| IRF4_HUMAN | Q15306 | ELTTSSPEGC[+57.0215]R | y6 | 618.7799 | 705.2984 | 623.784 | 715.3067 | 21.5 | 8.19 | 10.19 |
| IRF4_HUMAN | Q15306 | ELTTSSPEGC[+57.0215]R | y3 | 618.7799 | 392.1711 | 623.784 | 402.1793 | 21.5 | 8.19 | 10.19 |
| IRF4_HUMAN | Q15306 | LITAHVEPLLAR | y7 | 444.9381 | 797.488 | 448.2742 | 807.4962 | 19.2 | 17.41 | 19.41 |
| IRF4_HUMAN | Q15306 | LITAHVEPLLAR | y6 | 444.9381 | 698.4196 | 448.2742 | 708.4278 | 19.2 | 17.41 | 19.41 |
| IRF4_HUMAN | Q15306 | LITAHVEPLLAR | y5 | 444.9381 | 569.377 | 448.2742 | 579.3852 | 19.2 | 17.41 | 19.41 |
| IRF4_HUMAN | Q15306 | LITAHVEPLLAR | y1 | 444.9381 | 175.119 | 448.2742 | 185.1272 | 19.2 | 17.41 | 19.41 |
| IRF4_HUMAN | Q15306 | LITAHVEPLLAR | y8 | 444.9381 | 467.7771 | 448.2742 | 472.7812 | 19.2 | 17.41 | 19.41 |
| LCK_HUMAN | Q15306 | ITFPGLHELVR | y8 | 427.9153 | 920.5312 | 431.2514 | 930.5395 | 18.5 | 20.98 | 22.98 |

| | | | | | | | | | | |
|---|---|---|---|---|---|---|---|---|---|---|
| LCK_HUMAN | P06239 | ITFPGLHELVR | y7 | 427.9153 | 823.4785 | 431.2514 | 833.4867 | 18.5 | 20.98 | 22.98 |
| LCK_HUMAN | P06239 | ITFPGLHELVR | y5 | 427.9153 | 653.3729 | 431.2514 | 663.3812 | 18.5 | 20.98 | 22.98 |
| LCK_HUMAN | P06239 | ITFPGLHELVR | y4 | 427.9153 | 516.314 | 431.2514 | 526.3223 | 18.5 | 20.98 | 22.98 |
| LCK_HUMAN | P06239 | ITFPGLHELVR | y8 | 427.9153 | 460.7693 | 431.2514 | 465.7734 | 18.5 | 20.98 | 22.98 |
| LCK_HUMAN | P06239 | AANILVSDTLSC[+57.0215]K | y9 | 696.3636 | 1022.519 | 700.3707 | 1030.533 | 23.8 | 17.37 | 19.37 |
| LCK_HUMAN | P06239 | AANILVSDTLSC[+57.0215]K | y7 | 696.3636 | 810.3662 | 700.3707 | 818.3804 | 23.8 | 17.37 | 19.37 |
| LCK_HUMAN | P06239 | AANILVSDTLSC[+57.0215]K | y3 | 696.3636 | 394.1755 | 700.3707 | 402.1897 | 23.8 | 17.37 | 19.37 |
| LCK_HUMAN | P06239 | AANILVSDTLSC[+57.0215]K | b3 | 696.3636 | 257.1244 | 700.3707 | 257.1244 | 23.8 | 17.37 | 19.37 |
| NFKB2_HUMAN | P06239 | DSGEEAAEPSAPSR | y9 | 701.8077 | 885.4425 | 706.8118 | 895.4507 | 24 | 7.6 | 9.6 |
| NFKB2_HUMAN | Q00653 | DSGEEAAEPSAPSR | y8 | 701.8077 | 814.4054 | 706.8118 | 824.4136 | 24 | 7.6 | 9.6 |
| NFKB2_HUMAN | Q00653 | DSGEEAAEPSAPSR | y7 | 701.8077 | 743.3682 | 706.8118 | 753.3765 | 24 | 7.6 | 9.6 |
| NFKB2_HUMAN | Q00653 | DSGEEAAEPSAPSR | y6 | 701.8077 | 614.3257 | 706.8118 | 624.3339 | 24 | 7.6 | 9.6 |
| NFKB2_HUMAN | Q00653 | DSGEEAAEPSAPSR | y3 | 701.8077 | 359.2037 | 706.8118 | 369.212 | 24 | 7.6 | 9.6 |
| NFKB2_HUMAN | Q00653 | LFGLAQR | y6 | 402.74 | 691.3886 | 407.7441 | 701.3969 | 15 | 17.27 | 19.27 |
| NFKB2_HUMAN | Q00653 | LFGLAQR | y5 | 402.74 | 544.3202 | 407.7441 | 554.3284 | 15 | 17.27 | 19.27 |
| NFKB2_HUMAN | Q00653 | LFGLAQR | y4 | 402.74 | 487.2987 | 407.7441 | 497.307 | 15 | 17.27 | 19.27 |
| NFKB2_HUMAN | Q00653 | LFGLAQR | y3 | 402.74 | 374.2146 | 407.7441 | 384.2229 | 15 | 17.27 | 19.27 |
| NFKB2_HUMAN | Q00653 | LFGLAQR | y1 | 402.74 | 175.119 | 407.7441 | 185.1272 | 15 | 17.27 | 19.27 |
| NFKB2_HUMAN | Q00653 | AGAGAPELLR | y7 | 477.772 | 755.441 | 482.7761 | 765.4493 | 17.2 | 14.09 | 16.09 |
| NFKB2_HUMAN | Q00653 | AGAGAPELLR | y5 | 477.772 | 627.3824 | 482.7761 | 637.3907 | 17.2 | 14.09 | 16.09 |
| NFKB2_HUMAN | Q00653 | AGAGAPELLR | y5 | 477.772 | 314.1949 | 482.7761 | 319.199 | 17.2 | 14.09 | 16.09 |
| NFKB2_HUMAN | Q00653 | AGAGAPELLR | b2 | 477.772 | 129.0659 | 482.7761 | 129.0659 | 17.2 | 14.09 | 16.09 |
| NFKB2_HUMAN | Q00653 | AGAGAPELLR | b4 | 477.772 | 257.1244 | 482.7761 | 257.1244 | 17.2 | 14.09 | 16.09 |
| CIP2A_HUMAN | Q00653 | C[+57.0215]LEPTVALLR | y8 | 586.3288 | 898.5356 | 591.333 | 908.5439 | 20.5 | 20.02 | 22.02 |
| CIP2A_HUMAN | Q8TCG1 | C[+57.0215]LEPTVALLR | y7 | 586.3288 | 769.4931 | 591.333 | 779.5013 | 20.5 | 20.02 | 22.02 |
| CIP2A_HUMAN | Q8TCG1 | C[+57.0215]LEPTVALLR | y4 | 586.3288 | 472.3242 | 591.333 | 482.3325 | 20.5 | 20.02 | 22.02 |
| CIP2A_HUMAN | Q8TCG1 | C[+57.0215]LEPTVALLR | b2 | 586.3288 | 274.122 | 591.333 | 274.122 | 20.5 | 20.02 | 22.02 |
| CIP2A_HUMAN | Q8TCG1 | C[+57.0215]LEPTVALLR | b3 | 586.3288 | 403.1646 | 591.333 | 403.1646 | 20.5 | 20.02 | 22.02 |
| CIP2A_HUMAN | Q8TCG1 | IDLGFGTK | y6 | 425.7371 | 622.3559 | 429.7442 | 630.3701 | 15.7 | 17.33 | 19.33 |
| CIP2A_HUMAN | Q8TCG1 | IDLGFGTK | y5 | 425.7371 | 509.2718 | 429.7442 | 517.286 | 15.7 | 17.33 | 19.33 |
| CIP2A_HUMAN | Q8TCG1 | IDLGFGTK | y3 | 425.7371 | 305.1819 | 429.7442 | 313.1961 | 15.7 | 17.33 | 19.33 |
| CIP2A_HUMAN | Q8TCG1 | IDLGFGTK | b2 | 425.7371 | 229.1183 | 429.7442 | 229.1183 | 15.7 | 17.33 | 19.33 |
| CIP2A_HUMAN | Q8TCG1 | LAADVILK | y7 | 421.7709 | 729.4505 | 425.778 | 737.4647 | 15.6 | 16.04 | 18.04 |

| | | | | | | | | | | |
|---|---|---|---|---|---|---|---|---|---|---|
| CIP2A_HUMAN | Q8TCG1 | LAADVILK | y6 | 421.7709 | 658.4134 | 425.778 | 666.4276 | 15.6 | 16.04 | 18.04 |
| CIP2A_HUMAN | Q8TCG1 | LAADVILK | y2 | 421.7709 | 260.1969 | 425.778 | 268.2111 | 15.6 | 16.04 | 18.04 |
| CIP2A_HUMAN | Q8TCG1 | LAADVILK | b2 | 421.7709 | 185.1285 | 425.778 | 185.1285 | 15.6 | 16.04 | 18.04 |
| CIP2A_HUMAN | Q8TCG1 | LAADVILK | b5 | 421.7709 | 470.2609 | 425.778 | 470.2609 | 15.6 | 16.04 | 18.04 |
| CIP2A_HUMAN | Q8TCG1 | DGVPGLNIEELIEK | y11 | 763.409 | 1254.694 | 767.4161 | 1262.708 | 25.8 | 25.74 | 27.74 |
| CIP2A_HUMAN | Q8TCG1 | DGVPGLNIEELIEK | y10 | 763.409 | 1157.641 | 767.4161 | 1165.655 | 25.8 | 25.74 | 27.74 |
| CIP2A_HUMAN | Q8TCG1 | DGVPGLNIEELIEK | y6 | 763.409 | 760.4087 | 767.4161 | 768.4229 | 25.8 | 25.74 | 27.74 |
| CIP2A_HUMAN | Q8TCG1 | DGVPGLNIEELIEK | y11 | 763.409 | 627.8506 | 767.4161 | 631.8577 | 25.8 | 25.74 | 27.74 |
| CIP2A_HUMAN | Q8TCG1 | LQDLLETK | y6 | 480.274 | 718.3981 | 484.2811 | 726.4123 | 17.3 | 15.46 | 17.46 |
| CIP2A_HUMAN | Q8TCG1 | LQDLLETK | y4 | 480.274 | 490.2871 | 484.2811 | 498.3013 | 17.3 | 15.46 | 17.46 |
| CIP2A_HUMAN | Q8TCG1 | LQDLLETK | y2 | 480.274 | 248.1605 | 484.2811 | 256.1747 | 17.3 | 15.46 | 17.46 |
| CIP2A_HUMAN | Q8TCG1 | LQDLLETK | b2 | 480.274 | 242.1499 | 484.2811 | 242.1499 | 17.3 | 15.46 | 17.46 |
| CIP2A_HUMAN | Q8TCG1 | LQDLLETK | b3 | 480.274 | 357.1769 | 484.2811 | 357.1769 | 17.3 | 15.46 | 17.46 |
| CIP2A_HUMAN | Q8TCG1 | ALALAQADR | y7 | 464.7642 | 744.3999 | 469.7683 | 754.4081 | 16.8 | 11.78 | 13.78 |
| CIP2A_HUMAN | Q8TCG1 | ALALAQADR | y6 | 464.7642 | 673.3628 | 469.7683 | 683.371 | 16.8 | 11.78 | 13.78 |
| CIP2A_HUMAN | Q8TCG1 | ALALAQADR | y5 | 464.7642 | 560.2787 | 469.7683 | 570.287 | 16.8 | 11.78 | 13.78 |
| CIP2A_HUMAN | Q8TCG1 | ALALAQADR | y1 | 464.7642 | 175.119 | 469.7683 | 185.1272 | 16.8 | 11.78 | 13.78 |
| CIP2A_HUMAN | Q8TCG1 | ALALAQADR | b3 | 464.7642 | 256.1656 | 469.7683 | 256.1656 | 16.8 | 11.78 | 13.78 |
| DDX5_HUMAN | Q8TCG1 | TLSYLLPAIVHINHQPFLER | y10 | 787.7741 | 1290.67 | 791.1102 | 1300.678 | 32.2 | 24.36 | 26.36 |
| DDX5_HUMAN | P17844 | TLSYLLPAIVHINHQPFLER | y8 | 787.7741 | 1040.527 | 791.1102 | 1050.535 | 32.2 | 24.36 | 26.36 |
| DDX5_HUMAN | P17844 | TLSYLLPAIVHINHQPFLER | y5 | 787.7741 | 661.3668 | 791.1102 | 671.3751 | 32.2 | 24.36 | 26.36 |
| DDX5_HUMAN | P17844 | TLSYLLPAIVHINHQPFLER | y14 | 787.7741 | 835.9599 | 791.1102 | 840.964 | 32.2 | 24.36 | 26.36 |
| DDX5_HUMAN | P17844 | TLSYLLPAIVHINHQPFLER | b2 | 787.7741 | 215.139 | 791.1102 | 215.139 | 32.2 | 24.36 | 26.36 |
| DDX5_HUMAN | P17844 | FVINYDYPNSSEDYIHR | y10 | 711.3288 | 1217.555 | 714.6649 | 1227.563 | 29.3 | 18.86 | 20.86 |
| DDX5_HUMAN | P17844 | FVINYDYPNSSEDYIHR | y4 | 711.3288 | 588.3253 | 714.6649 | 598.3335 | 29.3 | 18.86 | 20.86 |
| DDX5_HUMAN | P17844 | FVINYDYPNSSEDYIHR | y10 | 711.3288 | 609.2809 | 714.6649 | 614.285 | 29.3 | 18.86 | 20.86 |
| DDX5_HUMAN | P17844 | FVINYDYPNSSEDYIHR | b2 | 711.3288 | 247.1441 | 714.6649 | 247.1441 | 29.3 | 18.86 | 20.86 |
| DDX5_HUMAN | P17844 | TGTAYTFFTPNNIK | y9 | 787.8961 | 1081.568 | 791.9032 | 1089.582 | 26.5 | 20.61 | 22.61 |
| DDX5_HUMAN | P17844 | TGTAYTFFTPNNIK | y8 | 787.8961 | 980.52 | 791.9032 | 988.5342 | 26.5 | 20.61 | 22.61 |
| DDX5_HUMAN | P17844 | TGTAYTFFTPNNIK | y7 | 787.8961 | 833.4516 | 791.9032 | 841.4658 | 26.5 | 20.61 | 22.61 |
| DDX5_HUMAN | P17844 | TGTAYTFFTPNNIK | y6 | 787.8961 | 686.3832 | 791.9032 | 694.3974 | 26.5 | 20.61 | 22.61 |
| DDX5_HUMAN | P17844 | TGTAYTFFTPNNIK | y5 | 787.8961 | 585.3355 | 791.9032 | 593.3497 | 26.5 | 20.61 | 22.61 |
| DOCK8_HUMAN | P17844 | LQAESFC[+57.0215]QR | y7 | 569.7691 | 897.3883 | 574.7733 | 907.3966 | 20 | 11.37 | 13.37 |

| | | | | | | | | | | |
|---|---|---|---|---|---|---|---|---|---|---|
| DOCK8_HUMAN | Q8NF50 | LQAESFC[+57.0215]QR | y6 | 569.7691 | 826.3512 | 574.7733 | 836.3595 | 20 | 11.37 | 13.37 |
| DOCK8_HUMAN | Q8NF50 | LQAESFC[+57.0215]QR | y5 | 569.7691 | 697.3086 | 574.7733 | 707.3169 | 20 | 11.37 | 13.37 |
| DOCK8_HUMAN | Q8NF50 | LQAESFC[+57.0215]QR | y2 | 569.7691 | 303.1775 | 574.7733 | 313.1858 | 20 | 11.37 | 13.37 |
| DOCK8_HUMAN | Q8NF50 | LQAESFC[+57.0215]QR | b2 | 569.7691 | 242.1499 | 574.7733 | 242.1499 | 20 | 11.37 | 13.37 |
| DOCK8_HUMAN | Q8NF50 | SPDFYEEVK | y7 | 557.2586 | 929.4251 | 561.2657 | 937.4393 | 19.6 | 15.33 | 17.33 |
| DOCK8_HUMAN | Q8NF50 | SPDFYEEVK | y6 | 557.2586 | 814.3981 | 561.2657 | 822.4123 | 19.6 | 15.33 | 17.33 |
| DOCK8_HUMAN | Q8NF50 | SPDFYEEVK | y5 | 557.2586 | 667.3297 | 561.2657 | 675.3439 | 19.6 | 15.33 | 17.33 |
| DOCK8_HUMAN | Q8NF50 | SPDFYEEVK | y2 | 557.2586 | 246.1812 | 561.2657 | 254.1954 | 19.6 | 15.33 | 17.33 |
| DOCK8_HUMAN | Q8NF50 | TSGIVLSSLPYK | y8 | 632.861 | 906.5295 | 636.8681 | 914.5437 | 21.9 | 19.36 | 21.36 |
| DOCK8_HUMAN | Q8NF50 | TSGIVLSSLPYK | y7 | 632.861 | 807.4611 | 636.8681 | 815.4753 | 21.9 | 19.36 | 21.36 |
| DOCK8_HUMAN | Q8NF50 | TSGIVLSSLPYK | y6 | 632.861 | 694.377 | 636.8681 | 702.3912 | 21.9 | 19.36 | 21.36 |
| DOCK8_HUMAN | Q8NF50 | TSGIVLSSLPYK | y3 | 632.861 | 407.2289 | 636.8681 | 415.2431 | 21.9 | 19.36 | 21.36 |
| DOCK8_HUMAN | Q8NF50 | TSGIVLSSLPYK | b3 | 632.861 | 246.1084 | 636.8681 | 246.1084 | 21.9 | 19.36 | 21.36 |
| DOCK8_HUMAN | Q8NF50 | LVIPILEAHR | y5 | 387.5765 | 625.3416 | 390.9126 | 635.3499 | 17 | 21.26 | 23.26 |
| DOCK8_HUMAN | Q8NF50 | LVIPILEAHR | y4 | 387.5765 | 512.2576 | 390.9126 | 522.2658 | 17 | 21.26 | 23.26 |
| DOCK8_HUMAN | Q8NF50 | LVIPILEAHR | y7 | 387.5765 | 418.2429 | 390.9126 | 423.247 | 17 | 21.26 | 23.26 |
| DOCK8_HUMAN | Q8NF50 | LVIPILEAHR | b2 | 387.5765 | 213.1598 | 390.9126 | 213.1598 | 17 | 21.26 | 23.26 |
| UBR5_HUMAN | Q8NF50 | VLLLPLER | y6 | 476.8131 | 740.4665 | 481.8173 | 750.4748 | 17.2 | 22.07 | 24.07 |
| UBR5_HUMAN | O95071 | VLLLPLER | y5 | 476.8131 | 627.3824 | 481.8173 | 637.3907 | 17.2 | 22.07 | 24.07 |
| UBR5_HUMAN | O95071 | VLLLPLER | y4 | 476.8131 | 514.2984 | 481.8173 | 524.3066 | 17.2 | 22.07 | 24.07 |
| UBR5_HUMAN | O95071 | VLLLPLER | y1 | 476.8131 | 175.119 | 481.8173 | 185.1272 | 17.2 | 22.07 | 24.07 |
| UBR5_HUMAN | O95071 | VLLLPLER | b2 | 476.8131 | 213.1598 | 481.8173 | 213.1598 | 17.2 | 22.07 | 24.07 |
| UBR5_HUMAN | O95071 | NVAIFTAGQESPIILR | y12 | 864.9858 | 1331.732 | 869.9899 | 1341.74 | 28.9 | 22.84 | 24.84 |
| UBR5_HUMAN | O95071 | NVAIFTAGQESPIILR | y11 | 864.9858 | 1184.663 | 869.9899 | 1194.672 | 28.9 | 22.84 | 24.84 |
| UBR5_HUMAN | O95071 | NVAIFTAGQESPIILR | y9 | 864.9858 | 1012.579 | 869.9899 | 1022.587 | 28.9 | 22.84 | 24.84 |
| UBR5_HUMAN | O95071 | NVAIFTAGQESPIILR | b2 | 864.9858 | 214.1186 | 869.9899 | 214.1186 | 28.9 | 22.84 | 24.84 |
| UBR5_HUMAN | O95071 | NVAIFTAGQESPIILR | b3 | 864.9858 | 285.1557 | 869.9899 | 285.1557 | 28.9 | 22.84 | 24.84 |
| UBR5_HUMAN | O95071 | LLC[+57.0215]DSVVLQPYLR | y10 | 788.4318 | 1189.658 | 793.4359 | 1199.666 | 26.6 | 22.3 | 24.3 |
| UBR5_HUMAN | O95071 | LLC[+57.0215]DSVVLQPYLR | y7 | 788.4318 | 888.5302 | 793.4359 | 898.5384 | 26.6 | 22.3 | 24.3 |
| UBR5_HUMAN | O95071 | LLC[+57.0215]DSVVLQPYLR | y6 | 788.4318 | 789.4618 | 793.4359 | 799.47 | 26.6 | 22.3 | 24.3 |
| UBR5_HUMAN | O95071 | LLC[+57.0215]DSVVLQPYLR | y5 | 788.4318 | 676.3777 | 793.4359 | 686.386 | 26.6 | 22.3 | 24.3 |
| UBR5_HUMAN | O95071 | LLC[+57.0215]DSVVLQPYLR | y4 | 788.4318 | 548.3191 | 793.4359 | 558.3274 | 26.6 | 22.3 | 24.3 |
| UBR5_HUMAN | O95071 | LLC[+57.0215]DSVVLQPYLR | y6 | 525.957 | 789.4618 | 529.293 | 799.47 | 22.3 | 22.3 | 24.3 |

| | | | | | | | | | | |
|---|---|---|---|---|---|---|---|---|---|---|
| UBR5_HUMAN | O95071 | LLC[+57.0215]DSVVLQPYLR | y5 | 525.957 | 676.3777 | 529.293 | 686.386 | 22.3 | 22.3 | 24.3 |
| UBR5_HUMAN | O95071 | LLC[+57.0215]DSVVLQPYLR | y4 | 525.957 | 548.3191 | 529.293 | 558.3274 | 22.3 | 22.3 | 24.3 |
| UBR5_HUMAN | O95071 | LLC[+57.0215]DSVVLQPYLR | y1 | 525.957 | 175.119 | 529.293 | 185.1272 | 22.3 | 22.3 | 24.3 |
| UBR5_HUMAN | O95071 | LLC[+57.0215]DSVVLQPYLR | b6 | 525.957 | 688.3334 | 529.293 | 688.3334 | 22.3 | 22.3 | 24.3 |
| UBR5_HUMAN | O95071 | AYPAAITILETAQK | y10 | 745.4167 | 1087.636 | 749.4238 | 1095.65 | 25.3 | 22.06 | 24.06 |
| UBR5_HUMAN | O95071 | AYPAAITILETAQK | y9 | 745.4167 | 1016.599 | 749.4238 | 1024.613 | 25.3 | 22.06 | 24.06 |
| UBR5_HUMAN | O95071 | AYPAAITILETAQK | y8 | 745.4167 | 903.5146 | 749.4238 | 911.5288 | 25.3 | 22.06 | 24.06 |
| UBR5_HUMAN | O95071 | AYPAAITILETAQK | y6 | 745.4167 | 689.3828 | 749.4238 | 697.397 | 25.3 | 22.06 | 24.06 |
| UBR5_HUMAN | O95071 | AYPAAITILETAQK | y4 | 745.4167 | 447.2562 | 749.4238 | 455.2704 | 25.3 | 22.06 | 24.06 |
| RUNX1_HUMAN | O95071 | TDSPNFLC[+57.0215]SVLPTHWR | y8 | 643.9807 | 995.5421 | 647.3168 | 1005.55 | 26.8 | 24.53 | 26.53 |
| RUNX1_HUMAN | Q01196 | TDSPNFLC[+57.0215]SVLPTHWR | y6 | 643.9807 | 809.4417 | 647.3168 | 819.45 | 26.8 | 24.53 | 26.53 |
| RUNX1_HUMAN | Q01196 | TDSPNFLC[+57.0215]SVLPTHWR | y5 | 643.9807 | 696.3576 | 647.3168 | 706.3659 | 26.8 | 24.53 | 26.53 |
| RUNX1_HUMAN | Q01196 | TDSPNFLC[+57.0215]SVLPTHWR | y5 | 643.9807 | 348.6824 | 647.3168 | 353.6866 | 26.8 | 24.53 | 26.53 |
| RUNX1_HUMAN | Q01196 | TDSPNFLC[+57.0215]SVLPTHWR | b2 | 643.9807 | 217.0819 | 647.3168 | 217.0819 | 26.8 | 24.53 | 26.53 |
| RUNX1_HUMAN | Q01196 | LSTAPDLTAFSDPR | y10 | 745.8779 | 1118.548 | 750.882 | 1128.556 | 25.3 | 18.88 | 20.88 |
| RUNX1_HUMAN | Q01196 | LSTAPDLTAFSDPR | y8 | 745.8779 | 906.468 | 750.882 | 916.4762 | 25.3 | 18.88 | 20.88 |
| RUNX1_HUMAN | Q01196 | LSTAPDLTAFSDPR | y6 | 745.8779 | 692.3362 | 750.882 | 702.3445 | 25.3 | 18.88 | 20.88 |
| RUNX1_HUMAN | Q01196 | LSTAPDLTAFSDPR | y2 | 745.8779 | 272.1717 | 750.882 | 282.18 | 25.3 | 18.88 | 20.88 |
| RUNX3_HUMAN | Q01196 | IPVDPSTSR | y7 | 486.2615 | 761.3788 | 491.2656 | 771.3871 | 17.5 | 9.17 | 11.17 |
| RUNX3_HUMAN | Q13761 | IPVDPSTSR | y6 | 486.2615 | 662.3104 | 491.2656 | 672.3187 | 17.5 | 9.17 | 11.17 |
| RUNX3_HUMAN | Q13761 | IPVDPSTSR | y5 | 486.2615 | 547.2835 | 491.2656 | 557.2917 | 17.5 | 9.17 | 11.17 |
| RUNX3_HUMAN | Q13761 | IPVDPSTSR | y8 | 486.2615 | 429.7194 | 491.2656 | 434.7236 | 17.5 | 9.17 | 11.17 |
| RUNX3_HUMAN | Q13761 | VTPSTPSPR | y7 | 471.2562 | 741.389 | 476.2603 | 751.3972 | 17 | 7.09 | 9.09 |
| RUNX3_HUMAN | Q13761 | VTPSTPSPR | y6 | 471.2562 | 644.3362 | 476.2603 | 654.3445 | 17 | 7.09 | 9.09 |
| RUNX3_HUMAN | Q13761 | VTPSTPSPR | y5 | 471.2562 | 557.3042 | 476.2603 | 567.3125 | 17 | 7.09 | 9.09 |
| RUNX3_HUMAN | Q13761 | VTPSTPSPR | y4 | 471.2562 | 456.2565 | 476.2603 | 466.2648 | 17 | 7.09 | 9.09 |
| RUNX3_HUMAN | Q13761 | VTPSTPSPR | y7 | 471.2562 | 371.1981 | 476.2603 | 376.2023 | 17 | 7.09 | 9.09 |
| LMNB1_HUMAN | Q13761 | AGGPTTPLSPTR | y8 | 577.8118 | 872.4836 | 582.816 | 882.4919 | 20.2 | 11 | 13 |
| LMNB1_HUMAN | P20700 | AGGPTTPLSPTR | y7 | 577.8118 | 771.4359 | 582.816 | 781.4442 | 20.2 | 11 | 13 |
| LMNB1_HUMAN | P20700 | AGGPTTPLSPTR | y6 | 577.8118 | 670.3883 | 582.816 | 680.3965 | 20.2 | 11 | 13 |
| LMNB1_HUMAN | P20700 | AGGPTTPLSPTR | y4 | 577.8118 | 460.2514 | 582.816 | 470.2597 | 20.2 | 11 | 13 |
| LMNB1_HUMAN | P20700 | ALYETELADAR | y9 | 626.3144 | 1067.5 | 631.3186 | 1077.509 | 21.7 | 15.75 | 17.75 |
| LMNB1_HUMAN | P20700 | ALYETELADAR | y8 | 626.3144 | 904.4371 | 631.3186 | 914.4453 | 21.7 | 15.75 | 17.75 |

| | | | | | | | | | | |
|---|---|---|---|---|---|---|---|---|---|---|
| LMNB1_HUMAN | P20700 | ALYETELADAR | y7 | 626.3144 | 775.3945 | 631.3186 | 785.4027 | 21.7 | 15.75 | 17.75 |
| LMNB1_HUMAN | P20700 | ALYETELADAR | y2 | 626.3144 | 246.1561 | 631.3186 | 256.1643 | 21.7 | 15.75 | 17.75 |
| MX2_HUMAN | P20700 | VLLEEGSATVPR | y10 | 635.8537 | 1058.548 | 640.8578 | 1068.556 | 22 | 14.91 | 16.91 |
| MX2_HUMAN | P20592 | VLLEEGSATVPR | y8 | 635.8537 | 816.421 | 640.8578 | 826.4293 | 22 | 14.91 | 16.91 |
| MX2_HUMAN | P20592 | VLLEEGSATVPR | y7 | 635.8537 | 687.3784 | 640.8578 | 697.3867 | 22 | 14.91 | 16.91 |
| MX2_HUMAN | P20592 | HALC[+57.0215]QFSSK | y8 | 539.2609 | 940.4557 | 543.268 | 948.4699 | 19.1 | 8.68 | 10.68 |
| MX2_HUMAN | P20592 | HALC[+57.0215]QFSSK | y7 | 539.2609 | 869.4186 | 543.268 | 877.4328 | 19.1 | 8.68 | 10.68 |
| MX2_HUMAN | P20592 | HALC[+57.0215]QFSSK | y2 | 539.2609 | 234.1448 | 543.268 | 242.159 | 19.1 | 8.68 | 10.68 |
| MX2_HUMAN | P20592 | HALC[+57.0215]QFSSK | b2 | 539.2609 | 209.1033 | 543.268 | 209.1033 | 19.1 | 8.68 | 10.68 |
| STAT1_HUMAN | P20592 | FHDLLSQLDDQYSR | y7 | 579.6128 | 896.4108 | 582.9489 | 906.4191 | 24.3 | 20.75 | 22.75 |
| STAT1_HUMAN | P42224 | FHDLLSQLDDQYSR | y6 | 579.6128 | 783.3268 | 582.9489 | 793.335 | 24.3 | 20.75 | 22.75 |
| STAT1_HUMAN | P42224 | FHDLLSQLDDQYSR | y5 | 579.6128 | 668.2998 | 582.9489 | 678.3081 | 24.3 | 20.75 | 22.75 |
| STAT1_HUMAN | P42224 | FHDLLSQLDDQYSR | y4 | 579.6128 | 553.2729 | 582.9489 | 563.2812 | 24.3 | 20.75 | 22.75 |
| STAT1_HUMAN | P42224 | FHDLLSQLDDQYSR | b3 | 579.6128 | 400.1615 | 582.9489 | 400.1615 | 24.3 | 20.75 | 22.75 |
| STAT1_HUMAN | P42224 | SLEDLQDEYDFK | y10 | 751.3383 | 1301.553 | 755.3454 | 1309.567 | 25.4 | 19.32 | 21.32 |
| STAT1_HUMAN | P42224 | SLEDLQDEYDFK | y9 | 751.3383 | 1172.511 | 755.3454 | 1180.525 | 25.4 | 19.32 | 21.32 |
| STAT1_HUMAN | P42224 | SLEDLQDEYDFK | y6 | 751.3383 | 816.341 | 755.3454 | 824.3552 | 25.4 | 19.32 | 21.32 |
| STAT1_HUMAN | P42224 | SLEDLQDEYDFK | y2 | 751.3383 | 294.1812 | 755.3454 | 302.1954 | 25.4 | 19.32 | 21.32 |
| STAT1_HUMAN | P42224 | SLEDLQDEYDFK | b2 | 751.3383 | 201.1234 | 755.3454 | 201.1234 | 25.4 | 19.32 | 21.32 |
| STAT1_HUMAN | P42224 | TELISVSEVHPSR | y9 | 485.2597 | 997.5061 | 488.5958 | 1007.514 | 20.7 | 14.84 | 16.84 |
| STAT1_HUMAN | P42224 | TELISVSEVHPSR | y7 | 485.2597 | 811.4057 | 488.5958 | 821.414 | 20.7 | 14.84 | 16.84 |
| STAT1_HUMAN | P42224 | TELISVSEVHPSR | y4 | 485.2597 | 496.2627 | 488.5958 | 506.2709 | 20.7 | 14.84 | 16.84 |
| STAT1_HUMAN | P42224 | TELISVSEVHPSR | y3 | 485.2597 | 359.2037 | 488.5958 | 369.212 | 20.7 | 14.84 | 16.84 |
| STAT1_HUMAN | P42224 | TELISVSEVHPSR | b2 | 485.2597 | 231.0975 | 488.5958 | 231.0975 | 20.7 | 14.84 | 16.84 |
| 2AAA_HUMAN | P42224 | LAGGDWFTSR | y9 | 555.2724 | 996.4534 | 560.2765 | 1006.462 | 19.6 | 18.97 | 20.97 |
| 2AAA_HUMAN | P30153 | LAGGDWFTSR | y8 | 555.2724 | 925.4163 | 560.2765 | 935.4245 | 19.6 | 18.97 | 20.97 |
| 2AAA_HUMAN | P30153 | LAGGDWFTSR | y5 | 555.2724 | 696.3464 | 560.2765 | 706.3547 | 19.6 | 18.97 | 20.97 |
| 2AAA_HUMAN | P30153 | LAGGDWFTSR | y4 | 555.2724 | 510.2671 | 560.2765 | 520.2753 | 19.6 | 18.97 | 20.97 |
| PP2AA_HUMAN | P30153 | ELDQWIEQLNEC[+57.0215]K | y8 | 852.8985 | 1033.498 | 856.9056 | 1041.512 | 28.5 | 22.87 | 24.87 |
| PP2AA_HUMAN | P67775 | ELDQWIEQLNEC[+57.0215]K | y7 | 852.8985 | 920.4142 | 856.9056 | 928.4284 | 28.5 | 22.87 | 24.87 |
| PP2AA_HUMAN | P67775 | ELDQWIEQLNEC[+57.0215]K | y4 | 852.8985 | 550.229 | 856.9056 | 558.2432 | 28.5 | 22.87 | 24.87 |
| PP2AA_HUMAN | P67775 | ELDQWIEQLNEC[+57.0215]K | y2 | 852.8985 | 307.1435 | 856.9056 | 315.1577 | 28.5 | 22.87 | 24.87 |
| PP2AA_HUMAN | P67775 | ELDQWIEQLNEC[+57.0215]K | b4 | 852.8985 | 243.6134 | 856.9056 | 243.6134 | 28.5 | 22.87 | 24.87 |

| | | | | | | | | | | |
|---|---|---|---|---|---|---|---|---|---|---|
| PP2AA_HUMAN | P67775 | GGWGISPR | y5 | 415.2194 | 529.3093 | 420.2235 | 539.3175 | 15.4 | 14.39 | 16.39 |
| PP2AA_HUMAN | P67775 | GGWGISPR | y4 | 415.2194 | 472.2878 | 420.2235 | 482.2961 | 15.4 | 14.39 | 16.39 |
| PP2AA_HUMAN | P67775 | GGWGISPR | y3 | 415.2194 | 359.2037 | 420.2235 | 369.212 | 15.4 | 14.39 | 16.39 |
| PP2AA_HUMAN | P67775 | GGWGISPR | y2 | 415.2194 | 272.1717 | 420.2235 | 282.18 | 15.4 | 14.39 | 16.39 |
| PP2AB_HUMAN | P67775 | GGWGISPR | y5 | 415.2194 | 529.3093 | 420.2235 | 539.3175 | 15.4 | 14.39 | 16.39 |
| PP2AB_HUMAN | P62714 | GGWGISPR | y4 | 415.2194 | 472.2878 | 420.2235 | 482.2961 | 15.4 | 14.39 | 16.39 |
| PP2AB_HUMAN | P62714 | GGWGISPR | y3 | 415.2194 | 359.2037 | 420.2235 | 369.212 | 15.4 | 14.39 | 16.39 |
| PP2AB_HUMAN | P62714 | GGWGISPR | y2 | 415.2194 | 272.1717 | 420.2235 | 282.18 | 15.4 | 14.39 | 16.39 |
| PP1B_HUMAN | P62714 | HDLDLIC[+57.0215]R | y7 | 521.2609 | 904.4557 | 526.2651 | 914.4639 | 18.5 | 13.99 | 15.99 |
| PP1B_HUMAN | P62140 | HDLDLIC[+57.0215]R | y6 | 521.2609 | 789.4287 | 526.2651 | 799.437 | 18.5 | 13.99 | 15.99 |
| PP1B_HUMAN | P62140 | HDLDLIC[+57.0215]R | y5 | 521.2609 | 676.3447 | 526.2651 | 686.3529 | 18.5 | 13.99 | 15.99 |
| PP1B_HUMAN | P62140 | HDLDLIC[+57.0215]R | y4 | 521.2609 | 561.3177 | 526.2651 | 571.326 | 18.5 | 13.99 | 15.99 |
| PP1B_HUMAN | P62140 | HDLDLIC[+57.0215]R | b2 | 521.2609 | 253.0931 | 526.2651 | 253.0931 | 18.5 | 13.99 | 15.99 |
| PP1A_HUMAN | P62140 | LLEVQGSRPGK | y8 | 395.2313 | 828.4686 | 397.9027 | 836.4828 | 17.3 | 9.65 | 11.65 |
| PP1A_HUMAN | P62136 | LLEVQGSRPGK | y7 | 395.2313 | 729.4002 | 397.9027 | 737.4144 | 17.3 | 9.65 | 11.65 |
| PP1A_HUMAN | P62136 | LLEVQGSRPGK | y6 | 395.2313 | 601.3416 | 397.9027 | 609.3558 | 17.3 | 9.65 | 11.65 |
| PP1A_HUMAN | P62136 | LLEVQGSRPGK | y3 | 395.2313 | 301.187 | 397.9027 | 309.2012 | 17.3 | 9.65 | 11.65 |
| PP1A_HUMAN | P62136 | HDLDLIC[+57.0215]R | y7 | 521.2609 | 904.4557 | 526.2651 | 914.4639 | 18.5 | 13.99 | 15.99 |
| PP1A_HUMAN | P62136 | HDLDLIC[+57.0215]R | y6 | 521.2609 | 789.4287 | 526.2651 | 799.437 | 18.5 | 13.99 | 15.99 |
| PP1A_HUMAN | P62136 | HDLDLIC[+57.0215]R | y5 | 521.2609 | 676.3447 | 526.2651 | 686.3529 | 18.5 | 13.99 | 15.99 |
| PP1A_HUMAN | P62136 | HDLDLIC[+57.0215]R | y4 | 521.2609 | 561.3177 | 526.2651 | 571.326 | 18.5 | 13.99 | 15.99 |
| PP1A_HUMAN | P62136 | HDLDLIC[+57.0215]R | b2 | 521.2609 | 253.0931 | 526.2651 | 253.0931 | 18.5 | 13.99 | 15.99 |
| STAT3_HUMAN | P62136 | GLSIEQLTTLAEK | y9 | 701.893 | 1032.557 | 705.9001 | 1040.571 | 24 | 22.79 | 24.79 |
| STAT3_HUMAN | P40763 | GLSIEQLTTLAEK | y7 | 701.893 | 775.456 | 705.9001 | 783.4702 | 24 | 22.79 | 24.79 |
| STAT3_HUMAN | P40763 | GLSIEQLTTLAEK | y6 | 701.893 | 662.3719 | 705.9001 | 670.3861 | 24 | 22.79 | 24.79 |
| STAT3_HUMAN | P40763 | GLSIEQLTTLAEK | b2 | 701.893 | 171.1128 | 705.9001 | 171.1128 | 24 | 22.79 | 24.79 |
| STAT3_HUMAN | P40763 | GLSIEQLTTLAEK | b3 | 701.893 | 258.1448 | 705.9001 | 258.1448 | 24 | 22.79 | 24.79 |
| STAT3_HUMAN | P40763 | EGGVTFTWVEK | y7 | 626.8141 | 910.4669 | 630.8212 | 918.4811 | 21.7 | 20.04 | 22.04 |
| STAT3_HUMAN | P40763 | EGGVTFTWVEK | y6 | 626.8141 | 809.4192 | 630.8212 | 817.4334 | 21.7 | 20.04 | 22.04 |
| RO52_HUMAN | P40763 | GGGSVC[+57.0215]PVC[+57.0215]R | y6 | 524.7368 | 790.3698 | 529.7409 | 800.3781 | 18.6 | 7.59 | 9.59 |
| RO52_HUMAN | P19474 | GGGSVC[+57.0215]PVC[+57.0215]R | y5 | 524.7368 | 691.3014 | 529.7409 | 701.3097 | 18.6 | 7.59 | 9.59 |
| RO52_HUMAN | P19474 | GGGSVC[+57.0215]PVC[+57.0215]R | y4 | 524.7368 | 531.2708 | 529.7409 | 541.279 | 18.6 | 7.59 | 9.59 |
| RO52_HUMAN | P19474 | GGGSVC[+57.0215]PVC[+57.0215]R | b4 | 524.7368 | 259.1037 | 529.7409 | 259.1037 | 18.6 | 7.59 | 9.59 |

| | | | | | | | | | | |
|---|---|---|---|---|---|---|---|---|---|---|
| RO52_HUMAN | P19474 | LQVALGELR | y7 | 499.8033 | 757.4567 | 504.8074 | 767.4649 | 17.9 | 18.42 | 20.42 |
| RO52_HUMAN | P19474 | LQVALGELR | y6 | 499.8033 | 658.3883 | 504.8074 | 668.3965 | 17.9 | 18.42 | 20.42 |
| RO52_HUMAN | P19474 | LQVALGELR | y4 | 499.8033 | 474.2671 | 504.8074 | 484.2753 | 17.9 | 18.42 | 20.42 |
| RO52_HUMAN | P19474 | LQVALGELR | y2 | 499.8033 | 288.203 | 504.8074 | 298.2113 | 17.9 | 18.42 | 20.42 |
| RO52_HUMAN | P19474 | LQVALGELR | y4 | 499.8033 | 237.6372 | 504.8074 | 242.6413 | 17.9 | 18.42 | 20.42 |
| MSRT1_SYNTH | P19474 | RGDSPASSPK | y2 | 505.2613 | 252.1798 | 505.2613 | 252.1798 | 17.9 | 4.43 | 6.43 |
| MSRT1_SYNTH | MSRT1_S | RGDSPASSPK | b3 | 505.2613 | 329.1568 | 505.2613 | 329.1568 | 17.9 | 4.43 | 6.43 |
| MSRT1_SYNTH | MSRT1_S | RGDSPASSPK | b8 | 505.2613 | 758.3428 | 505.2613 | 758.3428 | 17.9 | 4.43 | 6.43 |
| MSRT1_SYNTH | MSRT1_S | LGGNETQVR | y8 | 492.2608 | 870.4303 | 492.2608 | 870.4303 | 17.5 | 7.15 | 9.15 |
| MSRT1_SYNTH | MSRT1_S | LGGNETQVR | y7 | 492.2608 | 813.4089 | 492.2608 | 813.4089 | 17.5 | 7.15 | 9.15 |
| MSRT1_SYNTH | MSRT1_S | LGGNETQVR | y4 | 492.2608 | 513.3019 | 492.2608 | 513.3019 | 17.5 | 7.15 | 9.15 |
| MSRT1_SYNTH | MSRT1_S | AEFAEVSK | y6 | 444.7313 | 688.3756 | 444.7313 | 688.3756 | 16.1 | 10.23 | 12.23 |
| MSRT1_SYNTH | MSRT1_S | AEFAEVSK | y5 | 444.7313 | 541.3072 | 444.7313 | 541.3072 | 16.1 | 10.23 | 12.23 |
| MSRT1_SYNTH | MSRT1_S | AEFAEVSK | b3 | 444.7313 | 348.1554 | 444.7313 | 348.1554 | 16.1 | 10.23 | 12.23 |
| MSRT1_SYNTH | MSRT1_S | SGFSSVSVSR | y7 | 511.7607 | 731.3922 | 511.7607 | 731.3922 | 18.1 | 12.07 | 14.07 |
| MSRT1_SYNTH | MSRT1_S | SGFSSVSVSR | y6 | 511.7607 | 644.3601 | 511.7607 | 644.3601 | 18.1 | 12.07 | 14.07 |
| MSRT1_SYNTH | MSRT1_S | SGFSSVSVSR | y4 | 511.7607 | 458.2597 | 511.7607 | 458.2597 | 18.1 | 12.07 | 14.07 |
| MSRT1_SYNTH | MSRT1_S | ADEGISFR | y5 | 452.7236 | 589.3332 | 452.7236 | 589.3332 | 16.3 | 13.42 | 15.42 |
| MSRT1_SYNTH | MSRT1_S | ADEGISFR | y3 | 452.7236 | 419.2277 | 452.7236 | 419.2277 | 16.3 | 13.42 | 15.42 |
| MSRT1_SYNTH | MSRT1_S | ADEGISFR | b2 | 452.7236 | 187.0713 | 452.7236 | 187.0713 | 16.3 | 13.42 | 15.42 |
| MSRT1_SYNTH | MSRT1_S | DISLSDYK | y6 | 474.7418 | 720.3654 | 474.7418 | 720.3654 | 17 | 14.99 | 16.99 |
| MSRT1_SYNTH | MSRT1_S | DISLSDYK | y2 | 474.7418 | 318.1903 | 474.7418 | 318.1903 | 17 | 14.99 | 16.99 |
| MSRT1_SYNTH | MSRT1_S | DISLSDYK | b2 | 474.7418 | 229.1183 | 474.7418 | 229.1183 | 17 | 14.99 | 16.99 |
| MSRT1_SYNTH | MSRT1_S | LVNEVTEFAK | y8 | 579.3182 | 945.4767 | 579.3182 | 945.4767 | 20.2 | 16.25 | 18.25 |
| MSRT1_SYNTH | MSRT1_S | LVNEVTEFAK | y6 | 579.3182 | 702.3912 | 579.3182 | 702.3912 | 20.2 | 16.25 | 18.25 |
| MSRT1_SYNTH | MSRT1_S | LVNEVTEFAK | b2 | 579.3182 | 213.1598 | 579.3182 | 213.1598 | 20.2 | 16.25 | 18.25 |
| MSRT1_SYNTH | MSRT1_S | DQGGELLSLR | y8 | 549.2949 | 854.497 | 549.2949 | 854.497 | 19.2 | 17.95 | 19.95 |
| MSRT1_SYNTH | MSRT1_S | DQGGELLSLR | y3 | 549.2949 | 385.2433 | 549.2949 | 385.2433 | 19.2 | 17.95 | 19.95 |
| MSRT1_SYNTH | MSRT1_S | DQGGELLSLR | b2 | 549.2949 | 244.0928 | 549.2949 | 244.0928 | 19.2 | 17.95 | 19.95 |
| MSRT1_SYNTH | MSRT1_S | GLFIIDDK | y6 | 464.7651 | 758.4174 | 464.7651 | 758.4174 | 16.7 | 20.39 | 22.39 |
| MSRT1_SYNTH | MSRT1_S | GLFIIDDK | y5 | 464.7651 | 611.349 | 464.7651 | 611.349 | 16.7 | 20.39 | 22.39 |
| MSRT1_SYNTH | MSRT1_S | GLFIIDDK | y4 | 464.7651 | 498.265 | 464.7651 | 498.265 | 16.7 | 20.39 | 22.39 |
| MSRT1_SYNTH | MSRT1_S | YWGVASFLQK | y8 | 603.8235 | 857.4971 | 603.8235 | 857.4971 | 20.9 | 23.88 | 25.88 |

| | | | | | | | | | | |
|---|---|---|---|---|---|---|---|---|---|---|
| MSRT1_SYNTH | MSRT1_S | YWGVASFLQK | y6 | 603.8235 | 701.4072 | 603.8235 | 701.4072 | 20.9 | 23.88 | 25.88 |
| MSRT1_SYNTH | MSRT1_S | YWGVASFLQK | b2 | 603.8235 | 350.1499 | 603.8235 | 350.1499 | 20.9 | 23.88 | 25.88 |
| MSRT1_SYNTH | MSRT1_S | TDELFQIEGLKEELAYLR | y7 | 726.3836 | 903.481 | 726.3836 | 903.481 | 29.8 | 26.74 | 28.74 |
| MSRT1_SYNTH | MSRT1_S | TDELFQIEGLKEELAYLR | y14 | 726.3836 | 859.9712 | 726.3836 | 859.9712 | 29.8 | 26.74 | 28.74 |
| MSRT1_SYNTH | MSRT1_S | TDELFQIEGLKEELAYLR | b2 | 726.3836 | 217.0819 | 726.3836 | 217.0819 | 29.8 | 26.74 | 28.74 |
| MSRT1_SYNTH | MSRT1_S | TDELFQIEGLKEELAYLR | b3 | 726.3836 | 346.1245 | 726.3836 | 346.1245 | 29.8 | 26.74 | 28.74 |
| MSRT1_SYNTH | MSRT1_S | AVQQPDGLAVLGIFLK | y12 | 838.9949 | 1250.76 | 838.9949 | 1250.76 | 28 | 30.49 | 32.49 |
| MSRT1_SYNTH | MSRT1_S | AVQQPDGLAVLGIFLK | y8 | 838.9949 | 868.5746 | 838.9949 | 868.5746 | 28 | 30.49 | 32.49 |
| MSRT1_SYNTH | MSRT1_S | AVQQPDGLAVLGIFLK | y6 | 838.9949 | 698.4691 | 838.9949 | 698.4691 | 28 | 30.49 | 32.49 |
| MSRT1_SYNTH | MSRT1_S | AVQQPDGLAVLGIFLK | b2 | 838.9949 | 171.1128 | 838.9949 | 171.1128 | 28 | 30.49 | 32.49 |
| MSRT1_SYNTH | MSRT1_S | LGEYGFQNAL | y2 | 559.7831 | 210.1562 | 559.7831 | 210.1562 | 19.6 | 21.61 | 23.61 |
| MSRT1_SYNTH | MSRT1_S | LGEYGFQNAL | y1 | 559.7831 | 139.1191 | 559.7831 | 139.1191 | 19.6 | 21.61 | 23.61 |
| MSRT1_SYNTH | MSRT1_S | LGEYGFQNAL | b2 | 559.7831 | 171.1128 | 559.7831 | 171.1128 | 19.6 | 21.61 | 23.61 |
| MSRT1_SYNTH | MSRT1_S | LGEYGFQNAL | b5 | 559.7831 | 520.2402 | 559.7831 | 520.2402 | 19.6 | 21.61 | 23.61 |
