## Supplementary Table 5 for "Protein Interactome of the Cancerous Inhibitor of Protein Phosphatase 2A (CIP2A) in Th17 Cells"

**Su+A26+A1:H25+A1:H23+A26+A1:H25+A1:H27+A1:H26+A1:H25+A1:H24+A1:H26+A1:A1:H29**

The table lists the $log_2$(Relative abundance) data from the SRM analysis of the CIP2A vs. control IgG pull downs from three biological replicates for proteins validated for the CIP2A interactome.
The data integration and output made using the MSstats (3.8.4) plugin included in the Skyline software

| P.NO | UniProt Id | Protein name | Gene name | Replicate-1 | Replicate-2 | Replicate-3 |
|---|---|---|---|---|---|---|
| 1 | Q8TCG1 | CIP2A | KIAA1524 | 7,12 | 5,84 | 8,40 |
| 2 | P06239 | LCK | LCK | 6,98 | 5,05 | 8,92 |
| 4 | O95071 | UBR5 | UBR5 | 6,65 | 5,40 | 7,89 |
| 3 | P62140 | PP1B | PPP1CB | 6,85 | 4,98 | 8,71 |
| 5 | P62136 | PP1A | PPP1CA | 6,06 | 4,55 | 7,57 |
| 7 | P30153 | 2AAA | PPP2R1A | 5,07 | 3,75 | 6,39 |
| 8 | Q01196 | RUNX1 | RUNX1 | 4,98 | 3,72 | 6,24 |
| 9 | P19474 | TRIM21 | TRIM21 | 4,90 | 3,99 | 5,80 |
| 10 | P17844 | DDX5 | DDX5 | 4,80 | 3,41 | 6,19 |
| 11 | P42224 | STAT1 | STAT1 | 4,78 | 3,37 | 6,18 |
| 12 | Q8NF50 | DOCK8 | DOCK8 | 4,39 | 3,47 | 5,30 |
| 13 | Q15306 | IRF4 | IRF4 | 3,93 | 2,78 | 5,07 |
| 14 | P20592 | MX2 | MX2 | 3,77 | 2,62 | 4,92 |
| 15 | P20700 | LMNB1 | LMNB1 | 3,18 | 2,35 | 4,01 |
| 16 | P41240 | CSK | CSK | 3,18 | 2,34 | 4,02 |
| 17 | Q16666 | IF16 | IFI16 | 2,69 | 2,12 | 3,25 |
| 18 | Q00653 | NFKB2 | NFKB2 | 2,56 | 1,81 | 3,31 |
| 19 | P67775 | PP2AA | PPP2CA | 2,56 | 0,91 | 4,22 |
| 20 | P62714 | PP2AB | PPP2CB | 2,32 | 1,38 | 3,25 |
| 21 | Q13761 | RUNX3 | RUNX3 | 0,49 | 0,07 | 0,89 |

These data are available in PASSEL with the dataset identifier PASS01186
